# In search of a conditioned place preference test for mice to assess the severity of experimental procedures

**DOI:** 10.1101/2024.04.11.589117

**Authors:** A. Jaap, P. Kahnau, L. Lewejohann

## Abstract

To compare the severity of experimental procedures and behavioural tests from an animal’s perspective, novel methods are required. Theoretically, one feasible approach could be the use of a conditioned place preference test (CPP). This test employs the preference for a certain area in a test apparatus being associated with an experimental treatment. Traditionally, the CPP is used to investigate, for example, the effects of drugs. Instead we aimed to develop a protocol that would enable us to compare the effects of different experimental procedures conducted with mice.

Nine experiments with C57BL/6J mice were performed, varying the setup, the procedure duration, the stimuli as well as the presentation order. None of the tested protocols resulted in a distinct preference. Moreover, even simple protocols using food reward as a treatment failed to result in a conditioned place preference. In summary, none of the protocols was sufficient to form a reliable association between conditioned and unconditioned stimulus. We have scrutinized the experimental setup in detail, and we cannot present a solution yet. However, hopefully, our findings will help to create a working CPP to compare the severity of different experimental procedures for mice.

## 1. Introduction

This pilot study focusses, as the title states, on the search for a working protocol of a conditioned place preference test (CPP) for severity assessment. We want to emphasize beforehand that we were not successful in doing so, and think of this article as a description on what approaches have already been tried. The narration follows the experiments chronologically, explaining for each the results, the conclusions we drew, and the respective redesign we made for the next experiment. It is important to notice that our aim, severity assessment, imposed some framework conditions which we had to consider.

Severity assessment of experimental procedures conducted with laboratory mice, such as the water maze or the Barnes maze, is a complex task and to date reliable and comparable measures are still missing (Habedank et al., 2018). Usually, the severity is assessed based on physiological (e.g., weight, temperature), biochemical (e.g., corticosterone level), or behavioural measures (e.g., facial expression, alterations in typical behaviour like nest-building or wheel running) (Hohlbaum et al., 2018; Häger et al., 2018; Smith et al., 2018; Schwabe et al., 2019; Mierden, 2020). However, interpretation of these data can be challenging, especially when comparing experimental procedures and not treatments (Habedank et al., 2018). In addition, it does not take into account the perception of the situation by the animals themselves. To gain insight on the animals’ perspective, the effect of the experimental procedure could be investigated by means of preference tests (Habedank et al., 2018), including the CPP.

### 1.1. General principle of the CPP

The CPP test (or conversely the conditioned place aversion test) is commonly used to assess the effect of drugs, such as ethanol (preference) or lithium chloride (aversion) (Tzschentke, 1998, 2007; Cunningham, Gremel, & Groblewski, 2006; Wang, Wang, & Chen, 2014). The CPP has already been used to test for welfare by comparing the effect of morphine and saline on tumor-free and tumor bearing mice (Roughan et al., 2014). However, the CPP can also be used to assess other reinforcers like home cage odours (Fitchett, Barnard, & Cassaday, 2006), food (Imaizumi, Takeda, & Fushiki, 2000; Takeda et al., 2001) or male aggression (Martínez et al., 1995).

The CPP is a form of classical (Pavlovian) conditioning, in which two neutral stimuli (NS, afterwards: conditioned stimuli, CS) each become associated with a different motivationally significant stimulus (unconditioned stimuli, US). As a consequence of this learning procedure, the previously neutral CS are able to evoke a similar response to the one caused by the US. In the subsequent preference test, the animal is offered a choice between a spatial location near one of the CS. If the subject spends more time near one CS than near the other, this stimulus is interpreted as positively (or less negatively) associated than the second stimulus. Hence, with the help of a CPP, the valence of the unconditioned stimulus can be investigated.

In a next step, the CPP should be modifiable in such a way that an experimental procedure as a US is paired with a CS. Inspiration was taken from studies comparing the opportunity to run on a running wheel with no running (US) in a compartment with specific CSs (rats: Lett et al. (2001), Masaki & Nakajima (2008); hamsters: Antoniadis et al. (2000)): By offering the choice between two CS which represent both two different experimental procedures, a comparison between the effects of the two experimental procedures could be made. Hence, with the help of the CPP, severity assessment or at least severity comparison of experimental procedures should be possible. In this manner, for example the Morris Water Maze and the Barnes Maze, which both focus on spatial learning but differ with regard to the perceived severity from a human perspective, could be compared in regard of their severity from the mice’s perspective. Here, we aimed to develop such a protocol for CPPs to compare the severity of experimental procedures. However, we wanted to start small, using simpler experimental procedures with more defined, known valence, for example restraint (as e negative stimulus) or food reward (as a positive stimulus). Only if this protocol was successfully established, we wanted to use it to test, for example, the effect of the Morris Water Maze.

### 1.2. Various ways of CPP conduction

The main question was: How to best perform a CPP to compare experimental procedures? Because there are various ways to perform CPPs, even in drug testing. As an example for the great variety in CPP conductions, we have listed some studies and their differences in Table 1. As there is no existing report of the development of such CPP protocols, it is often unclear in which cases one is free to choose between multiple ways of conduction or in which cases only a specific combination works. In the following, we want to give a short overview how differently CPPs can be conducted because this also influenced our decisions (as explained later in the methods and results).

**Table 1:**
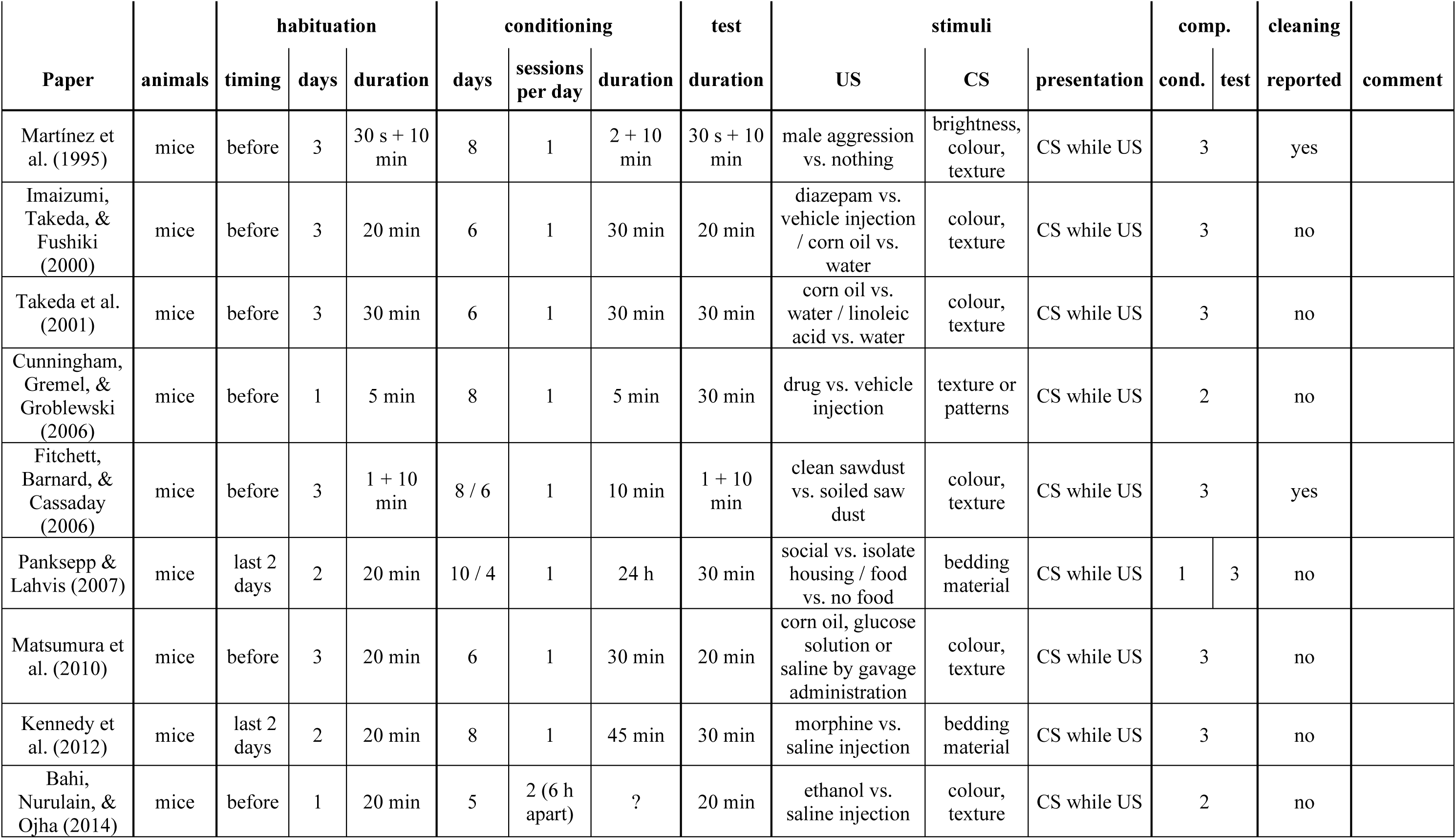
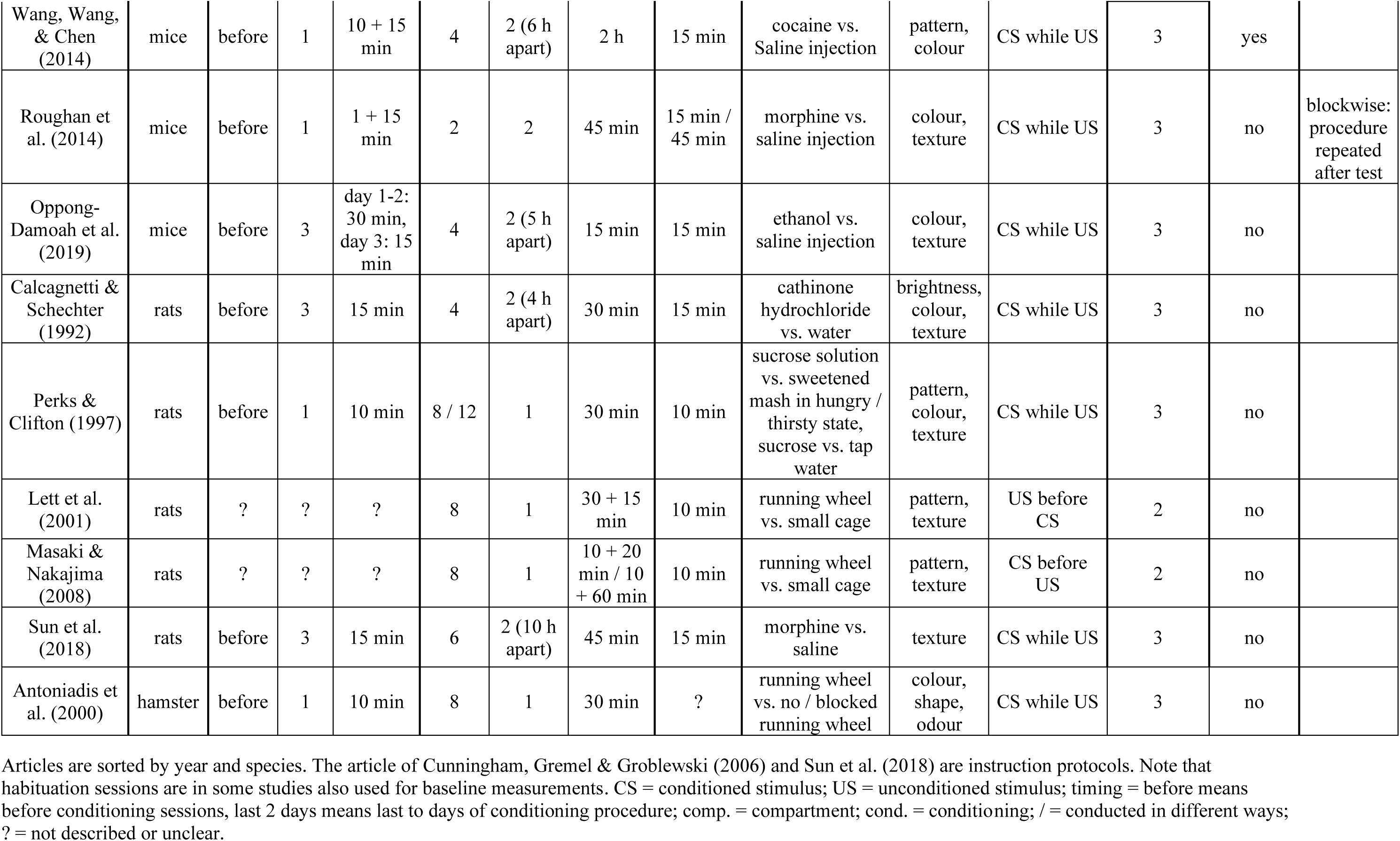
Examples of CPP experiments and their differing implementations.

In general, usually four or more conditioning sessions per treatment are conducted (i.e., pairing NS and US). Usually, this is done with one session per day (e.g., protocol by Cunningham, Gremel, & Groblewski, 2006). However, it is also possible, to perform two sessions per day, and thus, shortening the protocol (rats: Calcagnetti & Schechter (1992); Sun et al. (2018); mice: Bahi, Nurulain, & Ojha (2014); Wang, Wang, & Chen (2014); Oppong-Damoah et al. (2019)).

When planning a CPP, the choice of the CS is important, meaning whether to use visual (Wang, Wang, & Chen, 2014), tactile (Sun et al., 2018), olfactory cues (Antoniadis et al., 2000), or a combination of those (Cunningham, Patel, & Milner, 2006; Cunningham & Zerizef, 2014; Cunningham & Shields, 2018). This choice should not be underestimated as some stimuli are easier associated with specific reinforcers (US) than others (Garcia & Koelling, 1966).

Differences in the conduction of CPPs can also be found in the conditioning setup, in which the CS are presented: It is possible to use one compartment with exchanged stimuli for the conditioning sessions and a two or three compartment setup for the preference test (Panksepp & Lahvis, 2007). However, it is more common, to use a two-(Cunningham, Gremel, & Groblewski, 2006) or three-compartment (Sun et al., 2018) setup for both testing and conditioning, with the access to the second compartment blocked during the conditioning sessions. The advantage of this latter setup is that the spatial position of the compartments can also be used as a CS if translucent walls are used (Cunningham, Patel, & Milner, 2006; Cunningham & Zerizef, 2014).

An additional point which has to be considered for conditioning is the timing of stimulus presentation. When testing the effect of drugs, for which the CPP is most commonly used, US (drug) and CS (e.g., pattern) can occur at the same time. Thus, the scheduled time spent in the conditioning compartment in proximity to the CS depends on the time duration between drug administration and the onset of its effect (e.g., 5 min in the conditioning setup protocol for drugs in general: Cunningham, Gremel & Groblewski (2006), 2 h for cocaine: Wang, Wang, & Chen (2014), 30 min for access to corn oil or water: Takeda et al. (2001)). However, if experimental procedures are to be tested, we have to choose: present the US and CS simultaneously, present the US before the CS or present the CS before the US. This decision is important as timing might result in different effects: Presenting the pattern before the access to the running wheel (rats: Masaki & Nakajima (2008)) produced place *aversion*, while presenting the pattern simultaneously (hamsters: Antoniadis et al. (2000)) or afterwards (rats: Lett et al. (2000, 2001) Lett, Grant & Koh (2002)) produced place *preference*. This effect of presentation time is also known for drugs like amphetamine (Fudala & Iwamoto, 1990) and nicotine (Fudala & Iwamoto, 1986, 1987). Also, the duration of presentation of the US and the CS might depend on the tested US. In some cases, the simultaneous presentation of US and CS can last 24 h, as was done to compare social and isolated housing (Panksepp & Lahvis, 2007).

There are also great differences in timing (duration and time point) of the habituation to the setup: Some studies conducted the habituation before the conditioning sessions, ranging from 5 min on 1 day (Cunningham, Gremel, & Groblewski, 2006) to 10 min on 3 days (Martínez et al., 1995). Other studies performed it on 2 days at the last 2 days of conditioning, meaning before the final test, not before the conditioning sessions (for 20 min: (Panksepp & Lahvis, 2007; Kennedy et al., 2012)). When performing the habituation before the beginning of the first conditioning session, it can also be used as a baseline preference test, to compare the preference for the two CS before the conditioning sessions. If the habituation session is conducted at the end of the conditioning phase, habituation can only be conducted with a setup emptied from all stimuli (to not disturb the conditioning), and thus, baseline measurements are not possible.

Comparing different test durations (i.e., the final test), the range between studies is smaller: There are studies, which measured the stay preference within 10 min (rats: (Perks & Clifton, 1997), mice: Martínez et al. (1995)), and other in 45 min (mice: Roughan et al. (2014)). Thus, the duration of the final test is often chosen independently from the duration of the conditioning sessions.

Interestingly, not all studies report the time of day when conditioning or tests were conducted (e.g., no detailed instructions given in Cunningham, Gremel & Groblewski (2006), they just describe that it is performed during the light phase). However, this might have an influence, for example, on the motivation of the mice to gain food (Acosta et al., 2020; Koch et al., 2020).

### 1.3. Aim

In this present study, we aimed to establish a protocol for a CPP test which would allow a comparison of two experimental procedures and their effects (US) by pairing the respective procedure with different CS. We focused on the CPP protocol by Cunningham and colleagues (Cunningham, Gremel, & Groblewski, 2006) but due to our experimental question, we had to make some adjustments. For the general design of our CPP, we had several aims:

First, we wanted to develop a protocol suitable for mice, more precisely for C57BL/6J mice. This strain is the most commonly used in laboratory experiments; therefore, finding a working protocol to conduct CPP tests with experimental procedures would make the greatest impact for severity assessment.

Second, the CPP protocol for assessing severity must not be in any kind severe in itself. Otherwise it could cause misinterpretations of the results, and add to existing severity of the to be tested experimental procedures. This also led to the decision that for this CPP protocol, food or water deprivation (as used by Masaki & Nakajima (2008) and Lett et al. (2001) in their running wheel experiments) would not be feasible.

Third, in the optimal case, the CPP procedure should be able to detect not only large differences in severity of the tested experimental procedures but also subtle ones. For this reason, the experimental procedures which were used here as US are only mildly severe, and not moderate or severe (according to EU classification in annex VIII of the Directive 2010/63/EU). For example, the time in the restrainer was restricted to 1 min instead of the 15 min (Glavin et al., 1994) or even 6 h (Nievas et al., 2010) used in other studies.

Here, we report all of our attempts to establish such a CPP protocol for the comparison of experimental procedures. We want to emphasise that the experiments described were all preliminary tests which did not lead to an overall functioning protocol. However, we find it important to report them as an assistance for future research in this direction.

## 2. Materials and Methods

### 2.1. Overview

Altogether, we conducted twelve experiments, nine of them based on a conditioned place preference design. Three additional experiments (not CPP), which we conducted to test a different approach for the comparison of the severity, are described in the Supplements.

The experiments we conducted here were preliminary and served to develop a protocol to use CPP for severity assessment. In other words: In contrast to other experiments, we did not want to know the effect of a specific US (if it is aversive or attractive), but had to choose an US with a predictable effect to validate our protocol. For example, we used restraint or fixation as a negative US, weighing as a neutral US, or a food reward (millet) as a positive US. In this manner, we investigated whether the experiment protocol itself worked. Afterwards we wanted to move on to comparisons of procedures with uncertain outcomes.

However, none of the experiments led to the anticipated results and conditioning (at least not in the desired way). In search of a working protocol, we did not conceptualise a set of experiments beforehand but adjusted our procedures with each independent, new experiment, analysed the new result and again, returned to adjusting the procedure. In Table 2, the nine CPP experiments are summed up to give a better overview.

**Table 2:** Summary of the independent CPP procedures used in experiment 1 to 9.

| No. | Gr. | Age | Duration | US1 | US2 | NS → CS | CS Presentation | Setup | Room |
| --- | --- | --- | --- | --- | --- | --- | --- | --- | --- |
| 1 | 1 | 15 | 10 days | fixation | weighing | bedding | before + after | 1 | other |
| 2 | 1 | 17 | 6 days | fixation | weighing | gravel | before | 1 | other |
| 3 | 1 | 19 | 6 days | fixation | weighing | plates | before | 2 | other |
| 4 | 2 | 2 | 6 days | millet | weighing | plates | before | 2 | other |
| 5 | 2 | 9 | 6 days | almond milk | water | patterns | sim | 2 | other |
| 6 | 2 | 12 | 10 days | millet + bedding | bedding | patterns | sim | 2 | same |
| 7 | 3 | 13 | 10 days | millet | bedding | patterns | sim (30 s delay) | 3 | same |
| 8 | 3 | 14 | 10 days | restrainer | millet | patterns | after | 3 | same |

| No. | Gr. | Age | Duration | US1 | US2 | NS → CS | CS Presentation | Setup | Room |
| --- | --- | --- | --- | --- | --- | --- | --- | --- | --- |
| 9 | 4 | 10 | 10 days | restrainer | millet | patterns + plates | after | 3 | same |

No. = Number of experiment, Gr. = Group, Age = age in months of the mice, US = unconditioned stimulus, NS = neutral stimulus which becomes the CS = conditioned stimuli, sim = simultaneously, same = experiment was conducted in the husbandry room, other = experiment was conducted in a separate, experimental room than the usual husbandry room. In the results section, all drawn conclusions and the consequent sometimes multi-factorial changes for the next experiment are explained in detail.

In the following section, we will first go into details about general methods which were similar throughout all the experiments. A more detailed description of each experiment will then be given in section Experiments and Results, in which we provided always a short description of the experiment (and a reasoning why we decided upon this conduction) and directly afterwards the results of this experiment. We decided upon this structure because the alterations in the procedures were directly related to the results from the previous experiment, and we believe in this manner, our considerations will become more clear.

In addition, at the end of the article and in the Supplements, detailed information on the neutral, to be the conditioned stimuli (NS → CS, in the following called “conditioned stimulus” or CS) as well as the conduction of the procedures used as unconditioned stimuli (US) is provided. They are meant as a glossary, giving more detailed information when needed without interrupting the main description of the experiments.

### 2.2. Animals

#### 2.2.1. Groups

Throughout the course of the experiments, four groups of mice were enrolled. They each took part in different experiments:

##### All mice

All mice were female C57BL/6J CrL mice and purchased from Charles River (Sulzfeld, Germany). All mice within each group had different mothers and different foster-mothers to ensure maximal epigenetic variability within the inbred strain and avoid maternal effects. With the arrival at our institute, the mice within one group were housed together in one cage system. At the age of five weeks, transponders were implanted, a procedure performed under anaesthesia and analgesia (for details see section Transponder implantation). All groups were *always* handled by tunnel handling (for more details see https://wiki.norecopa.no/index.php/Mouse_handling, as well as Hurst & West (2010); Gouveia & Hurst (2017)).

##### Group 1

A group of thirteen mice was purchased in December 2017 at the age of 3 weeks. This group took part in experiments 1-3 (12 months of age at the start of experiment 1). This group participated beforehand in other experiments, e.g., a T-maze preference test (Habedank, Kahnau, & Lewejohann, 2021) as well as development and first tests of an RFID based tracking system for home cage based choice tests (Habedank et al., 2022).

##### Group 2

The second group consisting of twelve mice was purchased in June 2019 at the age of 4 weeks. This group took part in experiments 4-6 (2 months of age at the start of experiment 4). This group participated beforehand in other experiments, e.g., the validation of an RFID based tracking system for home cage based choice tests (Habedank et al., 2022) and in-between the experiments described here, they were also part of a T-maze test (Habedank, Kahnau, & Lewejohann, 2021).

##### Group 3

The third group consisted of twelve mice and was purchased in September 2019 at the age of 4 weeks. This group took part in experiments 7-8 (13 months at the start of experiment 7). This group participated beforehand in other experiments, e.g., the development of a home cage based cognitive bias test (Kahnau, Jaap, et al., 2023).

##### Group 4

The fourth group consisted of twelve female C57BL/6J CrL mice was purchased in September 2020 at the age of 4 weeks. This group took part in experiment 9 (10 months at the start of it). This group participated also beforehand in other experiments, e.g., the development of a home cage based cognitive bias test (Kahnau, Jaap, et al., 2023).

##### Additional remarks

It was noted a few weeks before experiment 5 that eleven of the twelve mice of group 2 completely or partly lacked their whiskers, probably due to plucking / barbering behaviour. The same was true for all twelve mice of group 3 in experiments 7 and 8. As explained in more detail in the discussion (section Whisker-loss), we decided against ordering a new group of mice, especially because we were conducting preliminary tests for a proof of concept. Instead we adjusted the CS, focusing on visual instead of tactile cues. In addition, we repeated experiment 8 with a fully whiskered group in experiment 9.

Please also note that we enrolled mice up to a high age for the same reasons. The arguments why we consider this as feasible are also given in the discussion (section Age). Moreover, we took care that the tests in which the animals participated beforehand were of a different nature, and thus, no after-effect of these was to be anticipated.

#### 2.2.2. Housing

Groups of mice were kept in two connected type IV macrolon cages (L x W x H: 598 x 380 x 200 mm, Tecniplast, Italy) with filter tops. As connection between the cages, a Perspex tunnel (40 mm in diameter) was used. The mice had been living in this system since they were around 2 months (group 1), 3 months (group 2) or 1 month (group 3 and 4). In group 3 and 4, the tunnel had been replaced by an AnimalGate (TSE systems, Germany) and the connected cage was an IntelliCage (TSE systems, Germany) during the months preceding the CPP.

Food (autoclaved pellet diet, LAS QCDiet, Rod 16, LASvendi, Germany) and tap water (two bottles each cage) were available *ad libitum*. Both cages were equipped with bedding material (Poplar Granulate 2-3 mm, Altromin, Germany) of 3-4 cm height, a red house (The MouseHouse, Tecniplast), nesting material (papers, cotton rolls, strands of paper nesting material), and two wooden bars to chew on. Both cages also contained a Perspex tunnel (40 mm in diameter, 17 cm long), which was used for tunnel handling. Group 3 and 4 also experienced weekly changing cage equipment (houses, nesting material, active enrichment filled with millet; this was introduced after the findings of Hobbiesiefken et al. (2021)).

Room temperature was maintained at 22 ± 3 ^∘^C, the humidity at 55 ± 15 %. Animals were kept at 12h/12h dark/light cycle with the light phase starting at 7:00 a.m. (winter time) or 8:00 a.m. (summer time), respectively. Between 6:30 and 7:00 a.m. (winter time) or 7:30 and 8:00 (summer time) a sunrise was simulated: A wake-up light (HF3510, Philips, Germany) in one corner of the room gradually increased the light intensity until the overhead light went on.

Animals were visually inspected in their home cages daily between 7:00 a.m. (winter time) and 8:00 a.m. (summer time) to 10 a.m. Once per week, home cages were cleaned and all mice were scored and weighed. In this context, mice also received a colour code on the base of their tails, using edding 750 paint markers, to facilitate individual recognition.

#### 2.2.3. Experimental Room

In experiments 1 to 5, we used a separate room for the experimental procedures and not the room in which the animals were kept in. Between experiments 4 and 5 (before 4.1, not a CPP experiment, see Supplements) the housing room was changed, and for logistical reasons, the experimental room had also to be changed. For both used experimental rooms, temperature was maintained at 22 ± 3 ^∘^C and humidity at 55 ± 15 %. The rooms had an automatic 12h/12h dark/light cycle with the light phase starting at 7:00 a.m. (winter time) or 8:00 a.m. (summer time), respectively.

#### 2.2.4. Transponder implantation

At the age of five weeks, transponders (FDX-B transponder according to ISO 11784/85; group 1: Planet-ID, Germany; group 2 – 4: Euro I.D., Germany) were implanted under the skin in the neck of the mice. To do so, in group 1 all mice obtained an analgesic (Meloxicam, 1mg/kg) two hours before the procedure. The transponder implantation itself was performed under isoflurane anesthesia (induction of anesthesia: 4 l/min 4 %; maintenance of anesthesia: 1 l/min 1-2 %). RFID (radio frequency identification) transponders were injected directly behind the ears subcutaneously in the neck, so that they were rostrocaudal oriented (for detailed description see Habedank et al. (2022)). After transponder implantation, mice were placed individually in a separate cage with bedding and sheets of paper, and monitored until they were fully awake again. Then they were returned to their home cage. In group 1, two mice lost their transponders after the first implantation, and for those two mice the transponder implantation was repeated at the age of 8 weeks.

For group 2 – 4, the administration time of the analgesic was altered to the evening before the procedure. We hoped to reduce transponder loss this way: By administering the Meloxicam earlier, the analgesic effect was expected to cease before the dark phase (active phase) after the implantation, and mice would be more hesitant to focus on the injection side. Implantation of the transponders was performed in the same way as in group 1. In group 2, no transponder was lost. In group 3 and 4, one transponder was lost and the procedure was repeated with the respective mouse the very next day (group 3) or five days later (group 4), respectively.

### 2.3. Setups

In total, three setups were used. For reasons of comprehensibility, details on the used setups will be explained in the specific experiments in which they were first put to use.

### 2.4. Conditioned and Unconditioned Stimuli

The used conditioned and unconditioned stimuli are mentioned in short in the description of the corresponding experiments. For more details on the stimuli, we provide a detailed description at the end of the article.

### 2.5. General Procedure

For our research question, we focused on the CPP protocol by Cunningham and colleagues (Cunningham, Gremel, & Groblewski, 2006). However, the details of the conduction differed throughout the experiments (as explained later in section Experiments and Results). However, for all of the following experiments, some parameters always remained the same:

#### Randomization

The pairing of neutral, to be conditioned stimulus (NS → CS) and unconditioned stimulus (US) was randomized for all mice (e.g., for half of the mice CS A was paired with US 1 and for half of the mice CS A was paired with US 2). Also, the mice were randomly assigned to start with US A or US B. Within these subsets, the presentation side of the CS was randomized in such a way that about half of the mice experienced CS A left and CS B right and the other half the other way around. In addition, the order in which the mice were taken out of the home cage for the sessions was randomized, and this was also the case for the order of mice in the baseline and the preference test. Moreover, the start compartment into which the mice were placed at the beginning of the baseline and final preference test (left or right) was randomized.

#### Blinding

Baseline and final preference tests were video recorded. Analysis of the video recordings were evaluated without knowledge of the pairing of treatment (procedure, US) and CS or side for the individual animals.

#### Exclusion

Animals, which did not change compartments even once during baseline or final preference test, were excluded from analysis. This was only relevant for setup 1. Here, in both experiments, one mouse (the same in both experiments) did not change cages even once during the 10 min. Therefore, we could not tell with certainty that the mouse was aware of both options, and excluded it from the analysis. In addition, in experiment 2, the baseline test was conducted with thirteen mice. However, one mouse had to be killed before the start of the conditioning sessions due to health issues independent from the experiment. Thus, the experiment itself was only performed with twelve mice.

#### Time

Experimental procedures were conducted close to the onset of the light phase, and therefore, close to the active phase of the mice. Experiment 1 started 1 h after the onset of light (here: 7:00 a.m.) with transportation and habituation. All other experiments started right after the onset of light (7:00 or 8:00 a.m., respectively) with transportation (where necessary) and habituation. Depending on the procedure, all experiments lasted then three to five hours, depending on schedule and if there was transportation.

#### Recording

All videos were recorded using either a webcam (Logitech C930e, Switzerland) or a Basler camera (Basler ace acA1920-40um, Basler AG, Germany with a prime lens LM8HC, 8 mm, F1,4-16, Kowa, Optimed Deutschland GmbH, Germany). We used iSpy 64 (version 7.0.3.0) for recording.

#### Conduction

All experiments were conducted by the same person (AJ), so the influence by different experimenters could be excluded.

#### Schedule

We used either a 10 day or a 6 day schedule. If the experiment lasted 10 days (based on the protocol by Cunningham, Gremel & Groblewski (2006)), there was 1 day for baseline testing, followed by 8 days of conditioning and one day for the final preference test. There was one conditioning session per day, with a break after session 4 of 2 days (weekend). Conditioning sessions were alternated, i.e., a mouse, which experienced US 1 in session 1, experienced it again in session 3, 5 and 7, while US 2 was presented in sessions 2, 4, 6 and 8.

If the experiment lasted 6 days, the first day for baseline testing was followed by 4 days of conditioning and one day for the final preference test. There was a break of a few days between the baseline test and the first conditioning session to evaluate the baseline for a bias in preference. Then conditioning was performed by conducting two sessions per day. Here, in contrast to other studies using the 6 day schedule (rats: Calcagnetti & Schechter (1992), mice: Wang, Wang, & Chen (2014); Oppong-Damoah et al. (2019)), we decided against a morning and an afternoon conditioning session because this would have resulted in waking the mice in the middle of their inactive phase, and we did not want to disrupt their circadian rhythm more than necessary. Instead, we performed the conditioning sessions directly after each other. In addition, to prevent a time effect (early morning versus late morning), the procedure tested last on one day was the first to be tested the next day (e.g. conditioning day 1: US 1, US 2, conditioning day 2: US 2, US 1, conditioning day 3: US 1, US 2, and so on).

### 2.6. Cleaning

For reasons of clarity and comprehensibility, we report cleaning procedures of the setups, US and CS to the Supplements.

### 2.7. Transportation

For those experiments, which were conducted in a experimental room instead of the housing room, transportation was necessary. In experiment 1 – 5, transportation cages were used for this procedure because transportation of the home cage setup was not possible. Mice were placed into a transportation cage (type IV macrolon cage, LWH: 598 x 380 x 200 mm, Tecniplast). The transportation cage was equipped with the usual bedding material, a red house (The MouseHouse, Tecniplast) and two sheets of paper as nesting material/enrichment. Food (pellet diet, LAS QCDiet, Rod 16, autoclavable, LASvendi) and water (tap water, two bottles) were provided *ad libitum*.

### 2.8. Video Analysis

During the baseline and the preference test, the animals were placed for 10 min into the setup. From this total duration (starting at the moment, the filter top or plate was placed on top of the setup or, in experiment 6 – 9, at the moment the mouse left the handling tunnel with all four feet), the first minute was taken as habituation time and not analysed any further. The remaining nine minutes were then analysed. Additional details on video analysis are given in context with the setup descriptions.

### 2.9. Statistical Analysis

Data was analysed with regard to side preference (left vs. right compartment), CS presented in the compartments (e.g., horizontal vs. vertical stripes) and US paired with the presented CS (e.g., millet vs. bedding). This was also done for the baseline test. In this case, the test was done for a compartment with which a specific procedure was going to be paired (“procedure compartment”), although the mice hadn’t experienced the combination of compartment and procedure yet. In general, the amount of time in each compartment was calculated as percentage, either of the complete time after 1 min habituation (9 min, setup 1 and 2) or of the total time in one of the compartments (9 min minus the time spent between compartments, setup 3).

All statistics and diagrams were calculated using R (version 4.0.3 to 4.1.0) and RStudio (version 1.4.1717-3, Posit PBC, Boston, USA). To test for normal distribution, the Shapiro–Wilk test was performed. In all experiments, the data did not differ significantly from a normal distribution (p > 0.05); therefore, a t-test was used to compare the preferences with a random chance level of 0.5. For comparison between pre-conditioning and post-conditioning results, a paired t-test was used. In all statistical tests, significance level was set to 0.05 and it was tested two-sided. Due to the exploratory approach of the study and because there were no significant differences to be corrected, the results are not Bonferroni corrected. Results in the text are always reported as the mean ± standard deviation. Plots are created using the ggplot2 package in R with the function geom_boxplot() which always shows the median (bar), first and third quartiles (lower and upper hinges), the largest value with additional 1.5 times inter-quartile ranges (whiskers), and all outliers beyond that (points).

## 3. Experiments and Results

In this section, we will give a short description of the procedure used in each experiment and directly provide the results.

Note that in this main article, we will present the results of the video analyses (the baseline and the final test). An overview of the results, i.e., mean, standard deviation, p-values for the tested factors the CS (cue), US (procedure), and the side, are given in Table 3, Table 4 and Table 5. In addition, in each section corresponding to the experiments, the results are depicted graphically as box-plot diagrams. Additional observations, e.g., on the behaviour of the mice during the procedure of the experiments, are described in the Supplements.

**Table 3:** Overview of the statistic results regarding the Conditioned Stimulus of the CPP experiments.

| exp. | n | comparison | what | average | statistics |
| --- | --- | --- | --- | --- | --- |
| 1 | 12 | baseline | cw vs. pure bedding | cw: $59.95 \pm 7.54$ % | $p = 0.0008034$, $t = -4.57$ |
| | | test | | cw: $64.88 \pm 9.52$ % | $p = 0.0002115$, $t = -5.4159$ |
| | | pre vs. after | | | $p = 0.216$, $t = 1.3127$ |
| 2 | 11 | baseline | quartz vs. marble gravel | q: $53.08 \pm 10.95$ % | $p = 0.3731$, $t = 0.93244$ |
| | | test | | q: $52.44 \pm 19.57$ % | $p = 0.6879$, $t = -0.41364$ |
| | | pre vs. after | | | $p = 0.4098$, $t = 0.86018$ |
| 3 | 12 | baseline | holes vs. slit plate | s: $50.19 \pm 4.63$ % | $p = 0.8923$, $t = -0.13859$ |
| | | test | | h: $54.26 \pm 12.70$ % | $p = 0.2701$, $t = 1.1614$ |
| | | pre vs. after | | | $p = 0.2405$, $t = -1.2407$ |
| 4 | 12 | baseline | holes vs. slit plate | s: $50.31 \pm 9.97$ % | $p = 0.9165$, $t = 0.10728$ |
| | | test | | s: $54.07 \pm 7.02$ % | $p = 0.0696$, $t = 2.01$ |
| | | pre vs. after | | | $p = 0.1382$, $t = -1.5988$ |
| 5 | 12 | baseline | dots vs. stripes | s: $53.64 \pm 8.39$ % | $p = 0.1611$, $t = -1.5026$ |
| | | test | | d: $52.32 \pm 8.04$ % | $p = 0.3393$, $t = 0.99901$ |
| | | pre vs. after | | | $p = 0.008077$, $t = -3.2257$ |
| 6 | 12 | baseline | cb vs. f pattern | f: $50.73 \pm 6.18$ % | $p = 0.6916$, $t = -0.40735$ |
| | | test | | f: $46.63 \pm 10.47$ % | $p = 0.2883$, $t = 1.1157$ |
| | | pre vs. after | | | $p = 0.3088$, $t = -1.0671$ |
| 7 | 12 | baseline | horizontal vs. vertical stripes | h: $52.18 \pm 10.17$ % | $p = 0.4737$, $t = -0.7419$ |
| | | test | | h: $50.10 \pm 9.95$ % | $p = 0.9718$, $t = -0.036123$ |
| | | pre vs. after | | | $p = 0.4464$, $t = -0.78976$ |
| 8 | 12 | baseline | horizontal vs. vertical stripes | h: $53.69 \pm 8.76$ % | $p = 0.1732$, $t = -1.4567$ |
| | | test | | v: $53.49 \pm 11.12$ % | $p = 0.3005$, $t = 1.0865$ |
| | | pre vs. after | | | $p = 0.009003$, $t = -3.1647$ |
| 9 | 12 | baseline | horizontal vs. vertical stripes | v: $50.42 \pm 12.51$ % | $p = 0.909$, $t = 0.11692$ |
| | | test | | h: $51.08 \pm 9.90$ % | $p = 0.7128$, $t = -0.37781$ |
| | | pre vs. after | | | $p = 0.6252$, $t = 0.50256$ |

All results were normally distributed, so either a t-test (baseline vs. chance and test vs. chance) or a paired t-test (baseline vs. test) were conducted. The number of animals is the number of animals after exclusion. All percentages are given as mean and standard deviation. exp. = experiment, n = number of animals, baseline = baseline vs. chance, test = test vs. chance, pre vs. after = pre-conditioning (baseline) vs. after conditioning (test), cw = comfort white bedding, cb = chessboard pattern, f = fabric-like pattern, abbreviations in the column “average” correspond to the stimuli in column “what”.

**Table 4:** Overview of the statistic results regarding the Unconditioned Stimulus of the CPP experiments.

| exp. | n | comparison | what | average | statistics |
| --- | --- | --- | --- | --- | --- |
| 1 | 12 | baseline | fixation vs. weighing | w: $51.18 \pm 12.79 \%$ | p = 0.7543, t = -0.3209 |
| | | test | | f: $52.27 \pm 18.07 \%$ | p = 0.6719, t = 0.4351 |
|  |  | pre vs. after |  |  | p = 0.3945, t = -0.8861 |
| 2 | 11 | baseline | fixation vs. weighing | w: $56.98 \pm 8.77 \%$ | p = 0.02476, t = -2.6394 |
| | | test | | w: $53.04 \pm 19.48 \%$ | p = 0.6164, t = -0.51702 |
|  |  | pre vs. after |  |  | p = 0.5599, t = -0.60308 |
| 3 | 12 | baseline | fixation vs. weighing | w: $50.83 \pm 4.55 \%$ | p = 0.5388, t = -0.63443 |
| | | test | | f: $50.77 \pm 13.44 \%$ | p = 0.8459, t = -0.19893 |
|  |  | pre vs. after |  |  | p = 0.6804, t = -0.42303 |
| 4 | 12 | baseline | millet vs. weighing | m: $53.86 \pm 9.12 \%$ | p = 0.1708, t = -1.4652 |
| | | test | | m: $51.27 \pm 8.10 \%$ | p = 0.5993, t = 0.54097 |
|  |  | pre vs. after |  |  | p = 0.321, t = 1.0391 |
| 5 | 12 | baseline | almond milk vs. water | am: $50.74 \pm 9.18 \%$ | p = 0.7855, t = 0.27892 |
| | | test | | am: $50.86 \pm 8.34 \%$ | p = 0.7292, t = 0.35517 |
|  |  | pre vs. after |  |  | p = 0.5415, t = -0.63009 |
| 6 | 12 | baseline | millet + bedding vs. bedding | b: $52.15 \pm 5.81 \%$ | p = 0.2255, t = -1.284 |
| | | test | | m+b: $52.42 \pm 10.75 \%$ | p = 0.4513, t = 0.78089 |
|  |  | pre vs. after |  |  | p = 0.2527, t = -1.2073 |
| 7 | 12 | baseline | millet vs. bedding | m: $51.02 \pm 10.37 \%$ | p = 0.7386, t = 0.34226 |
| | | test | | m: $54.83 \pm 8.57 \%$ | p = 0.07697, t = 1.9511 |
|  |  | pre vs. after |  |  | p = 0.148, t = -1.5561 |
| 8 | 12 | baseline | millet vs. restrainer | r: $54.17 \pm 8.52 \%$ | p = 0.118, t = -1.6958 |
| | | test | | r: $54.42 \pm 10.76 \%$ | p = 0.1818, t = -1.4255 |
|  |  | pre vs. after |  |  | p = 0.9367, t = 0.081281 |
| 9 | 12 | baseline | millet vs. restrainer | r: $55.36 \pm 11.19 \%$ | p = 0.1247, t = -1.6622 |
| | | test | | r: $54.63 \pm 8.70 \%$ | p = 0.09202, t = -1.8456 |
|  |  | pre vs. after |  |  | p = 0.8125, t = -0.24293 |

All results were normally distributed, so either a t-test (baseline vs. chance and test vs. chance) or a paired t-test (baseline vs. test) were conducted. The number of animals is the number of animals after exclusion. All percentages are given as mean and standard deviation. exp. = experiment, n = number of animals, baseline = baseline vs. chance, test = test vs. chance, pre vs. after = pre-conditioning (baseline) vs. after conditioning (test), abbreviations in the column “average” correspond to the stimuli in column “what”.

**Table 5:** *Overview of the statistic results regarding the compartments of the CPP experiments*.

| exp. | n | comparison | average | statistics |
| --- | --- | --- | --- | --- |
| 1 | 12 | baseline | l: $52.58 \pm 12.56 \%$ | p = 0.4911, t = 0.71238 |
| | | test | l: $53.55 \pm 17.84 \%$ | p = 0.5044, t = 0.69015 |
|  |  | pre vs. after |  | p = 0.8137, t = -0.24142 |
| 2 | 11 | baseline | l: $53.04 \pm 10.96 \%$ | p = 0.3787, t = 0.921 |
| | | test | l: $52.93 \pm 19.49 \%$ | p = 0.6287, t = 0.49887 |
|  |  | pre vs. after |  | p = 0.9869, t = 0.016807 |
| 3 | 12 | baseline | l: $50.09 \pm 4.63 \%$ | p = 0.946, t = 0.069252 |
| | | test | r: $50.19 \pm 13.46 \%$ | p = 0.9628, t = 0.047662 |
|  |  | pre vs. after |  | p = 0.9811, t = -0.02421 |
| 4 | 12 | baseline | r: $53.70 \pm 9.19 \%$ | p = 0.1903, t = -1.396 |
| | | test | l: $50.64 \pm 8.18 \%$ | p = 0.7889, t = 0.2744 |
|  |  | pre vs. after |  | p = 0.08055, t = -1.9244 |
| 5 | 12 | baseline | r: $50.41 \pm 9.20 \%$ | p = 0.8795, t = -0.15523 |
| | | test | r: $53.63 \pm 7.49 \%$ | p = 0.1209, t = -1.681 |
|  |  | pre vs. after |  | p = 0.2042, t = -1.3497 |
| 6 | 12 | baseline | l: $52.43 \pm 5.68 \%$ | p = 0.166, t = 1.4834 |
| | | test | l: $46.08 \pm 10.26 \%$ | p = 0.2127, t = -1.3229 |
|  |  | pre vs. after |  | p = 0.1011, t = 1.7891 |
| 7 | 12 | baseline | l: $56.54 \pm 7.87 \%$ | p = 0.01503, t = 2.8779 |
| | | test | l: $56.38 \pm 7.38 \%$ | p = 0.01224, t = 2.9925 |
|  |  | pre vs. after |  | p = 0.9536, t = 0.059569 |
| 8 | 12 | baseline | l: $52.60 \pm 9.18 \%$ | p = 0.3477, t = 0.98089 |
| | | test | l: $53.53 \pm 11.11 \%$ | p = 0.2947, t = 1.1003 |
|  |  | pre vs. after |  | p = 0.7711, t = -0.29821 |
| 9 | 12 | baseline | l: $56.65 \pm 10.42 \%$ | p = 0.04917, t = 2.2106 |
| | | test | l: $56.82 \pm 6.97 \%$ | p = 0.006033, t = 3.3902 |
|  |  | pre vs. after |  | p = 0.956, t = -0.056474 |

All results were normally distributed, so either a t-test (baseline vs. chance and test vs. chance) or a paired t-test (baseline vs. test) were conducted. The number of animals is the number of animals after exclusion. All percentages are given as mean and standard deviation. exp. = experiment, n = number of animals, baseline = baseline vs. chance, test = test vs. chance, pre vs. after = pre-conditioning (baseline) vs. after conditioning (test), l = left, r = right.

### 3.1. Experiment 1

#### 3.1.1. Starting Considerations

For the first CPP, we started with simple experimental procedures for which we could predict the effect: weighing as a supposedly neutral procedure and fixation (restraint by hand) as a supposedly negative procedure. The schedule was based on the protocol by Cunningham, Gremel & Groblewski (2006) and we used a conditioning setup which was simple and resembled a smaller version of their home cage (two connected cages). As CS we used two types of bedding material, which differed in visual, olfactory and tactile cues, arguing that these would be natural, but distinct stimuli.

#### 3.1.2. Setup 1: Two cages connected via a tunnel

Setup 1 was used for the baseline test and the preference test in experiment 1 and 2. Two type II cages (LWH: 225 x 167 x 140 mm, Tecniplast) were connected with a tunnel (4 cm in diameter, 24 cm long). Both cages were closed by a lid and a filter top. Between the cages, an automated positioning system based on light barriers was installed (see Figure 1): The light barriers were 15 cm apart and connected to an Arduino Leonardo micro controller with a real-time clock, an SD card and a time display. Whenever changing cages and passing through the tunnel, a mouse would interrupt the two light barriers. A self-written software ensured that each interruption of the light barriers was saved with a time stamp onto the SD card. If the left light barrier was interrupted last, the mouse was counted as belonging to the left cage, and if the right light barrier was interrupted last, the mouse belonged to the right cage, respectively. As the positioning system did not register the start time (i.e., when the lid was closed), we used video recordings to get this time point. To verify the automatic positioning method, for the habituation session the automated results were compared to video recordings. The positioning system used here was a prototype of the system which later on became the “MoPSS”: The Mouse Positioning Surveillance System is an open-source RFID based tracking system suitable for individual tracking of group housed mice (Habedank et al., 2022).

**Figure 1:**
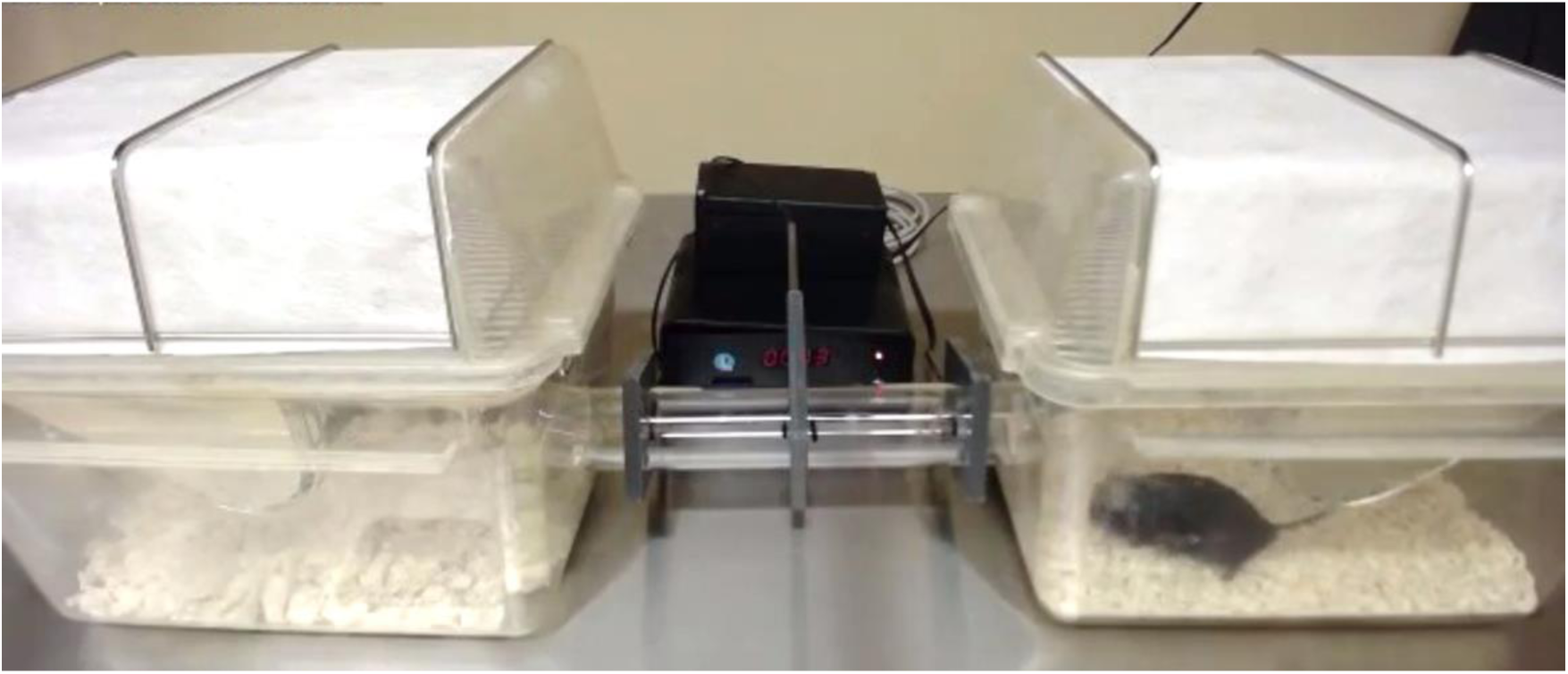
Setup 1 as used during the experiments 1 and 2. Two cages were connected via a tunnel, with either bedding material or different types of gravel as conditioned stimuli (CS). The picture is a screenshot from the videos which were used to get the start time.

For conditioning sessions, an independent type II cage was used. Note that, therefore, this setup did not support the usage of external spatial cues because conditioning sessions were performed in a cage differing from the test cage (similar to a one-compartment design, see Cunningham & Zerizef (2014)).

#### 3.1.3. Procedure

In the first CPP, two experimental procedures, weighing and fixation (restraint by hand), were compared using setup 1. The CS, two different bedding materials, were presented before <u>and</u> after the US.

Weighing was conducted in the same manner as during the weekly cage cleaning and was expected to have a neutral effect. Fixation was conducted on a lid and was expected to have an aversive effect (Stuart & Robinson, 2015; Spyrka & Hess, 2018). Note that animals experienced this procedure before for health monitoring at the arrival and Meloxicam application before transponder implantation. The experiment had a 10 day schedule (see section General Procedure).

Baseline test, conditioning sessions, and final preference test were not conducted in the husbandry room but in a separate, experimental room. Mice (n = 13) were placed in a transportation cage and moved to the experimental room. There, the mice were given 30 min to habituate to the environment. After the experimental procedures, mice were transported back to the husbandry room and returned to their home cage system.

During baseline and final preference test, mice were placed in one of the connected cages. Mice were free to move between cages, and their position was recorded for 10 min by an automated positioning system based on light barriers (as described above).

For the conditioning sessions, the mouse was taken out of the transportation cage and placed in a conditioning cage with bedding material as CS. Each mouse stayed in this cage for 5 min, before the respective procedure (fixation or weighing) was performed. If the mouse was hesitant to leave the handling tunnel (into the glass jar for weighing or onto the lid for fixation), the tunnel was turned or tilted until the mouse softly slipped out of it. After the experimental procedure, the mouse was then again placed for 5 min into the conditioning cage. Afterwards it was returned to the transportation cage.

#### 3.1.4. Results

Already during the baseline test, mice had a strong preference for one of the cues: Mice spend significantly more time in the cage with the comfort white bedding than the cage with the pure bedding (59.95 ± 7.54 %; p < 0.001, t = -4.57). However, in this first experiment the analysis was not conducted before the start of the conditioning sessions, so we continued with the experiment unaware of the bias. After the conditioning sessions in the final preference tests, preference for comfort white was unchanged (see Figure 2, 64.88 ± 9,52 %; p < 0.001, t = - 5.4159). No preference for US or a side bias were observed. Thus, pairing the bedding material with a supposedly negative (fixation) or a neutral (weighing) procedure had no impact on the preference for the bedding material.

**Figure 2:**
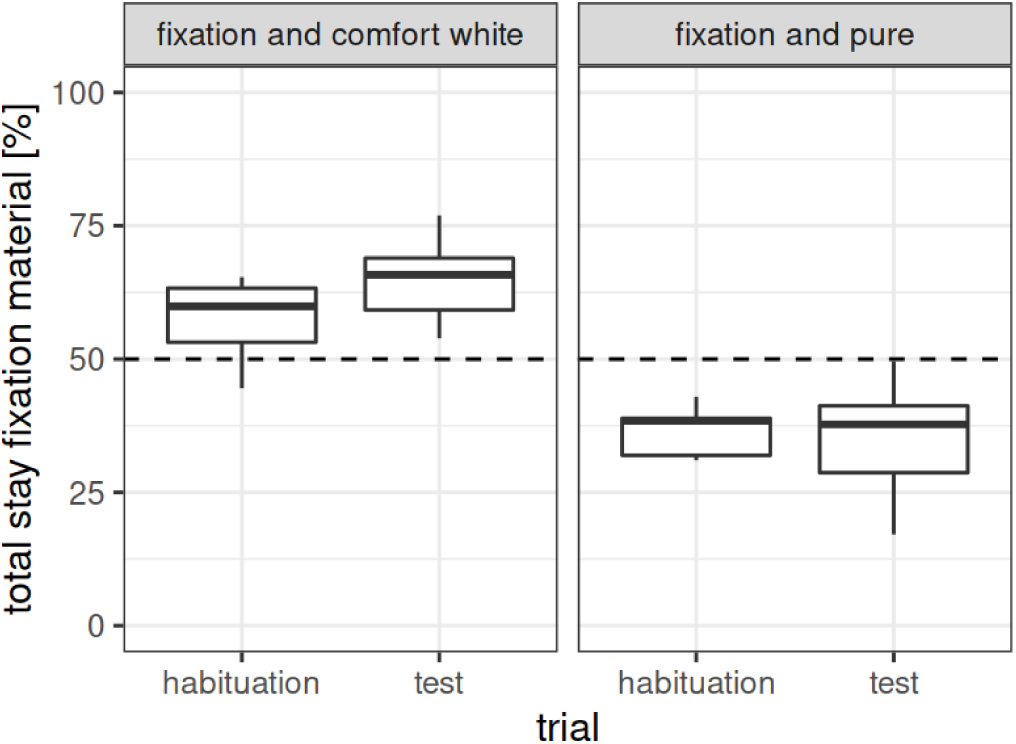
Duration of stay of experiment 1 (n = 12). Depicted is the time spent (in percent) on the bedding paired with a specific procedure (US). For a better visual impression, we split the results with regard to cue combination (CS – US pairing) and trial: In the habituation (= baseline) trial the initial preference pre-conditioning can be seen, whereas in the test trial, the post-conditioning preference can be seen. Thus, with successful conditioning, <u>both</u> subsets of cue combinations should show a decrease <u>or</u> increase from baseline to test. Dotted lines represent chance level (50 %). US: fixation or weighing, CS: comfort white or pure bedding material.

### 3.2. Experiment 2

#### 3.2.1. Procedure

The results from experiment 1 indicated that the used CS (bedding material) had a strong influence itself. No conditioning effect was apparent.

Therefore, as a second experiment, we repeated experiment 1 with the same group of mice (group 1), now using different flooring materials: two different types of gravel. Gravel has the advantage of providing multiple cues (tactile, visual and potentially also olfactory) but is less easily manipulated by the mice and thus, probably less interesting as stimulus in itself (and thus, no potentially competing US).

The same two experimental procedures, weighing and fixation (restraint by hand), were compared, and we used the same setup (setup 1). Because some studies report opposing conditioning effects if the CS are presented before or after the US (see section Introduction and section Timing of US and CS), we now only presented it <u>before</u> the US, in case this caused the missing conditioning effect.

Also, to shorten the time needed for an experiment, we changed the time schedule from 10 days to 6 days (based on Calcagnetti & Schechter (1992); Wang, Wang, & Chen (2014); Oppong-Damoah et al. (2019), more details are given in section General Procedure). Conditioning sessions were shortened from 5 to 3 min to prevent prolongation of the procedure into the afternoon. Everything else resembled experiment 1.

#### 3.2.2. Results

To be sure that there was no initial cue preference, this time, baseline preference was tested and analysed first before proceeding to the conditioning. A first comparison between pumice and marble gravel revealed a preference for pumice (64.85 ± 10.78 %, p < 0.001, t = 4.9692), probably because marble gravel is colder to the touch than pumice gravel with its air enclosures. In a second baseline test, marble gravel was maintained but this time compared to quartz gravel. Here, mice showed no preference for either of the materials (quartz: 53.08 ± 10.95 %; p = 0.3731, t = 0.93244). Thus, we used these two materials for the conditioning.

It has to be noted that just by random matching of mice and US – CS pairing, there was a statistic preference for one of the procedure compartments before the actual conditioning procedure (weighing compartment: 56.98 ± 8.77 %; p < 0.05, t = -2.6394). However, this preference was not apparent in the final test after the conditioning (weighing compartment: 53.04 ± 19.48 %; p = 0.6164, t = -0.51702). No preference for cue or side was found. As can be seen in Figure 3, the variability of data in the final test nearly doubled compared to the baseline test.

**Figure 3:**
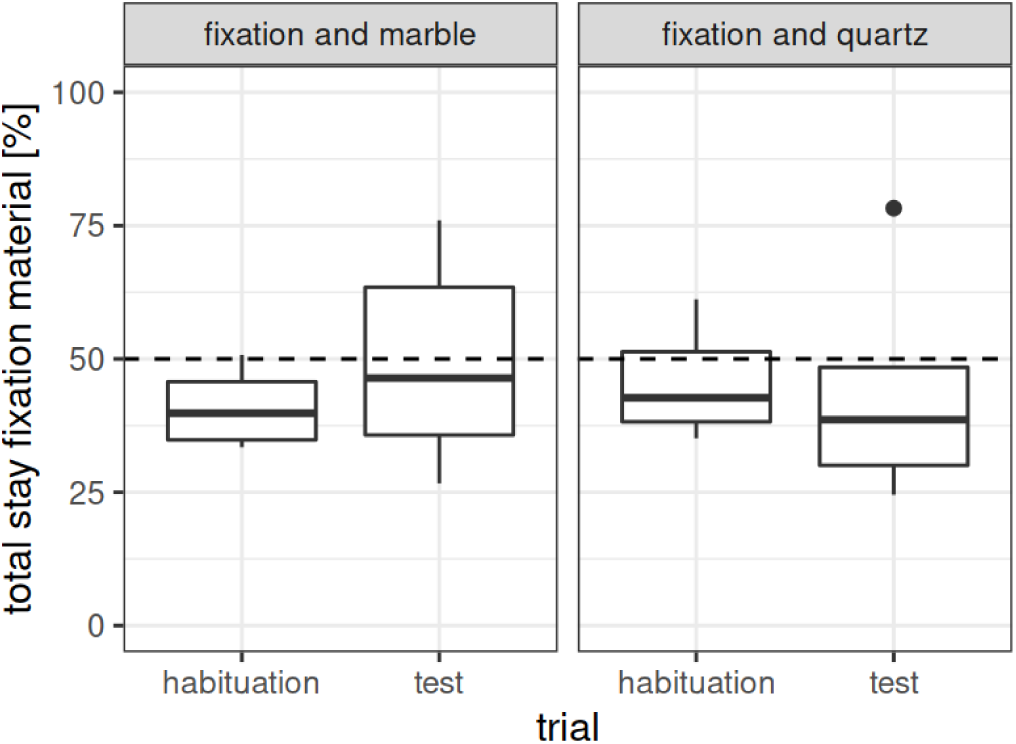
Duration of stay of experiment 2 (n = 11). Depicted is the time spent (in percent) on the pattern paired with a specific procedure (US). For a better visual impression, we split the results with regard to cue combination (CS – US pairing) and trial: In the habituation (= baseline) trial the initial preference pre-conditioning can be seen, whereas in the test trial, the post-conditioning preference can be seen. Thus, with successful conditioning, <u>both</u> subsets of cue combinations should show a decrease <u>or</u> increase from baseline to test. Dotted lines represent chance level (50 %). US: fixation or weighing, CS: marble or quartz gravel.

### 3.3. Experiment 3

#### 3.3.1. Procedure

In the previous experiment, the gravel itself still seemed to have an influence on the behaviour of the mice and thus, might function as an US in itself. To reduce this effect, we changed the CS to be more similar to the one used by Cunningham, Gremel & Groblewski (2006), using plates (“grid” versus “hole” floor structure). This reduced the CS to tactile cues only.

In addition, the setup was altered to a more common design, also more similar to the one described by Cunningham, Gremel & Groblewski (2006): setup 2, see below. Because of the setup change, the former automatic analysis was now done manually, by analysing video recordings. The CS was again presented <u>before</u> the US.

As in experiment 2, we used a 6 day schedule, and the same group of mice participated (group 1, n = 12), as no previous conditioning effect was apparent. However, we slightly adjusted the US: We considered that the surface on which each procedure is performed might operate as a CS in itself, especially as during fixation mice have a prominent contact with the surface. To prevent this, we harmonized the surfaces for both procedures: Instead of using different containers for the two procedures (experiment 1 and 2 weighing: glass, fixation: lid), we used now a small cage. For weighing it was used the right way round, and for fixation it was turned upside down. Thus, the tactile cue of the surface was similar.

In addition, to prevent additional potentially stressful effects, we now refrained from forcing the mice out of the handling tunnel. When placing the mouse on the surface, the back opening of the handling tunnel was sealed by hand and the experimenter waited until the mouse left the tunnel by itself (in rare cases, this lasted up to a minute). As we observed differences in the latencies to leave the tunnel during the conditioning sessions between the procedures, we recorded them on the last conditioning day. However, as we had no baseline recordings, this data stays anecdotal and is described in the Supplements only.

Note that during the conduction of the experiment it was observed that both metal plates in the conditioning setup got noticeably cold after the cleaning with ethanol between mice (for more details see Supplements).

#### 3.3.2. Setup 2: One cage separated by a barrier

Setup 2 was used for baseline test, conditioning sessions and the final test in experiments 3 – 6. The second setup was designed based on the model of Cunningham, Gremel & Groblewski (2006): Guiding bars were added to a type III cage (LWH: 425 x 276 x 153 mm, Tecniplast) to apply a small barrier (2 cm height) or a plate (about 13 cm height). Thus, the cage was divided into two areas of the same size, which contained different CS (see Figure 4A and 4B). The small barrier was used during the habituation and preference test to have an actual visual and physical separation of the two areas. The plate was used during conditioning sessions to restrict the mice to one compartment.

**Figure 4:**
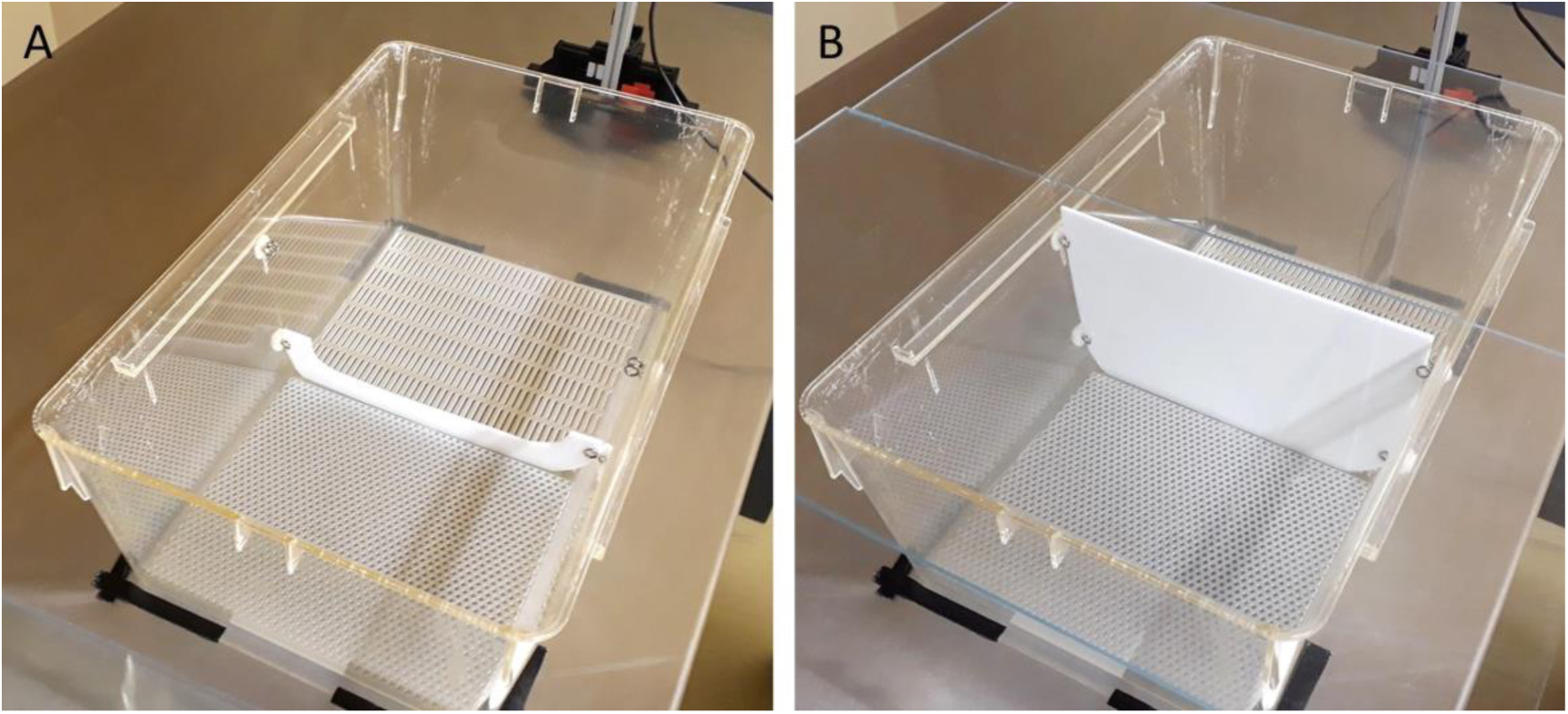
Setup 2 as used during the experiments 3 – 6. (A) Barrier and (B) plate: One cage was separated by a barrier, with either different plates or floor patterns as CS.

Note that as the outside walls were transparent, this setup allowed external spatial cues, such as proximity and colour of walls. In this manner, every CS used in this setup also included a spatial cue (left vs. right side of the setup). In this setup, the CS were positioned at the floor only, either metal plates (tactile cues, experiment 3 and 4) or patterns (visual cues, experiment 5 and 6).

The setup was closed by a Perspex plate, so that video recordings were possible from above. The camera was applied to a metal construction, which was built in the first experiment out of beams by fischertechnik GmbH, Germany, and later replaced by MakerBeam B.V., The Netherlands, see Supplements.

For video analysis, the time spent in each component of the setup was recorded manually (experiments 3 and 4) or with the help of the open source program BORIS (experiments 5 – 6, Behavioral Observation Research Interactive Software, Version 7.9.8, (Friard & Gamba, 2016)). A change of compartment was counted whenever a mouse moved with all four paws over the barrier in the other compartment. Time in-between, when the mouse was partly in one and partly in the other compartment, was counted as still belonging to the side the mouse was last.

#### 3.3.3. Results

The baseline test revealed no preference for cue, side or procedure compartment. Visually comparing the results of the baseline with baseline results of the previous CPP experiments (using setup 1), it is noticeable that the variability range of the data is smaller than before. However, the variability became larger again during the final test (see Figure 5).

**Figure 5:**
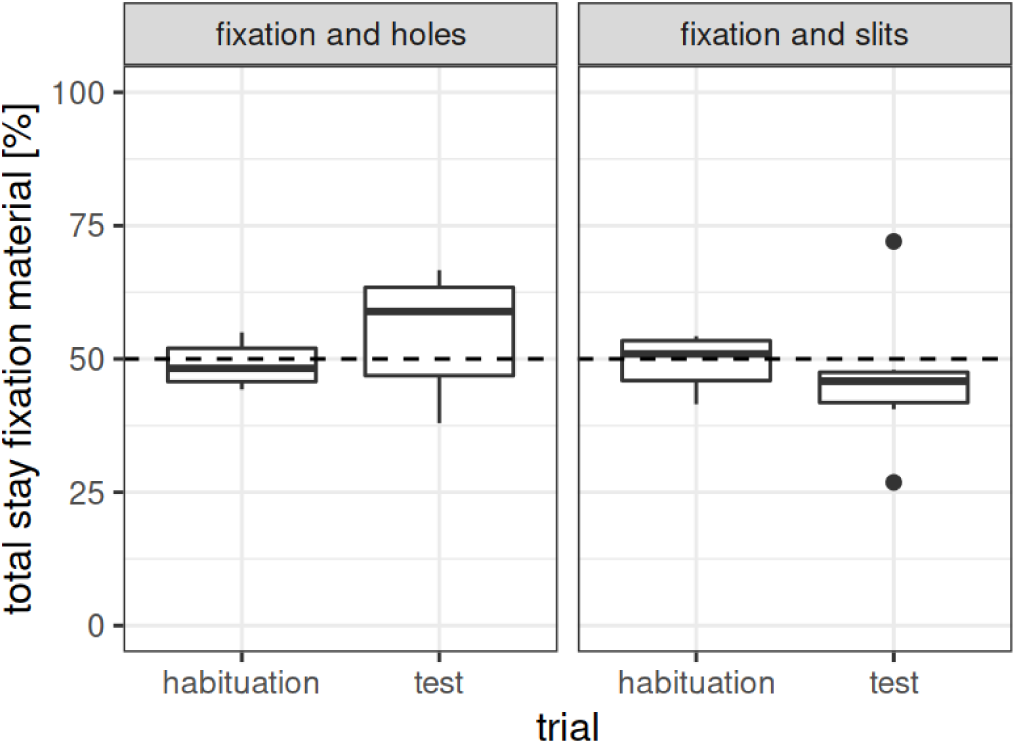
Duration of stay of experiment 3 (n = 12). Depicted is the time spent (in percent) on the pattern paired with a specific procedure (US). For a better visual impression, we split the results with regard to cue combination (CS – US pairing) and trial: In the habituation (= baseline) trial the initial preference pre-conditioning can be seen, whereas in the test trial, the post-conditioning preference can be seen. Thus, with successful conditioning, <u>both</u> subsets of cue combinations should show a decrease <u>or</u> increase from baseline to test. Dotted lines represent chance level (50 %). US: fixation or weighing, CS: holes ot slits metal flooring plates.

In the final test there was no preference of the mice for procedure compartment, cue or side (see Tables 2, 3 and 4). Thus, although we took special care to harmonize the experimental procedures with regard to the surface to exclude it as an involuntary CS, no preference for a procedure compartment was measurable. It is possible that the temperature of the plates after cleaning with ethanol might have affected the conditioning procedure.

In general, comparing setup 1 and 2, it is noticeable that the mice changed compartments more often (setup 1: between 0 and 12, setup 2: between 6 and 59 compartment changes per mouse and test).

### 3.4. Experiment 4

#### 3.4.1. Procedure

After experiment 3, we considered that the nature of the fixation procedure (pushing the mouse onto the surface) might suppress learning any other CS than the surface itself. For the fourth CPP, we therefore changed the US. We kept weighing as a supposedly neutral procedure and compared it with food reward using millet as a supposedly positive procedure. We decided upon this US as there are already CPP studies using food reward (Imaizumi, Takeda, & Fushiki, 2000; Takeda et al., 2001).

We enrolled group 2 (n = 12), which was naive to the conditioning procedure. We kept the setup from before (setup 2) and the same CS, i.e., metal plates with holes or slits. Again, a 6 day schedule was used, and for the conduction of the experiment, mice were transported to a separate, experimental room (as in experiments 1 to 3).

In the previous experiment, mice defecated and urinated noticeably, which could be a sign of fear (Gray & McNaughton, 2003; Hurst & West, 2010). To reduce this, group 2 was habituated to the setup on three days for 1 min in their husbandry room before the baseline test. Note that many studies (including the protocol by Cunningham, Gremel & Groblewski (2006)) do not perform additional habituation sessions beyond the baseline measurement, which was the reason why we did not perform habituation sessions in the preceding experiments.

To ensure that the mice would consume the millet during the conditioning sessions, on 3 days before the start of the experiment in total 6 g of millet were placed into the home cage in the morning after the onset of the light phase.

The glass jar for weighing was the same as used in experiment 1 and 2 but unfamiliar to the new group until the start of the experiment (i.e., a different container was used for the weekly weighing). For the food reward procedure, a mouse was placed into a type III cage filled with bedding material, which had 0.1 g millet at one end of the cage.

During both procedures, the time to leave the handling tunnel onto the surface and the time, when the mouse re-entered the tunnel after the procedure was noted for all conditioning sessions. In addition, during the millet procedure, it was also noted, when the mouse began feeding and when it stopped to do so. This was done to see if a change in behaviour occurred over time. To not extent the scope of this article, statistics and results are reported in the Supplements.

Due to the observations in experiment 3 regarding the temperature change of the plates after cleaning, in this experiment we waited at least 90 s after the cleaning, to return to approximately room temperature, before a mouse was placed onto the plates.

#### 3.4.2. Results

There was no preference for procedure compartment, cue or side during the baseline test. In the final preference test, there was a tendency towards a preference for the slit plate (cue), although it did not reach significance (see Figure 6, 54.07 ± 7.02 %; p = 0.0696, t = 2.01). In addition, comparing the results before and after conditioning, there was a tendency for an increased side preference (p = 0.08055, t = -1.9244). However, there was no preference for a procedure compartment (millet compartment: 51.27 ± 8.10 %; p = 0.5993, t = 0.54097). This time, standard deviations of baseline and final test were similar.

**Figure 6:**
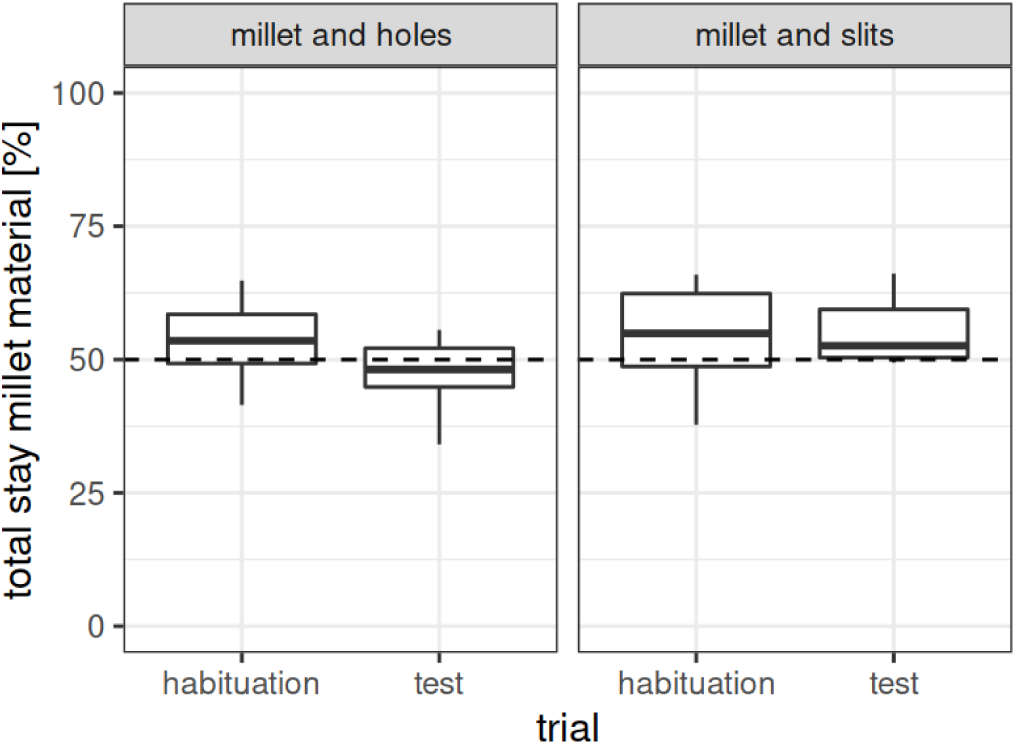
Duration of stay of experiment 4 (n = 12). Depicted is the time spent (in percent) on the pattern paired with a specific procedure (US). For a better visual impression, we split the results with regard to cue combination (CS – US pairing) and trial: In the habituation (= baseline) trial the initial preference pre-conditioning can be seen, whereas in the test trial, the post-conditioning preference can be seen. Thus, with successful conditioning, <u>both</u> subsets of cue combinations should show a decrease <u>or</u> increase from baseline to test. Dotted lines represent chance level (50 %). US: millet or weighing, CS: holes vs. slits metal flooring plates.

### 3.5. Experiment 5

#### 3.5.1. Procedure

Between CPP experiment 4 and 5, we conducted two other experiments, to follow a different strategy than CPP to compare the effect of procedures. A description of them can be found in the Supplements (experiments 4.1 and 4.2).

Looking at the preceding CPP experiments and their inconclusive results, it seemed clear to us that a fundamental element in the experiments was not working. Therefore, we decided to reproduce a “basic” protocol of the CPP as closely as possible. Only after a successful reproduction of the protocol we wanted then to move on and alter it to compare experimental procedures.

To do so, we used fluids as US similar to Imaizumi, Takeda & Fushiki (2000); Takeda et al. (2001); Matsumura et al. (2010). Here, we compared tap water and almond milk, as we already knew from other studies in our research group (Kahnau, Jaap, et al., 2023) that almond milk is a preferred good. We used setup 2, which is designed similar to Cunningham, Gremel & Groblewski (2006)). For the presentation of fluids during the conditioning sessions, a Perspex plate with a hole was placed on top of the compartment and the nipple of the fluid bottle was inserted through the hole. Mice were filmed to monitor whether the mice drank the fluids. After 3 min, the mouse was returned to the transportation cage.

As CS visual patterns were used (designed as described by Cunningham, Gremel & Groblewski (2006)). The CS were presented <u>simultaneously</u> with the US. We again conducted the 6 day schedule. Baseline test, conditioning sessions and final preference test were conducted in a separate, experimental room (for logistic reasons not the same as in the experiments before but it had similar conditions). As group 2 (n = 12) showed no apparent conditioning in the previous experiment, it participated again in this experiment. As they were already familiar with the general setup (not the CS), no additional habituation was deemed necessary.

#### 3.5.2. Results

During the baseline test, there was no preference found for side, pattern or the procedure compartment. During the sessions, it was noted that mice seldom tasted the fluids and never actually drank them. In addition, due to dribbling of the bottles the floor was expectantly wet.

In the final preference test, there was also no side, cue, or fluid preference (see Tables 2, 3 and 4). Comparing the results from the baseline test (pre-conditioning) with the final preference test (post-conditioning), there was no change in duration of stay on the pattern paired with almond milk (procedure compartment) or side. Interestingly, however, there was a significant increase in duration of stay on the dot pattern (cue; see Figure 7, p < 0.01, t = -3.2257).

**Figure 7:**
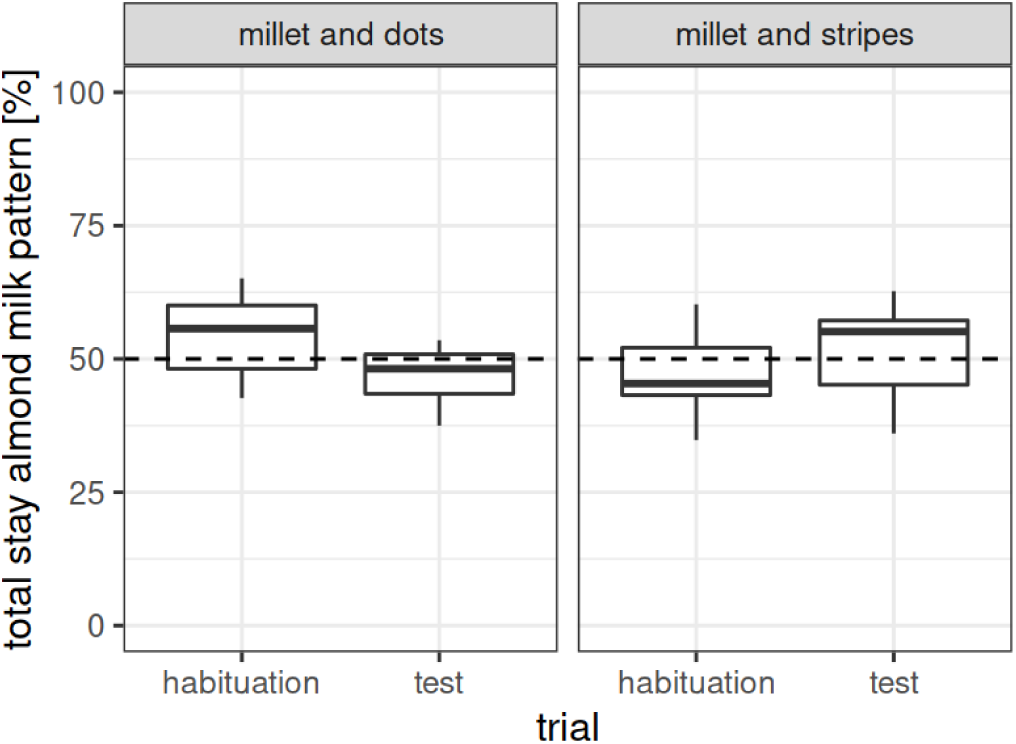
Duration of stay of experiment 5 (n = 12). Depicted is the time spent (in percent) on the pattern paired with a specific procedure (US). For a better visual impression, we split the results with regard to cue combination (CS – US pairing) and trial: In the habituation (= baseline) trial the initial preference pre-conditioning can be seen, whereas in the test trial, the post-conditioning preference can be seen. Thus, with successful conditioning, <u>both</u> subsets of cue combinations should show a decrease <u>or</u> increase from baseline to test. Dotted lines represent chance level (50 %). US: almond milk or water, CS: dots or stripes visual pattern.

It has to be noted, that not all of the mice tasted the fluid during conditioning sessions and only one mouse was observed actually drinking. Thus, the actual pairing of US and CS might not have taken place (no experiencing the US, for more details on behavioral observations during the experiment see Supplements).

### 3.6. Experiment 6

#### 3.6.1. Procedure

In the preceding experiment, the consummation of the fluids was low, which interfered with the mice actually experiencing the US. To determine under which conditions almond milk or millet were consumed by the mice in a new environment, before CPP experiment 6, a series of pre-tests was performed with the same group of mice (group 2, n = 12). We observed that millet was consumed more readily than almond milk in a separate cage, with more mice feeding on some of the grains while no tasting of the fluids was observable. In addition, consummation rates increased with repetition. Therefore, we got to the conclusion that habituation (or the lack thereof) might play an important role.

Thus, we repeated an alternate version of experiment 5: The whole experiment was now conducted in the housing room to avoid a new environment. Instead of placing the mice first into a transportation cage, mice were directly taken out of their home cage to be placed in the conditioning setup. To facilitate the handling of the mice, i.e., placing it in or taking it out of the home cage, the filter tops of the home cage system were removed at the start of each experimental day. The mice were given 10 min to habituate to the changed light condition before the start of the experiment. The filter tops were put back on top of the cages after the last mouse had finished its session. In addition, to exclude the ethanol odour as an additional cue or stress factor (for more details see Supplements and Discussion section Choice of CS), we changed our cleaning procedure from ethanol to water and cleaned only if mice urinated or defecated.

Moreover, we focused on the advantage of millet consumption compared to almond milk shown in the pre-tests, with the goal of finding first a working protocol and altering it afterwards to specific additional procedures. Thus, as US either millet mixed with bedding material (= millet) or only bedding material (= no millet) was provided. For conditioning sessions, mice were placed for 3 min individually into one half of the conditioning setup. A Perspex plate with a hole was placed on top of the compartment to prevent escape. On the floor of the cage, a small amount of bedding material mixed either with 0.1 g millet or no millet was available for the mice. To monitor whether the mice consumed the millet, sessions were video recorded.

Mice were already familiar with millet and feeding in a type III cage due to the pre-tests. We again used setup 2, which is very similar to the type III cage of the pre-tests. Two self-designed patterns consisting of black and white blocks leading to a fabric texture-like and a chessboard pattern were used as CS. We returned to a 10 day schedule as conducted in experiment 1 (similar to the protocol of Cunningham, Gremel & Groblewski (2006)) as this seems to be the more conventional procedure.

#### 3.6.2. Results

During the baseline test, there was no preference for side, pattern or procedure compartment (see Tables 2, 3 and 4). During conditioning session, mice readily ate the millet. However, there was also no preference for procedure compartment, cue or side in the final preference test (see Figure 8). There was no change in duration of stay comparing pre-and post-conditioning.

**Figure 8:**
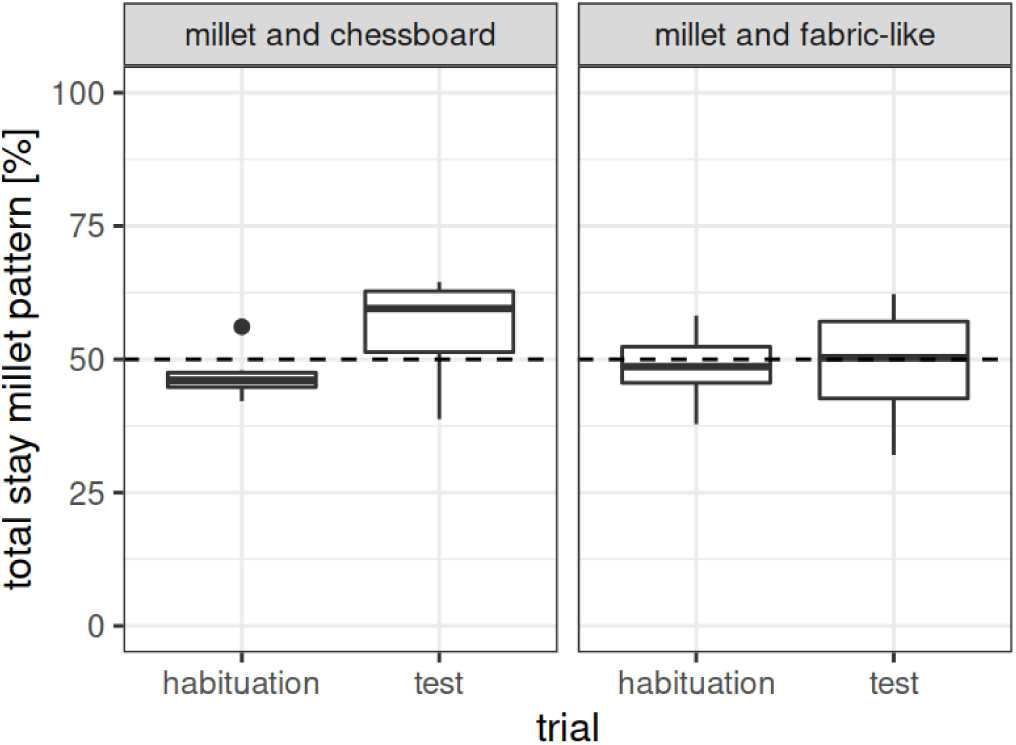
Duration of stay of experiment 6 (n = 12). Depicted is the time spent (in percent) on the pattern paired with a specific procedure (US). For a better visual impression, we split the results with regard to cue combination (CS – US pairing) and trial: In the habituation (= baseline) trial the initial preference pre-conditioning can be seen, whereas in the test trial, the post-conditioning preference can be seen. Thus, with successful conditioning, <u>both</u> subsets of cue combinations should show a decrease <u>or</u> increase from baseline to test. Dotted lines represent chance level (50 %). US: millet with bedding or only bedding, CS: chessboard or fabric-like visual pattern.

### 3.7. Experiment 7

#### 3.7.1. Procedure

A possible confounding factor of experiment 6 might have been that the patterns were not easily distinguishable for the mice. In addition, it is possible that the mice have to experience the onset of the experimental procedure in the respective compartment for a successful conditioning. This is based on thoughts from a study by Goltseker & Barak (2018) and similar to foraging strategies as discussed already by Herrmann et al. (1982). Why should the mouse “wait” in the final test near the millet-CS when it has never experienced the “filling” of the millet? In this case, the mouse might expect that the consumed millet will not be refilled, and therefore, the former millet-environment might be considered as empty as the no-millet-environment.

As a consequence, the procedure for experiment 7 was similar to the procedure in experiment 6, with the following alterations:

Firstly, to reduce potential visual influence from the outside environment, we used a new setup (setup 3, description below) instead of setup 2, which had opaque instead of transparent walls (based on Sun et al. (2018)). The compartments were now separated by a wall either with or without a hole, which reduced the view into the other compartment.

Secondly, a new group of mice was enrolled, which was naive to the CPP test (group 3, n = 12). Because this group was also naive to the CPP setup, we had four 3 min sessions of habituation (without the CS). This was done to ensure that mice would feed on the millet during the actual experiment.

Thirdly, visual pattern of horizontal or vertical black and white stripes were used as CS (similar to Lett et al. (2001)). These patterns have also been used by Wong & Brown (2007) and validated as distinguishable for C57BL/6 mice in the respective age. The patterns were applied to three of four walls of each compartment (compare setup 2: CS on the floor).

Fourthly, mice were first placed inside the compartment without any US being present. After 30 s the US was added: either 0.1 g millet or a visually similar amount of bedding material. With this, we wanted to ensure that the patterns were perceived before the US was presented.

Apart from that, we used the same 10 day schedule as in experiment 6. Procedures were conducted in the husbandry room (see experiment 6).

#### 3.7.2. Setup 3: Two compartments separated by a wall

Setup 3 was used for baseline test, conditioning sessions and the final test in experiment 7 – 9. Two compartments (LWH: 32 x 11 x 20 cm) consisting of grey plastic were joint together (see Figure 9A, based on Sun et al. (2018)). In experiment 7 and 8 the floor consisted of smooth grey plastic. In experiment 9, we placed 3D printed black plates containing structures on the floor to add tactile cues. Three of the four walls were additionally covered with Perspex plastic, behind which a sheet of laminated paper with a pattern could be placed. The fourth wall was the connection to the other compartment and was removable. This wall contained either a hole (experiment 7 and 8: 6 cm width and 7 cm height, experiment 9: 6 cm diameter) to allow the mice to change compartments (for habituation, baseline and test), or not (conditioning sessions). Between both compartments, there was a small area of approximately 2 cm length. In experiment 7 and 8, this was not altered, leaving the same grey plastic for floors and walls as for the rest of the setup. In experiment 9, the small area between the compartments was filled with a fitting wall of 3 cm width (3D printed, black PLA). Accordingly, the separation wall for the conditioning sessions was printed out of the same material.

**Figure 9:**
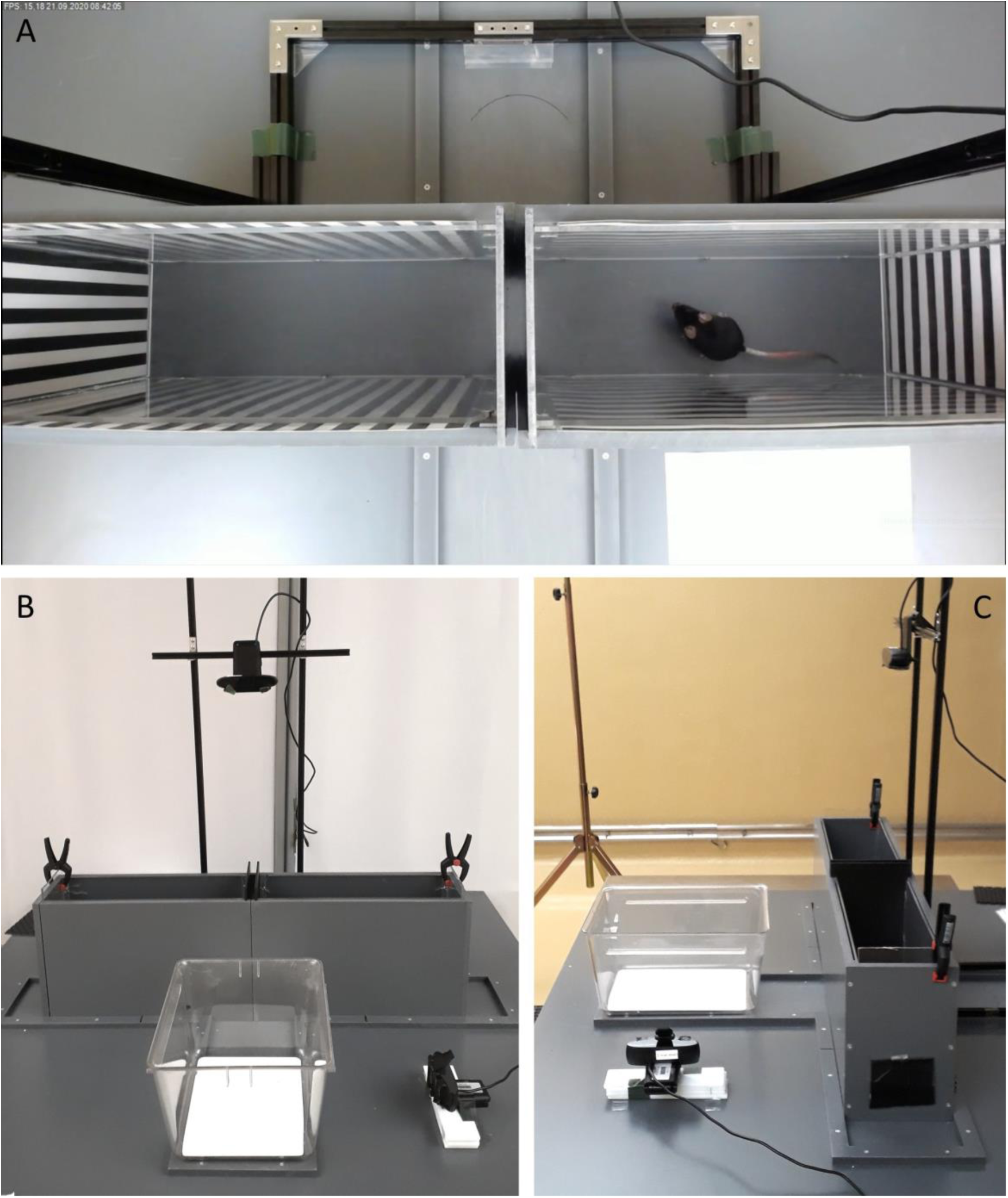
Setup 3 used during the experiments. Two compartments are separated by a wall, used during experiments 7 – 9 with different wall patterns as conditioned stimuli (CS). (A) View from above, picture taken as a screenshot from the video recordings used for analysis. (B) Front view and (C) side view of the setup. In experiment 9, the separator between the two compartments did not contain an open space and the floor was covered with additional 3D printed plates. The type III cage depicted in (B) and (C) was used in experiment 9 for conditioning sessions, with millet or restraint in a restrainer as procedure.

A metal construction (MakerBeam B.V.) was built to hold a camera, so that the apparatus could be filmed from above. Note that this setup excluded external spatial cues due to its opaque walls.

For video analysis, the time spent in each component of the setup was recorded with the help of BORIS. The area between the two compartments was very small and a mouse could still be partly in one of the compartments. Therefore, we counted a mouse as leaving one compartment, as soon as it had its head in the area between the compartments. We counted a mouse entering a compartment when all four paws were inside it. Duration of stay was normalised for time spent in one of the compartments (excluding the time in-between compartments).

#### 3.7.3. Results

In the baseline test, there was a significant preference found for the left half but not for pattern or procedure compartment. Note that the setup here in comparison to the last experiment had opaque walls, and therefore, should have excluded environmental effects too an even higher degree. We reasoned that an effective conditioning should erase the side preference and thus continued with the test without additional changes.

However, in the final preference test, the significant side preference was still apparent (see Figure 10, left: 56.38 ± 7.38 %; p = 0.01224, t = 2.9925), while there was no pattern preference (horizontal stripes: 50.10 ± 9.95 %; p = 0.9718, t = -0.036123) and only a tendency towards procedure preference (millet: 54.83 ± 8.57 %; p = 0.07697, t = 1.9511). Comparing the results from the baseline test (pre-conditioning) with the final preference test (post-conditioning), the preference did not change for any of the factors (including procedure compartment).

**Figure 10:**
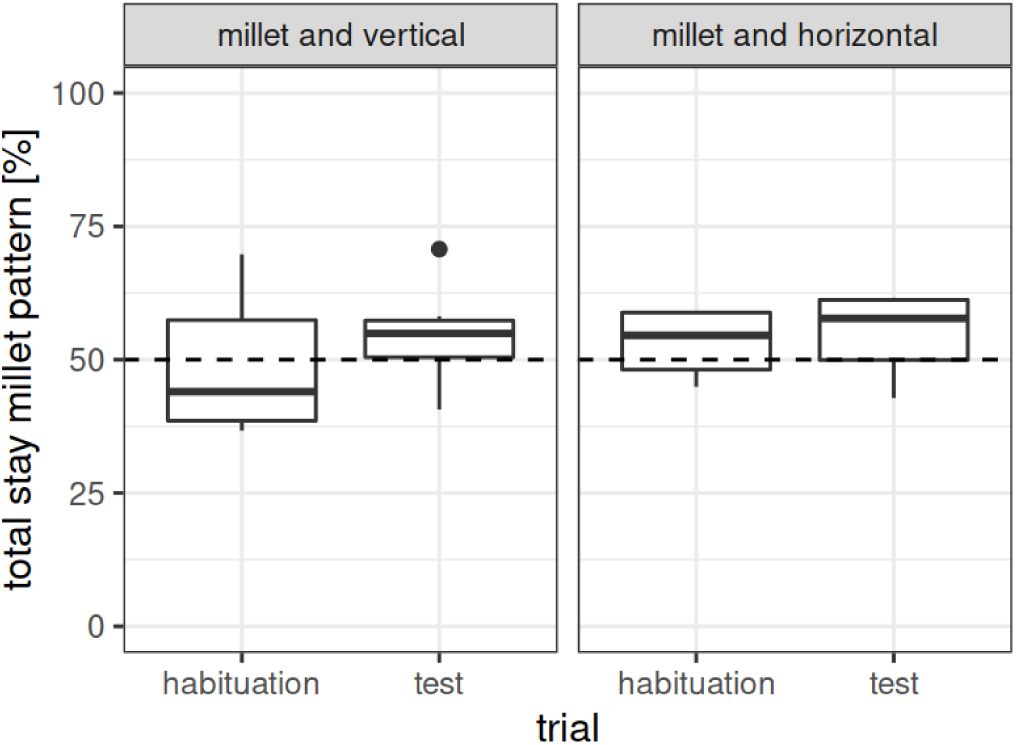
Duration of stay of experiment 7 (n = 12). Depicted is the time spent (in percent) on the pattern paired with a specific procedure (US). For a better visual impression, we split the results with regard to cue combination (CS – US pairing) and trial: In the habituation (= baseline) trial the initial preference pre-conditioning can be seen, whereas in the test trial, the post-conditioning preference can be seen. Thus, with successful conditioning, <u>both</u> subsets of cue combinations should show a decrease <u>or</u> increase from baseline to test. Dotted lines represent chance level (50 %). US: millet or bedding, CS: vertical or horizontal stripes as visual pattern.

### 3.8. Experiment 8

#### 3.8.1. Procedure

Until now, all CPP experiments were designed as forward conditioning, meaning first the CS was presented and then the US (except for experiment 1, were the CS was presented both before and after the US). By first presenting the US and then the CS, backward conditioning is also possible (Lett et al., 2000, 2001; Lett, Grant, & Koh, 2002; Belke & Wagner, 2005; Nakajima, 2020). This was aimed for in experiment 8.

Group 3 (n = 12) took part and we used setup 3 (opaque walls and compartments separated by a wall either with or without a hole) and a 10 day schedule. Due to the altered US – CS timing, the US procedures were adapted: As a supposedly positive procedure (millet), mice were placed individually in a bedding filled cage, into which immediately 0.1 g millet were given. Mice had 1 min to consume the millet, before they were taken out of the cage again. As a supposedly negative, stressful procedure (restrainer, Glavin et al. (1994); Zimprich et al. (2014)), mice were placed in a bedding filled cage and then immediately transferred into a restrainer, in which they had to stay for 1 min.

Because the mice obviously did not learn any associations with the CS in the experiment before, we used the same visual CS (horizontal or vertical stripes). However, we made sure that the formerly positive paired CS (millet) for each mouse was now paired with the negative, stressful procedure.

Experimental procedures were conducted in the husbandry room. Due to the backward conditioning, mice were now directly taken out of their home cage and placed in the cage in which the procedure (millet or restrainer) took place. After the procedure conduction, mice were transferred into the conditioning setup, in which they stayed for 3 min, before they were taken out and returned to their home cage.

#### 3.8.2. Results

In the baseline test, there was no significant preference found for side, pattern or the procedure compartment (see Tables 2, 3 and 4). In the final preference test, there was again no significant side, pattern or procedure compartment preference (see Figure 11). Comparing the results from the baseline test (pre-conditioning) with the final preference test (post-conditioning), the duration of stay near the vertical pattern increased (p < 0.01, t = -3.1647).

**Figure 11:**
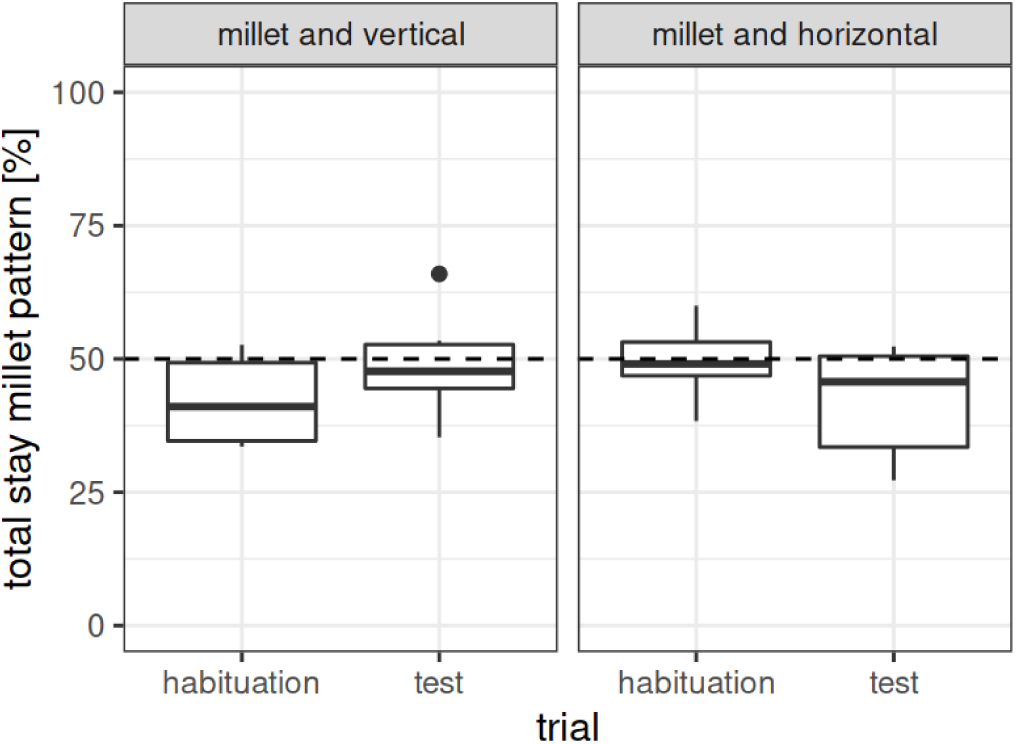
Duration of stay of experiment 8 (n = 12). Depicted is the time spent (in percent) on the pattern paired with a specific procedure (US). For a better visual impression, we split the results with regard to cue combination (CS – US pairing) and trial: In the habituation (= baseline) trial the initial preference pre-conditioning can be seen, whereas in the test trial, the post-conditioning preference can be seen. Thus, with successful conditioning, <u>both</u> subsets of cue combinations should show a decrease <u>or</u> increase from baseline to test. Dotted lines represent chance level (50 %). US: millet or restrainer, CS: vertical or horizontal stripes as visual pattern.

### 3.9. Experiment 9

#### 3.9.1. Procedure

Between experiment 8 and 9, one additional experiment took place. It was not based on CPP but focused on behavioral changes, and is therefore reported only in the Supplements (experiment 8.1). Experiment 9 was conceptualized as a repetition of experiment 8 with a different group of mice (group 4, n = 12), which were fully whiskered (in contrast to group 3). We used setup 3 (opaque walls and compartments separated by a wall either with or without a hole) with the same visual CS (horizontal or vertical stripes). Moreover, an additional CS was added to the floor to potentially increase the conditioning effect (Cunningham, Patel, & Milner, 2006; Cunningham & Zerizef, 2014). For this, plates with tactile structures were used, designed after the study by Mei et al. (2020) with an 8-arm maze.

We kept the US (millet and restraint) but this time, mice were placed in a type II cage (LWH: 225 x 167 x 140 mm, Tecniplast) without bedding. Instead, the floor was covered with a white plate with protruding bars or dots (3D printed, PLA). This was done to investigate whether mice formed an association between the tactile floor cues in the conditioning cage and the procedure. If so, we expected mice to show a greater hesitancy to leave the handling tunnel onto the floor combined with the negative procedure, compared to the positive procedure. For this reason, the conduction of the conditioning sessions was video recorded from the side, to record the time the mice took to exit the handling tunnel and enter the cage (for more details on the video analysis and results see Supplements).

During conditioning sessions, mice were placed inside the procedure environment (type II cage) and immediately, either 0.1 g millet were added or the mice were transferred into a restrainer. After 1 min, the mouse was either taken out of the cage and placed into the conditioning setup (millet), or the restrainer was placed into the conditioning setup and opened to release the mice (restrainer).

As mice were unfamiliar with the setup, we had 4 consecutive days of habituation, in which mice were habituated to the procedure environment and the conditioning setup: During these habituation sessions, the procedure environment (type II cage) contained a smooth white plate without additional tactile structures, while the conditioning setup contained smooth black plates without additional tactile structures and no visual wall patterns (plain grey walls). Mice were placed individually for 3 min into the procedure environment and then 3 min into the conditioning setup. Afterwards, they were returned to their home cage. To reduce the time for habituation, we interlaced the procedures, i.e., while one mouse was habituating to the conditioning setup, another mouse was habituating to the procedure environment. Habituation to the millet was not necessary as mice experienced millet as part of an active enrichment in their home cage (see section Housing).

#### 3.9.2. Results

In the baseline test, there was a significant preference for side (left: 56.65 ± 10.42 %; p < 0.05, t = 2.2106), although we used opaque walls and similar light conditions. We argued that this preference should be overcome by a successful conditioning (similar to a “biased” study design, see Cunningham, Gremel & Groblewski (2006)) and continued with this setup.

In the final preference test, there was a tendency towards a preference for the restrainer procedure compartment (see Figure 12, 54.63 ± 8.70 %; p = 0.09202, t = -1.8456). However, if one compares the results from pre-and post-conditioning, it becomes obvious that there is no significant increase of preference (p = 0.8125, t = -0.24293) but probably merely a reduction of data variability. Instead, the side preference remained (left: 56.82 ± 6.97 %; p < 0.01, t = 3.3902). In addition, we did not find a difference between experimental procedures regarding the latency to leave the handling tunnel (for more details see Supplements).

**Figure 12:**
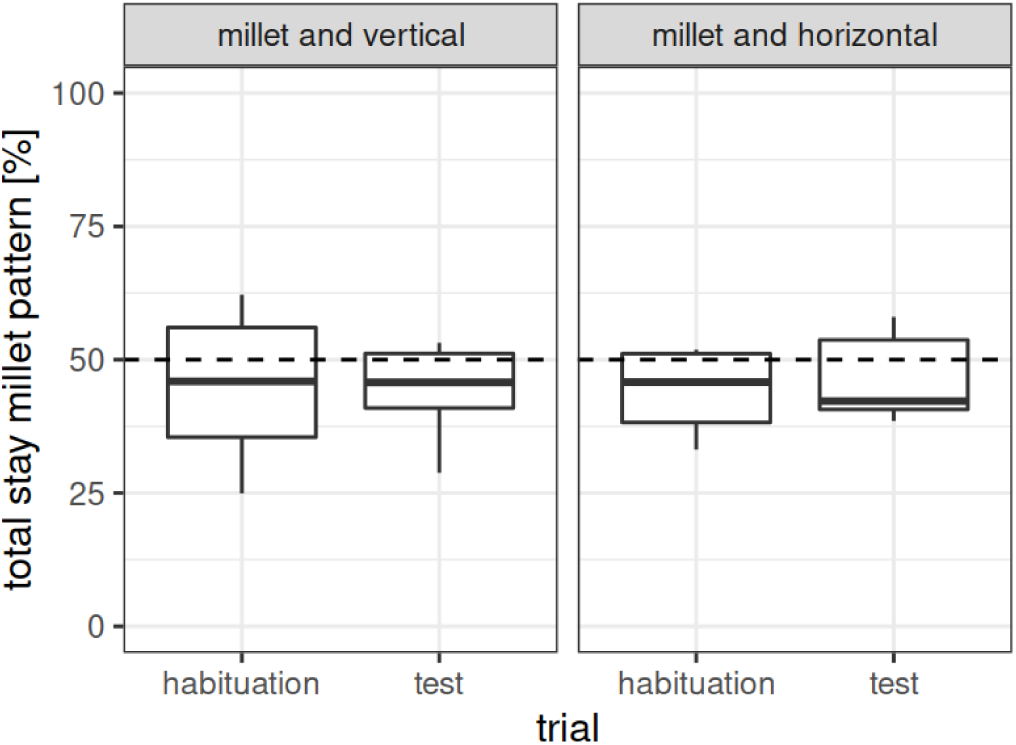
Duration of stay of experiment 9 (n = 12). Depicted is the time spent (in percent) on the pattern paired with a specific procedure (US). For a better visual impression, we split the results with regard to cue combination (CS – US pairing) and trial: In the habituation (= baseline) trial the initial preference pre-conditioning can be seen, whereas in the test trial, the post-conditioning preference can be seen. Thus, with successful conditioning, <u>both</u> subsets of cue combinations should show a decrease <u>or</u> increase from baseline to test. Dotted lines represent chance level (50 %). US: millet or restrainer, CS: vertical or horizontal stripes as visual pattern. Note that in this experiment the visual pattern was also combined with a fixed tactile cue.

## 4. Discussion

Existing successful conditioned place preference / aversion tests show various differences in their conduction, which might give the impression that there are multiple ways towards the goal. However, it seems rather that the specific suitable way has to be found for each specific experimental question which is addressed. Communication with other researchers during this set of experiments gave even the impression that the CPP paradigm seems to be difficult to conduct even for established protocols and stimuli.

Here, we aimed on developing a CPP protocol to pair an experimental procedure as US with a NS → CS to test the emotional valence of the experimental procedure. In this manner, it should be feasible to compare different experimental procedures with regard to their severity from the mice’s perspective.

However, finding such a protocol proved to be a challenge. As we have described above, we took several approaches, none of them leading to the desired effect. All the more, we would like to emphasise some conclusions we can draw from these experiments which might be helpful for future researches. However, it has always to be considered that enrolment of different species, strains, age, sex, housing, or other factors might influence the results to an unknown extent and comparisons have to be done with caution.

### 4.1. Choice of CS

In general, stimuli are not equally effective for all kinds of US (Garcia & Koelling, 1966); therefore, the choice of cues is very important. In the following, we will discuss the used cues; note that as conditioning did not work, they are not per se “CS”, but remained NS. However, to simplify the discussion, we will nevertheless speak of CS.

In experiment 1 and 2, we chose different flooring materials (bedding and gravel) as CS. This was successfully done before in other CPP studies (Meisel & Joppa, 1994; Kennedy et al., 2012; Meyer & Alberts, 2016). Flooring materials combine multiple cues, including visual, tactile and olfactory cues multiple cues which can improve CPP acquisition (Cunningham & Shields, 2018). Thus, mice should have been able to easily discriminate between the two chosen CS in this experiment – which can be confirmed as the mice showed a significant preference for one of the CS from the start. However, this pre-preference is a disadvantage as it might affect the conditioning (e.g., by working as a positive US itself). Thus, although there are some studies, like Lett et al. (2001), which did not perform a baseline preference test, we highly recommend a baseline test to ensure that the preconditions are the same. In addition, it should be analysed before the beginning of the conditioning sessions, to be able to change the CS and start again. In general, of course, a preference for one of the CS can also develop during conditioning sessions, independently from the effect of the US. Especially if the mice experience one material as more interesting than the other, a preference formation for one of the CS might be facilitated, as was the case here using flooring material which can be manipulated by the mice (e.g., gnawing, digging, see Supplements for observations during conditioning sessions). Thus, although flooring materials such as bedding material or gravel provide multiple useful cues, we would not recommend them for future experiments.

In experiment 3 and 4, metal plates were used as CS. Especially mesh or grid floors are frequently used as CS in other studies (mesh: Martínez et al. (1995); Oppong-Damoah et al. (2019), grid: Matsumura et al. (2010); Bahi, Nurulain, & Ojha (2014), mesh vs. grid: Fitchett, Barnard & Cassaday (2006)). The metal plates in our experiments contained a visual and a tactile cue (grid vs. holes, similar to the protocol of Cunningham, Gremel & Groblewski (2006)). As the metal was very bright, however, the visual cues did not have a high contrast. Moreover, the metal combined with the ethanol cleaning led to an unexpected additional cue: temperature. It has to be noted that in the protocol by Cunningham, Gremel & Groblewski (2006) with similar cues, no information is given on such a side effect. However, there is also no information given on recommended cleaning methods at all.

In the subsequent experiments visual cues as described in the protocol by Cunningham, Gremel & Groblewski (2006) (experiment 5) or similar to Lett et al. (2001) and Masaki & Nakajima (2008) (experiment 7 to 9) were used. The latter patterns were chosen especially because Wong & Brown (2007) showed that even mice up to 24 months of age are capable of distinguishing these patterns. In experiment 6, we used a new, self-designed set of patterns. We took special care that they would contain equal amounts of black and white squares without being too similar. However, it was a self-designed pattern without reference from any other study, and there was no distinct result in preference. Thus, we can not say if the mice were able to discriminate between the two patterns. If mice could not distinguish the cues, however, this might be a factor causing the resulting lack of preference (for cue or paired US compartment). Thus, using stimuli which cause no baseline preference is important -but it is equally important to ensure that the mice are able to distinguish (and memorize) the two patterns.

In general, the provided CS should match the abilities of the mice. In several of our experiments, some (experiment 5 and 6) or all mice (experiment 7 and 8) had reduced whiskers, and in these experiments, we took care to use visual cues instead of tactile ones (see also section Whisker-loss).

In addition, it is possible that mice perceived cues as important that were not intended. For example, a part of the experimental procedure itself might have been experienced as a more prominent CS which overshadowed other CS (for more see section Choice of US). Also the cleaning method might have worked as an olfactory cue (for more thoughts on that see section Cleaning).

### 4.2. Choice of US

As already mentioned, the here described experiments were preliminary. We aimed to develop a working protocol to use CPP for severity assessment, and therefore, we had to choose an US, here: an experimental procedure, with a predictable effect to validate our protocol.

However, some tested experimental procedures seemed not to be sufficient as a US with regard to successful conditioning and, thus, yielded no measurable effect. For example, in experiment 5, we used almond milk as a positive reinforcer. From other studies in our research group (Kahnau, Jaap, et al., (2023), also Habedank, Kahnau, & Lewejohann (2021)), we knew that mice prefer almond milk over other fluids and are also willing to work for the access to it. However, during conditioning sessions, not all mice tasted the fluid and only one mouse actually drank from it. This might have been related to insufficient habituation (see section Habituation), meaning that mice refrained from testing the fluid in an environment differing from their home cage. As a result, without experiencing the US, pairing of US and CS was probably unsuccessful. For this reason, we repeated this experiment with improved habituation and a different reward (millet) (experiment 6), as we knew from pre-tests that mice were more ready to consume millet in a new environment.

In addition, US and CS might influence each other. For example, in experiment 1 and 2, the US (fixation) contained a strong cue itself: the grid which was used to perform the fixation procedure. It is possible that the grid blocked or overshadowed any other cue (including the chosen CS). Blocking occurs if animals learn first the association between one stimulus and a consequence (e.g., grid and fixation), before this first stimulus is accompanied by a second stimulus (e.g., the chosen CS, in experiment 1 and 2: bedding material or gravel). The new stimulus (the chosen CS) will then be perceived as adding no new information and, therefore, conditioning for this new stimulus will be blocked (Shettleworth, 1998). Overshadowing, on the other hand, describes the effect of two simultaneously presented stimuli (e.g., grid and chosen CS) about each of which the animal learns less than if they had been presented alone. Thus, the conditioning response during the test in which only one stimulus is presented (e.g., flooring material as chosen CS) would turn out smaller (Shettleworth, 1998). In this case, it might be advisable to use no second CS but a cue that is already present during the procedure, e.g., use a surface for fixation that is then also present in the CPP setup.

It has also to be considered that the US we chose for our experiments might have been too weak in their effect, causing no distinct preference or aversion. As explained in the introduction, we refrained from using more severe experimental procedures as US as one of our aims for the CPP protocol was to detect also subtle changes in severity. Still, after the conduction of several experiments without the expected results, we have to consider that CPP experiments might not be suitable for the measurement of mild effects. In the review of Huston et al. (2013), the authors discuss what is actually conditioned in a CPP. One of the theories states that mice show some kind of “superstitious behaviour” after conditioning because they want to repeat the behaviour (i.e., being close to the respective CS) which lastly caused the positive reinforcement (Huston et al., 2013). Thus, using strong reinforcers might cause a stronger behavioural response. In our case, with a weaker reinforcer, the pull towards the CS might also be weaker, and as a result, might cause not the expected distinct differences in duration of stay in the two compartments. Possible alterations to potentially increase the effect of the US could be, e.g., prolonging the time the mice spent in the restrainer or conducting food deprivation before millet presentation. As CPP protocols using experimental procedures are hard to find, we can only refer to similar studies, which, however, also differ in their approach. For example, to measure the effect of wheel running with a CPP test, some studies used water deprivation (Lett et al., 2001; Masaki & Nakajima, 2008), while others did not (golden hamsters: Antoniadis et al. (2000)).

In general, it is also possible that mice might have learned an association between the US and some unintentional, unknown stimulus. This theory is based on behaviours by the mice which we observed especially during the first experiments: Mice were more hesitant to leave the handling tunnel right before the conduction of the (negative) US. This observation is described and discussed in the Supplements.

### 4.3. Choice of Setup

In the course of this study, we used three different setups. Setup 1 consisted of two cages connected by a tube with an automatic system detecting the position. As we used a similar connection system for the home cages, we expected the mice to readily accept this setup. However, both experiments using this setup (experiment 1 and 2), one mouse (the same for both experiments) changed cages during the baseline test but not during the preference test. It’s unclear why the mouse showed this behaviour, and if, for example, this was a sign of stress.

The next setup we tried, consisted of one cage separated into two halves by a small barrier (experiments 3 – 6). This setup was based on the setup used by Cunningham, Gremel & Groblewski (2006). The barrier was added a) to have a visual segregation of the compartments for the mice, and b) to facilitate the analysis (determining to which half of the cage the mice belonged). In this setup mice changed compartments more often. Indeed, one might consider the question whether the mice actually received the setup not as two compartments but as one large compartment (with an obstacle in the middle). This could imply that they did not experience the situation as providing a choice. In addition, without a visual barrier between the compartments, it can be argued that wherever the mice were staying, they still had (visual) contact with the other compartment: In experiment 3 and 4 we used tactile and visual cues (metal plates), while in experiment 5 and 6, using the same setup, we had visual cues only (patterns). As a result of conditioning, mice should seek out the CS which they associate with the preferred US (Huston et al., 2013). Nevertheless, in this setup, mice might not need to stay in one specific compartment to do so - because the cue (pattern) is visible from both compartments. This might not have been essential for other CPP studies, otherwise Cunningham, Gremel & Groblewski (2006) would probably not recommend this setup. Here, however, with a potentially weak US, the behavioural response to be as close as possible might be weaker. Following this line of thought, we reduced the (visual) contact with the CS of the second compartment in setup 3. This setup resembled the description of Masaki & Nakajima (2008) and Bahi, Nurulain, & Ojha (2014).

Note that setup 1 and 2 both had transparent walls, which enables mice to see external visual-spatial (room) cues. During the course of the experiments, care was taken not to change any external cues which were visible for the mice. In setup 1, external visual cues should not have played an important role: When using a one-compartment conditioning procedure, it was shown that visual-spatial cues do not lead to a CPP as they are only partially predictive (Cunningham, Patel, & Milner, 2006; Cunningham & Zerizef, 2014). In setup 2 with its two-compartment design, however, these external cues might also have had an influence. Interestingly, in 2 out of the 3 experiments in setup 3, there were side (compartment) preferences in the baseline and preference test. Setup 3 had all non-transparent walls and external visual-spatial cues were excluded. Thus, potential cues were restricted to the ceiling (more than 2 m above the setup), which contained a symmetric metal structure but no direct lights. It is possible that this structure was not completely symmetrically above the setup. Otherwise, it is also possible that the non-transparent walls caused a difference in lightning of the compartments, although we took special care to position the setup exactly between the light sources. If the lighting actually did cause the side preference, it also remains unclear, why in experiment 8, which was exactly positioned the way as experiment 7, this side preference was not observed.

As it can be seen in Table 1, many studies use setups with 3 compartments instead of 2. The advantage of a 3-compartment-setup is that the animal can be placed into a neutral area at the start of the test. The disadvantage, however, could be that the animal spends too much time in this neutral area throughout the test. As we wanted to develop a protocol, which would be suitable also to compare the effect of two aversive procedures (instead of comparing one of them with a neutral control, as is done in most studies), a neutral area would have resembled a refuge in which mice could refrain from making a choice. For this reason, we did not test a 3-compartment-setup.

### 4.4. Habituation

In the course of the experiments, we found habituation to be an important tool. As a first example: In pre-tests between experiment 5 and 6, we observed that a thorough habituation to the US environment was needed for a reward to be used as a US. Otherwise the mice did not consume the reward (millet or almond milk).

As a second example: In many experiments, we noticed a high defecation and urination rate. This is assumed to be a sign of distress or fear (Gray & McNaughton, 2003; Hurst & West, 2010). As this also happened during baseline tests, this seems to be setup and / or procedure related. Our aim was to find a protocol for CPP to measure the severity of specific experimental procedures. Thus, a protocol used for measurement (the CPP) which itself causes distress or fear should be reconsidered. Unfortunately, observations on urination and defecation during the procedure are not described in other studies so we cannot compare these observations. However, we assume that even if such behaviour was shown it is most often not reported in other studies.

In general, any sign of stress should not be underestimated as stress could influence conditioning: For example, acute restraint stress can influence behavioural flexibility such as reversal learning (Thai, Zhang, & Howland, 2012) and the retrieval of short-time and long-time memory (Li et al., 2012), which is mandatory for conditioning. The stress levels of our experiment should be relatively low compared to acute restraint stress, e.g., we had 1 min in a restrainer instead of several minutes up to an hour. Still, we do not know at which stress level the disturbance of memory processes starts. Habituation to the setup and the procedure could be an easy way to reduce stress during the actual CPP procedure.

In general, habituation (or repetition of the same procedure) improves the reliability of behavioural data (Rudeck et al., 2020) - in this case, this include the behaviour during baseline and final test as well as during conditioning sessions. In other words, the first time the mice are placed into the setup should perhaps not be the baseline recording. If mice are already familiar with the setup and the procedure, novelty induced variance might be reduced (Rudeck et al., 2020), resulting in more profound data.

However, for habituation in a CPP experiment, two things have to be considered:

First, the question arises whether it is possible to “over habituate” the mice. We would argue that this is not possible, as long as the US and CS stimuli remain unknown before (in the same manner as in the novel object recognition test, example protocol: (Lueptow, 2017). Instead we can assume that the more familiar the mice are with all other stimuli, the more attentive they should become towards a change.

Second, we can expect that habituation is most effective when setup and procedure (handling, timing, environment etc.) resemble the actual CPP protocol as much as possible. Of course, this does not mean that the to be tested CS or US should be familiar to the mice before the beginning of the CPP. Instead, the setup can be used in a similar procedure. For example, in experiment 9, we used plates with tactile structure and wall patterns as visual stimuli during the CPP as CS. During the habituation, we used similar plates but they were smooth, without additional tactile stimuli, and there were no patterns on the walls (i.e., grey walls instead of ones with black and white stripes). We had 4 days of habituation, in which also the procedure was mimicked, meaning we first placed the mice into the experimental environment for several minutes and then into the conditioning setup. This seemed to be quite efficient as in the baseline test (with the “real” tactile and visual stimuli), only 25% of mice urinated or defecated. In comparison: In experiment 4, we had habituation sessions, but they were short and not in the experimental room in which the CPP took place later on. Thus, we had not a different environment and also a differing procedure (no transport beforehand). In this experiment, 83% mice urinated or defecated during baseline recording.

### 4.5. Timing of US and CS

Regarding the presentation of US and CS, it has to be decided whether to present the CS before, simultaneously (both considered to be forward conditioning) or after the US (backward conditioning).

In experiment 1, we presented the stimuli before <u>and</u> afterwards with the argumentation that we wanted to be “sure” to get an effect. This is, however, misleading because presentation of the CS before or after the US can have opposite effects. For example, in a study with rats, Lett et al. (2001) presented one CS before the US (fluid, taste conditioning) and one CS after the US (pattern, place conditioning). They used wheel running as a US, which led to an aversion of the CS presented before the US (taste aversion) but a preference for the CS presented after the US (place preference; for more examples see Introduction). In the study, the authors do not give an explanation for this phenomenon, nevertheless, the opponent-process theory could provide a sufficient explanation. In short, a positive US, which has an attractive effect before its onset, can have an aversive effect after its presentation due to the removal of the positive stimulation (and the other way around for negative US, (Solomon, 1980)). Thus, if presenting the CS before the US leads to place preference, and presenting it afterwards to an aversion, the opposite effects might cancel each other out.

Forward conditioning is the more common procedure, whereby presenting first the CS and then the US (as we did in experiment 2 – 4) is not as common as presenting them simultaneously (see also Table 1). In experiment 5 and 6 we tested this simultaneous presentation of US and CS. However, especially in experiment 6, presentation of the US (in this case millet) did not work as expected: When placing the mice inside the conditioning cage, they often immediately started feeding and only later explored the cage (observations during conditioning are described in the Supplements). It is possible that the mice still perceived the cues during feeding. However, it might still have been <u>after</u> the perception of the odour of the millet. Thus, the odour might have blocked the patterns as a CS ((Shettleworth, 2009), also explained in section Choice of US).

To circumvent this, in experiment 7, millet was placed inside the cage with a delay. This approach was inspired by the study of Goltseker & Barak (2018), in which conditioned aversion was more prominent if mice experienced first the compartment with the CS and then the onset of the US (in their experiment: water flooding). However, as we only had a slight but not sufficient tendency after conditioning for a procedure preference, the timing might still not have been chosen sufficiently. We had 30 s until the US (millet or bedding) was placed in the compartment and 2.5 min afterwards for the mice to consume the millet (if present). Thus, mice were in contact with the CS for some time, while the US was already removed (eaten). It is possible that this time span was to long, and the “simultaneous“ presentation might not have worked properly.

For experiment 8 and 9, we used a different approach and changed to backwards conditioning. However, unlike Lett et al. (2000, 2001), Lett, Grant, & Koh (2002) and Belke & Wagner (2005) which also used backwards conditioning, we were not able to establish a conditioned place preference or aversion. The main differences between the mentioned experiments and ours are: First, our experiments were conducted with mice, not rats. It is possible that perception and learning differs between those species in ways that might be important here. Second, our procedure times were much shorter. The US (millet or restrainer) took only 1 min, and we confined the mice to the conditioning compartment for 3 min. In the study conducted of Lett et al. (2000), 2 h of wheel running were followed by 30 min in the conditioning compartment, Belke & Wagner (2005) had 2.5 h of wheel running with a fixed interval schedule followed by 30 min in the compartment, Lett et al. (2001) had 30 min of wheel running followed by 15 min in the compartment, and Lett, Grant, & Koh (2002) used 2 or 22 h of wheel running and 30 min in the compartment. All in all, these time frames are much longer than what we used. This might only be necessary for wheel running but not for other US. In addition, confining the animals for 30 min into a small chamber might in itself be stressful, not to mention the separation from the group. Especially, if we want to use conditioned place preference for severity assessment, it seems not feasible to use a protocol which itself might already influence the affective state of the animal.

### 4.6. Timing of the General Procedure

As already mentioned in the Introduction, few studies report details on the procedure not specifically related to the conditioning. For example: At what time of day does the conditioning exactly take place? What time difference lies between the first mouse and the last? Is there a transport to a different room? If one mouse is taken out of the home cage, how long before the other mice of its group are taken out, and thus, how long does the overall disruption last for the whole group?

We conducted our experiments during the light phase (as also described in the protocol by Cunningham, Gremel & Groblewski (2006)), starting closely after the lights went on. Pernold et al. (2019) found that C75BL/6J mice show an additional burst of activity after lights on. In addition, all groups of mice taking part in the experiments were used to the cage cleaning procedure once a week, which was also conducted in the same time frame (approximately hours 1 to 3 after lights on). Still, we observed that while the first mice (order was randomized) were always active, some of the last mice sometimes had to be woken up. This might have influenced their perception. Therefore, it would be very valuable to know if other studies had similar observations.

In all experiments, the participating mice lived together as one large group in a cage. With this, we can argue that we had no difference of treatment between cages (in comparison to other studies as we had only one), for example, with regard to the length of disruption due to the experiment (transport and stay in the experimental room, duration of removal of the filter top and light changes and so on). On the other hand, the overall disruption of the circadian rhythm might have been larger with this procedure because in total, the time the home cage conditions were affected by the experimental procedure was of course longer.

### 4.7. Transport

In the course of experiments, we learned that conducting the procedures in an experimental room (not the husbandry room) has to be considered carefully. First, habituation to the room conditions could never be as profound as for the husbandry room. Second, the transport itself includes several stimuli which might influence the perception of later presented stimuli. Thus, either the transport itself (and everything related to it) needs habituation to become an irrelevant stimulus, or the experiment should best take place in the room in which the animals are kept. Keeping the animals in the experimental room some days before the start of the experiment and during its conduction could be considered if possible.

### 4.8. Sound

Some studies, including the protocol by Cunningham, Gremel & Groblewski (2006), use sound attenuated chambers. Most studies, however, do not report whether they use them or not. For our experiments, no sound attenuated chambers were used. Thus, it is possible that sounds from other mice (in the home cage or other mouse groups kept in the same room) might have influenced the results. Experiments 6 – 9 were conducted in the husbandry room. In experiment 6 and 9, the group participating in the CPP experiment was the only group kept in this room. However, in experiment 7 and 8, a second group (group 2 from previous CPP experiments) was present and involved in a home cage based consumer demand test. This involved motor noises from an automatic door. However, we would argue that the mice should have been thoroughly habituated to these noises, and therefore, they should not have functioned as a new stimulus or a stimulus with relevant information on the procedure.

### 4.9. Whisker-loss

Barbering of fur and/or whiskers is a known, chronic problem in C57BL/6J mice (Kahnau et al., 2022). Whisker-loss can lead to altered behaviour, e.g., in the object recognition, marble burying and the open field test (Haridas et al., 2018; Tur & Belozertseva, 2018). In addition, the barbering mice are seen as a model for the disorder trichotillomania. However, it was shown that these mice show no reduced learning ability with the exception of a extra dimensional shift task (Garner et al., 2011), which is why they still should be suitable for conditioning.

In our experiments, only simple learning was required. Thus, in the sense of the 3R (Russell & Burch, 1959), we decided against ordering a new group of mice for the experiments (5 and 6). Still, we changed the previous experimental design from a tactile cue as CS to a visual one, as the whisker-loss should not have influenced the mice’s ability to perceive visual cues. In addition, we basically repeated experiment 8 with a fully whiskered group in experiment 9, to additionally compare the results of a whiskered and de-whiskered group. However, in both experiments, no procedure preference was found. This shows that the whisker-loss per se does not make the difference in the results.

Unfortunately, reporting levels of barbering are very low when it comes to studies unrelated to the investigation of this behaviour. It is therefore unknown if and how many study results are influenced by it. We here openly report that some of our mice groups showed whisker-loss. In these experiments, visual (not tactile) patterns were used as CS. We argue that in the studies mentioned above, in which altered behaviour was found, the tactile information was crucial. In our case, with our changed setup, on the other hand, mice should have been able to be conditioned even with missing tactile information.

### 4.10. Age

It has to be noted that mice took part in experiments until a rather old age of up to 19 months: In the sense of the 3R we wanted to “re-use” animals which had already participated in other mild behavioural experiments instead of ordering new ones, especially because we were conducting preliminary tests for a proof of concept.

In general, repeatability of activity measures increases with the age of mice (Brust, Schindler, & Lewejohann, 2015). It was shown that C57BL/6J mice performed well in visual detection, pattern discrimination and visual acuity tasks even until 24 months (Wong & Brown, 2007; Kahnau et al., 2021). Furthermore, as in the cited experiments their ability was tested using tasks relying on learning and memory, the conclusion can be drawn that these are also still intact. In addition, exploration, locomotor function and motivation of old mice seem to be initially similar (15 to 24 months versus 3 to 12 months), and only decline with prolongation of tasks (more than 10 min) (Jackson et al., 2021). However, as conditioning sessions as well as preference tests stay within this range, age-induced reduction should not influence the experiments per se. Moreover, operant and pavlovian conditioning learning as well as the motivation for appetitive reward are not impaired by age, when comparing C57BL/6 mice of 3, 6 and 15 months of age (Harb et al., 2014). Therefore, we argued that enrolment of older mice is feasible.

### 4.11. Cleaning

At least in the field of CPP studies, the cleaning procedure is usually not reported (examples: Cunningham, Gremel & Groblewski (2006), Sun et al. (2018), see also Table 1). Exceptions are Fitchett, Barnard & Cassaday (2006) (diluted detergent), Wang, Wang, & Chen (2014) (70% ethanol solution) and Carboni & Vacca (2003) (water and the soap used for cleaning of the home cages). Especially as, in contrast to our experiments, these published studies were successful in their conditioning, the missing information on cleaning states a problem.

In our experiments, we started with ethanol cleaning between the mice (experiments 1 – 5), arguing that ethanol erases potential olfactory cues from previous mice, and thereby, also potential influences from previous mice (Arakawa et al. (2008), rats: Wallace, Gorny, & Wishaw (2002)). However, the ethanol odour might have an influence on habituation and stress (see section Habituation). As a result, cleaning with ethanol might influence whether mice consume available millet. For this reason, in the experiments 6 – 9 (in which the consumption of the millet was an important part of the experiment), we refrained from disinfection between mice and just cleaned with water when necessary.

In addition, as already mentioned, we encountered unexpected effects of the ethanol cleaning procedure during experiment 3, in which the metal plates got notably cold due to the cleaning procedure. To prevent this, in experiment 4 we set a fixed time between the two procedures. However, there might still have been a difference in temperature between the plates. In this case, the temperature of the plates might interfere with the conditioning, as it was shown that different ambient temperatures can affect the formation of place preference (Dickinson & Cunningham, 1998). It would have been helpful to have recommendations from other studies, for example, Cunningham, Gremel & Groblewski (2006) which used similar plates, how they handled this cleaning-temperature-problem.

### 4.12. Conclusion

We performed 9 CPP experiments in search for a conditioning protocol which would enable us to compare different experimental procedures with regard to their severity. However, none of the tested protocols resulted in a distinct preference for one of the procedures. Even simpler procedure protocols using a food reward as a treatment (similar to Takeda et al. (2001); Matsumura et al. (2010)), failed to do so. We propose two possible explanations:

First, the effect of the tested US might not be strong enough to result in conditioning. As a consequence, an US with a stronger emotional effect (be it attractive or aversive) could yield more distinct results. However, in this study we wanted to find a protocol which would also be efficient in comparing mild severity. Using “stronger” US would, therefore, have gone beyond the scope of the addressed question.

Second, mice might not have been able to associate the CS as the important cue with the tested procedure (US). This might be related to the conduction of the CPP itself, involving everything from handling to setup, choice of CS and CS–US timing. Potential pitfalls in those factors were already discussed above. Moreover, the following question might arise: Does the experimental procedure (and thus, the US itself) maybe involve too much own stimuli, which overshadow or block the presented CS? In this case, it could be more effective to use stimuli already present in the experimental procedure (inspired by Fitchett, Barnard & Cassaday (2006)). Note that the existing studies pairing a CS with wheel running involved rats (Lett et al., 2000, 2001; Lett, Grant, & Koh, 2002) and hamsters (Antoniadis et al., 2000), which might perceive the conditioning procedure different from mice.

Conducting the experimental <u>in</u> the CPP setup, as for example done in the study of Martínez et al. (1995), which compared male encounter (aggression) to no encounter, would not be possible for most experimental procedures. However, in experiment 1 and 2, we used a grid to restrain the mice by hand, and a glass jar for weighing. Using the grid and the glass as flooring structures in the final test could lead to a more pronounced preference than we got in our experiments. In this case, special care would have to be taken to exclude baseline preference (as in experiment 4 regarding the procedure’s surfaces) and randomisation of CS–US pairing (one half with CS1 and US1, and one half with CS2 and US1). In addition, finding the best CS as part of the experimental procedure might require pre-studies. For example, comparing Water Maze and Barnes Maze, the obvious stimulus would probably be water. But the mice might have an aversion against water (compared to no water) even before conditioning. Therefore, it might be more advisable to use, e.g., different colours for the walls of the mazes, which then, in return, could also be less relevant for the mice during the procedure. Thus, pairing would again become more difficult. As this would lead to a complete new set of experiments, we decided to draw up an intermediate state with this article.

To conclude, we still believe that finding a conditioning protocol to compare severity of different experimental procedures should be possible. The here reported experiments provide helpful information on the research so far and can hopefully function as the basis for subsequent developments of conditioning protocols. All in all, our study shows that CPP is probably more of a test that is specialized for specific questions and research conditions. In the area of severity assessment, we suggest relying on other testing methods in the future. Methods of direct or automated behavioural observation in the home cage have proven to be promising in this regard (Kahnau, Mieske, et al., 2023).

## 5. Additional Method Details

### 5.1. Conditioned Stimuli

#### 5.1.1. Flooring Material (Tactile, Visual and Odour Cues)

Experiment 1:

As CS, bedding material was used, similar to the CPP experiments by Fitchett, Barnard & Cassaday (2006) and Meisel & Joppa (1994). We used “pure” and “comfort white” bedding material (JRS, J. Rettenmaier & Söhne GmbH + Co KG, Germany). Although they both consist of cellulose, they are distinguishable in size, texture and, even for a human, in odour (for a picture see Supplements).

Experiment 2:

As a modification of experiment 1, different types of gravel were used (obtained in a local DIY store). We intended to use pumice and marble but as mice showed a clear preference for pumice during a baseline test (probably due to its differing thermal characteristics), we used marble and quartz instead. The gravel was thoroughly washed and disinfected through autoclaving before using it for the experiment. Marble and quartz differed in colour and shape (for a picture see Supplements). Thus, they included visual and tactile cues.

#### 5.1.2. Metal Plates (Tactile and Visual Cues)

Experiments 3 and 4:

In both experiments, plates were used, referring to the studies of Cunningham, Gremel & Groblewski (2006) and Cunningham, Patel, & Milner (2006) (only tactile stimuli, no additionally coloured walls). Here, we used a metal plate with holes and a metal plate with slits (for a picture see Supplements). Plates were obtained in a local DIY store and cut to 180 x 210 mm to fit into one half of a type III cage. Both types of plates consisted of aluminium. Thus, flooring materials included a visual (dots vs. stripes) and a tactile cue (holes vs. slits).

In addition, all plates might have thermal cues, amplified by cleaning with ethanol (for more details, see Supplements).

#### 5.1.3. Patterns (Visual Cues)

Experiments 5:

A laminated paper with a patterns was placed directly under each cage half (setup 2). The patterns were designed after the description of Cunningham, Gremel & Groblewski (2006): One pattern consisted of black circles (diameter: 6.4 mm), each row shifted slightly in its center to the one before. The other pattern consists of wide black lines (widths: 3.2 mm, space between edges: 6.4 mm) (stripes).

Experiments 6:

Self-designed patterns were used, consisting of black and white squares. Care was taken that the patterns contained the same amount of black and white space, but differed in block size and alignment of blocks (shifted vs. linear). This resulted in one pattern similar to a chessboard and one similar to fabric texture (for a picture see Supplements).

Experiments 7, 8 and 9:

Patterns were either horizontal or vertical black and white stripes (similar to Wong & Brown (2007)).

#### 5.1.4. Plastic Plates (Tactile and Visual Cues)

Experiments 9:

Tactile cues in addition to visual cues (referring to studies of Cunningham, Patel, & Milner (2006) and Cunningham & Zerizef (2014)) were used. This was done by adding 3D printed plastic plates (material: PLA) onto the floor of the procedure environment and the conditioning setup (for a picture see Supplements).

Plates for the procedure environment (conditioning session) were white, 2 mm high and contained protruding tactile cues: either bars or dots. Plates for the conditioning setup were black, 2 mm high and contained holed tactile cues: either squares (diagonally organised) or rounded crosses (in parallel), similar to the patterns described by Mei et al. (2020). Plates for the habituation beforehand (procedure environment and test setup) were smooth without any additional tactile cues.

### 5.2. Unconditioned Stimuli

#### 5.2.1. Fixation (Restraint by Hand)

Experiments 1 and 2:

The mouse was taken out of the conditioning compartment and placed on a lid on top of a cage. If the mouse did not leave the handling tunnel voluntarily, it was tilted until the mouse gently slid out. While holding the tail with one hand, the animals got restraint by taking the loose skin of the scruff between thumb and index finger of the other hand and lifting the mouse off the lid (as described, e.g., in Hurst & West (2010)). The mouse was held for 20 s, before it got released straight into the conditioning compartment (experiment 1) or back onto the surface (experiment 2). Afterwards, the mouse was returned to the transportation cage.

Experiment 3:

The mouse was taken out of the conditioning compartment and placed on the surface (on top of an upside down cage, type 1144B, LWH: 331 x 159 x 132 mm, Tecniplast). The back opening of the handling tunnel was sealed by hand and it was waited until the mice left the tunnel by itself. While holding the tail with one hand, the animals got restraint by taking the loose skin of the scruff between thumb and index finger of the other hand and lifting the mouse off the surface. The mouse was held for 20 s, before it got released back onto the surface. Afterwards, the mouse was returned into the transportation cage.

#### 5.2.2. Weighing

Experiments 1, 2, 3 and 4:

The mouse was taken out of the conditioning compartment and placed into the weighing vessel (experiment 1, 2, 4: glass jar; experiment 3: cage type 1144B, LWH: 331 x 159 x 132 mm, Tecniplast) on top of a scale. In case of the glass jar (experiment 1, 2, 4), a lid was placed on top to prevent the mouse from climbing out. After weight had been noted, the mouse was placed into to conditioning compartment (experiment 1) or to the transportation cage (experiment 2 and 4).

#### 5.2.3. Millet in Separate Cage

Experiment 4:

The mouse was taken out of the conditioning compartment and placed in a type III cage (LWH: 425 x 276 x 153 mm, Tecniplast) filled with bedding material (the same as in the home cage). The cage contained at one end a 0.1 g millet (approximately 16 grains; Goldhirse, Spielberger Mühle, Germany) as food reward. The mouse was taken out of the cage immediately and returned to the transportation cage, a) after it stopped feeding (independent from the amount eaten) or b) if the mouse did not start feeding: after 1 min. Feeding was defined as sitting beside the millet for some time and eating the grains audibly; feeding was only counted as such if more than 3 grains were consumed.

Mice were habituated to the millet before the experiment by offering it for three days in the morning, three times 2 g in different places in the home cage.

Experiment 8:

The mouse was taken out of the home cage and placed in a type III cage (LWH: 425 x 276 x 153 mm, Tecniplast) filled with bedding material (the same as in the home cage). Millet (0.1 g) was added at one end of the cage directly after placing the mouse in the cage. The mouse was taken out of the cage and placed into the conditioning compartment, a) when the mouse stopped feeding on the millet and no remaining grains were visible, or b) at the latest after 1 min.

Mice were familiar with millet from previous experiments.

Experiment 9:

The mouse was taken out of the home cage and placed into a type III cage (LWH: 425 x 276 x 153 mm, Tecniplast) which had a 3D printed white plate with tactile stimuli on its floor. The mouse had 5 s to inspect the cage (and its stimuli), before 0.1 g of millet were placed into the cage, always at the same spot. After 1 min, the mouse was placed into the conditioning compartment (setup 3).

Mice were familiar with millet from active enrichment in their home cage.

#### 5.2.4. Millet and Bedding

Experiments 4 and 8:

see section Millet in Separate Cage

Experiment 6:

For this procedure, millet and bedding material (the same used in the home cage) were used. In one treatment, the bedding material was presented alone, in the other, it was mixed with millet. Mice were familiar with millet from previous experiments and also habituated to feeding on millet outside their home cage environment.

The procedure was performed in the conditioning compartment itself (setup 2). In the middle of the compartment, a small amount of bedding material (as much as fitted into a 0.5 ml Eppendorf tube) was placed, which was either mixed with 0.1 g millet or not. After placing the mice inside the compartment, the compartment was covered by a Perspex plate.

Experiment 7:

For this procedure, millet and bedding material (the same used in the home cage) were used (unmixed). Mice were unfamiliar with millet and <u>not</u> habituated to feeding on millet outside their home cage environment but were thoroughly habituated to the conditioning compartment (without the CS) so we expected mice to immediately feed on the millet.

The procedure was performed in the conditioning compartment itself (setup 3). After the mouse was placed inside the conditioning compartment, it was waited 30 s before the treatment was applied. The treatment was either 0.1 g millet or a visually similar amount of bedding material. The compartment was not covered with a plate.

#### 5.2.5. Restrainer

Experiment 8:

The mouse was taken out of the home cage and placed into a type III cage (LWH: 425 x 276 x 153 mm, Tecniplast) filled with bedding material (the same as in the home cage). It was then guided or pushed by hand into the tunnel of the custom-built restrainer. The restrainer had to barriers which reduced the inner space to a different volume (detailed description and picture of the device see Supplements). The first, outer restrainer barrier was inserted and the timer was started. If possible by the size of the mouse, the second, inner barrier was then inserted, to further restrict the space. After 1 min, the mouse was directly released into the conditioning compartment (setup 3).

Experiment 9:

The same custom-built restrainer as in experiment 8 was used. The mouse was taken out of the home cage and (similar to millet, experiment 9) placed into a type III cage (LWH: 425 x 276 x 153 mm, Tecniplast) which had a 3D printed white plate with tactile stimuli on its floor. The mouse had 5 s to inspect the cage (and its stimuli) and was then guided or pushed by hand into the tunnel of the restrainer. Barriers were inserted in the same manner as in experiment 8, and after 1 min, the mouse was directly released into the conditioning compartment (setup 3).

#### 5.2.6. Fluids

Experiment 5:

As US diluted almond milk and tap water were used: The almond milk was prepared using 1 part “Mandel Drink” (containing water, 7 % almonds, and salt, Alnatura GmbH, Germany) and 3 parts tap water from the husbandry room. The diluted almond milk was prepared in the morning right before the first conditioning session and kept in a fridge from thereon for the next days, so only the amount which would be used this day was taken into the experimental room. Mice were unfamiliar with almond milk.

This procedure was performed in the conditioning compartment (setup 2). The compartment was covered by a perspex plate, and access to the respective fluid was given through a bottle inserted into a hole in the covering plate.

During the first day of conditioning (session 1 and 2), bottles were filled with 200 ml fluids and it was noted that bottles trickled a lot, leading not only to drops on the floor but to a puddle with approximately 5 cm diameter. Because mice hesitated to step into the puddle, the bottle directly above became too far away for some mice to reach it. To improve this situation, for the second day of conditioning, the plate for bottle insertion was altered, so that it hang lower (about 2 cm lower than before). In addition, bottles were filled with 600 ml of fluid, which reduced the trickling (although it did not stop, see picture in Supplements).

## 6. Ethical Approval

All experiments were approved by the Berlin state authority, Landesamt für Gesundheit und Soziales, under license No. G 0182/17 and were in accordance with the German Animal Protection Law (TierSchG, TierSchVersV).

Experiments were preregistered at the Animal Study Registry:

DOI: 10.17590/asr.0000112 (experiment 1, experiment 2 as comment)

DOI: 10.17590/asr.0000142 (experiment 3, experiment 4 as comment)

DOI: 10.17590/asr.0000161 (experiment 4.1, no CPP, see Supplements)

DOI: 10.17590/asr.0000191 (experiment 4.2, no CPP, see Supplements)

DOI: 10.17590/asr.0000208 (experiment 5)

DOI: 10.17590/asr.0000215 (experiment 6)

DOI: 10.17590/asr.0000227 (experiment 7)

DOI: 10.17590/asr.0000231 (experiment 8)

DOI: 10.17590/asr.0000236 (experiment 8.1, no CPP, see Supplements)

DOI: 10.17590/asr.0000260 (experiment 9)

## Acknowledgments

The authors thank the animal caretakers, especially Carola Schwarck, for their support in the animal husbandry, and Birk Urmersbach for his support with the automated video analysis in experiment 4.2 (no CPP experiment, see supplements).

## 10. Author contributions

Anne Jaap: Conceptualization, Data Curation, Formal Analysis, Investigation, Methodology, Project Administration, Resources, Validation, Visualization, Writing – Original Draft Preparation

Pia Kanau: Conceptualization, Methodology, Resources, Writing – Review & Editing

Lars Lewejohann: Conceptualization, Funding Acquisition, Project Administration, Resources, Supervision, Writing – Review & Editing

## 11. Supplementary Material

For supplementary material, e.g., regarding three additional, non-CPP experiments see: Supplements.pdf.

The raw data of the experiments can be found under: https://doi.org/10.5281/zenodo.10877368

