## Supplementary Material for "In search of a conditioned place preference test for mice to assess the severity of experimental procedures"

Anne Jaap[1*], Pia Kahnau[2] & Lars Lewejohann[1,2]

[1] Institute of Animal Welfare, Animal Behavior and Laboratory Animal Science, Freie Universität Berlin, Königsweg 67, D-14163 Berlin, Germany
[2] German Federal Institute for Risk Assessment (BfR), German Center for the Protection of Laboratory Animals (Bf3R), Max-Dohrn-Straße 8-10, D-10589 Berlin, Germany
*

**6 Discussion: Latency to Leave Tunnel** **17**

**7 Experiment 4.1 (Choice Test)** **18**

### 1 Introduction

In the following, short additional information is given on the Conditioned Place Preference experiments (CPP experiments, see main paper) and on three experiments with alternative approaches.
To match the structure of the main paper, this supplementary material will also start with information on conditioned stimuli and unconditioned stimuli, followed by details on cleaning procedures. Then, additional observations of the CPP experiments are reported. Additional information on the CPP experiments of the main paper include:

- details on the conditioned stimuli used in
  experiment 1 and 2 (flooring material, see section 2.1),
  experiment 3 and 4 (metal plates, see section 2.2),
  experiment 6 (patterns, see section 2.3),
  and experiment 9 (plastic plates, see section 2.4);

- details on unconditioned stimuli of
  experiment 5 (presentation of fluids, see section 3.7)
  and experiment 8 (restrainer, see section 3.3.2);

- details on cleaning of the three setups (see section 4.1), Conditioned Stimuli (see section 4.2), including observation on temperature change of metal plates) and Unconditioned Stimuli (see section 4.3)

- general observations during the conditioning, including the latency to leave the handling tunnel if recorded (all CPP experiments, see section 5);

- discussion of the latency to leave the handling tunnel (see section 6).

In the end, three additional non-CPP experiments are described. They were conducted between CPP 4 and 5 (called 4.1 and 4.2), and between CPP 8 and 9 (called 8.1).

### 2 Additional Information on Conditioned Stimuli

#### 2.1 Flooring Material

##### 2.1.1 Experiments 1 and 2

In experiment 1 and 2, different types of flooring material were used (combining tactile, visual and potentially also olfactory cues, Fig. 1).

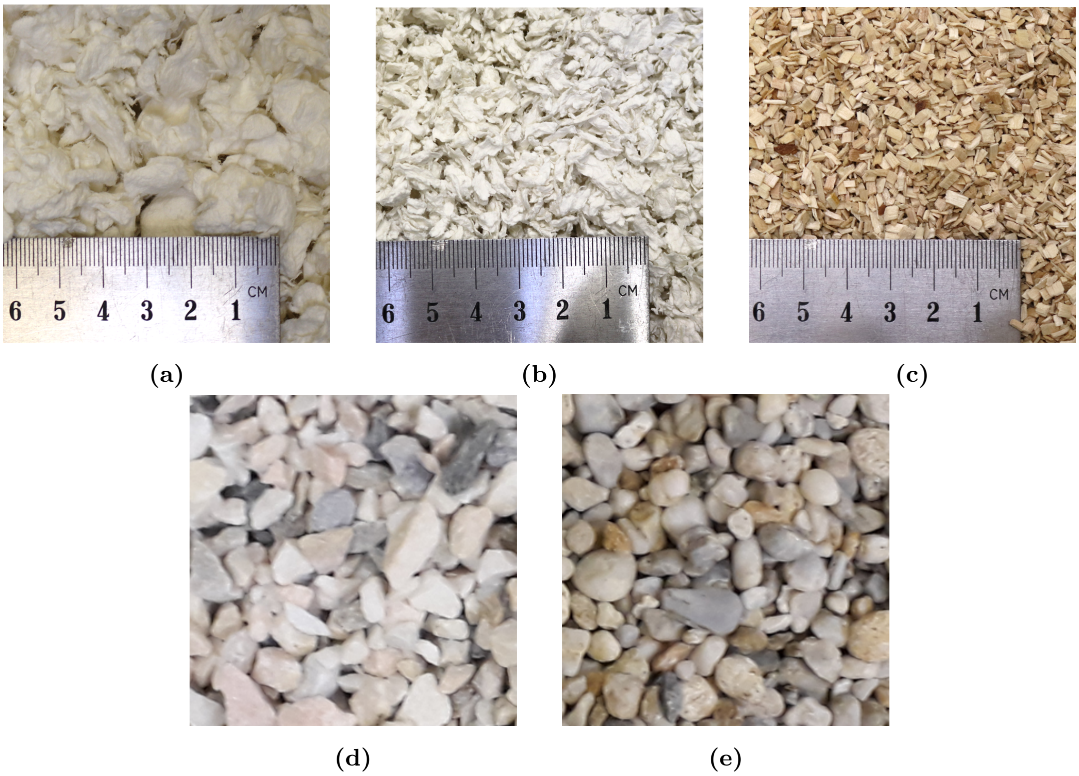

**Figure 1:** Flooring materials used as conditioned stimuli. Comfort White (a) and Pure (b) bedding material were used in experiment 1 and consist of cellulose. (c) The Poplar Granulate bedding material consists of poplar chips. This bedding material was not used in the conditioned place preference test but was used during normal husbandry conditions. Marble (d) and quartz (e) were used in experiment 2.

#### 2.2 Metal Plates

##### 2.2.1 Experiments 3, 4 and 4.1

In experiment 3, 4 and 4.1 (no CPP experiment), plates were used, referring to the studies of Cunningham et al. 2006a, Cunningham et al. 2006b (only tactile stimuli, no additionally colored walls, see Fig. 2). In experiment 3 and 4, plates with holes and slits were used, both types consisting of aluminium. In experiment 4.1, we used metal grid and a metal plate with larger holes, both plates consisting of steel. All plates were obtained in a local DIY store and cut to 180 x 210 mm to fit into one half of a type III cage. Thus, flooring materials included a visual (dots vs. stripes) and a tactile cue (holes vs. slits). In addition, plates were placed in setup 2, which provided a spatial cue (left vs. right side of the conditioning cage. However, all plates might also have contained thermal cues.

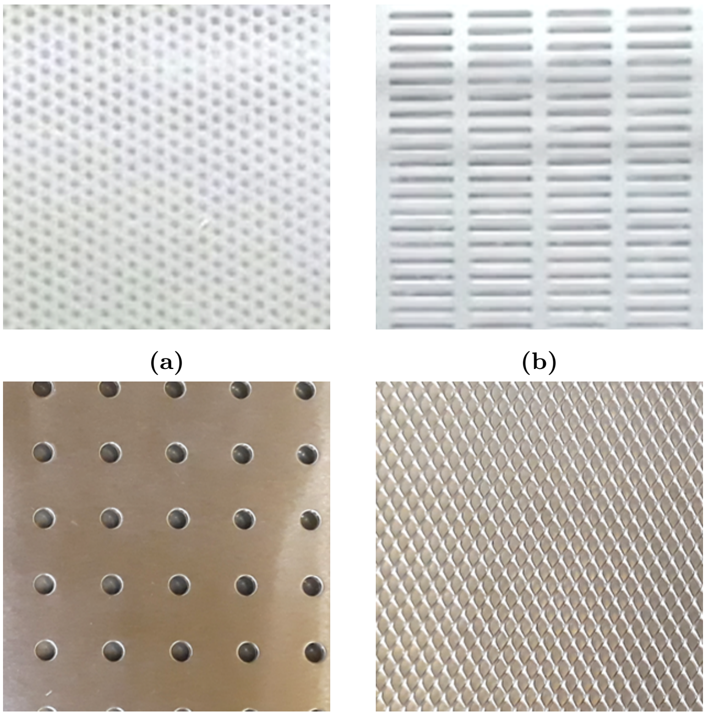

**(a)** **(b)** **(c)** **(d)**

**Figure 2:** Metal plates used as conditioned stimuli. A plate with small holes (a) and a plate with slits (b) were used in experiment 3 and 4. A plate with fewer but larger holes (c) and a grid (d) were used in experiment 4.1 (no CPP experiment).

#### 2.3 Patterns

##### 2.3.1 Experiment 6

Although already described in the main paper, we provide here additional pictures (Fig. 3) to show how the fabric-texture like pattern of experiment 6 looked like.

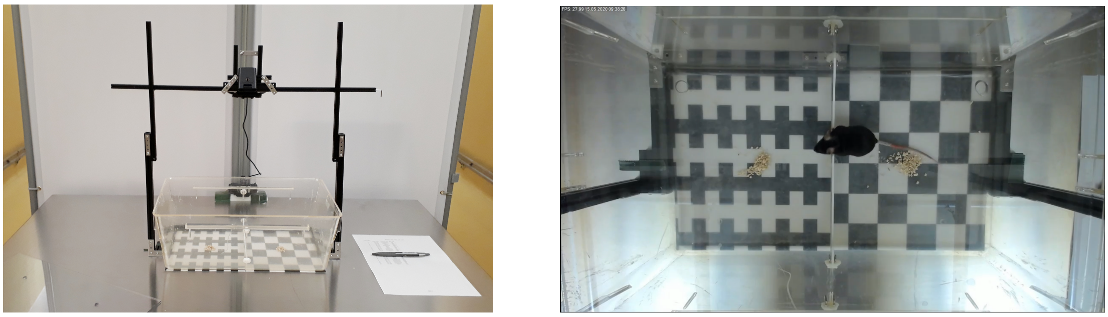

**(a)** General setup **(b)** View from above

**Figure 3:** General setup and patterns used in experiment 6.

##### 2.3.2 Experiment 8.1

In addition to the patterns used in experiment 5 – 9 (see main paper), in experiment 8.1 (no CPP experiment), we used the patterns from experiment 5 again but on the walls, leading to dots and horizontal stripes. The patterns were placed on all 4 walls (setup 2).

#### 2.4 Plastic Plates

##### 2.4.1 Experiments 8.1 and 9

In experiment 8.1 (no CPP experiment) and 9, we used tactile cues in addition to visual cues (referring to studies of Cunningham and Zerizef 2014, Cunningham et al. 2006b). This was done by adding 3D printed plastic plates (PLA) onto the floor of the procedure environment (experiment 8.1 and 9) and the conditioning setup (experiment 9). Plates were black and contained holed tactile cues: either holes or slits. To increase contrast, white paper sheets were placed underneath the opaque cage floor. The plates are depicted in Fig. 4.

### 3 Additional Information on Unconditioned Stimuli

#### 3.1 Weighing

##### 3.1.1 Experiment 4.1

Before each mouse, the weighing vessel (here a glass jar) was cleaned with 70 % ethanol in order to eliminate any olfactory cues by previously tested mice.
The to be tested mouse was taken out of the conditioning compartment by tunnel handling and placed into the weighing vessel on top of a scale. A lid was placed on top to prevent the mouse from climbing out. After weight had been noted, the mouse was taken out of the weighing vessel by tunnel handling and placed into to conditioning compartment.

#### 3.2 Millet in Separate Cage

##### 3.2.1 Experiment 8.1

In experiment 8.1 (no CPP experiment), the mice were taken out of the home cage by tunnel handling, and placed into a type III cage (LWH: 425 x 276 x 153 mm, Tecniplast, Italy) with patterns on the walls and tactile and visual patterns on the floor. After 15 s, 0.1 g millet were added (always to the same spot) and mice had 1 min access to it. Afterwards, they were taken out by tunnel handling and placed back into their home cage. Between mice, the cage and its floor plate were cleaned with 70 % ethanol. Mice were familiar with millet from active enrichment in their home cage.

#### 3.3 Restrainer

##### 3.3.1 Experiment 4.1

The to be tested mouse was taken out of the conditioning cage by tunnel handling. If the tail of the mouse was inside the tunnel and not graspable, a smaller sealed tunnel was used to push the mouse gently backwards until the tail was available. The mouse was then pulled carefully

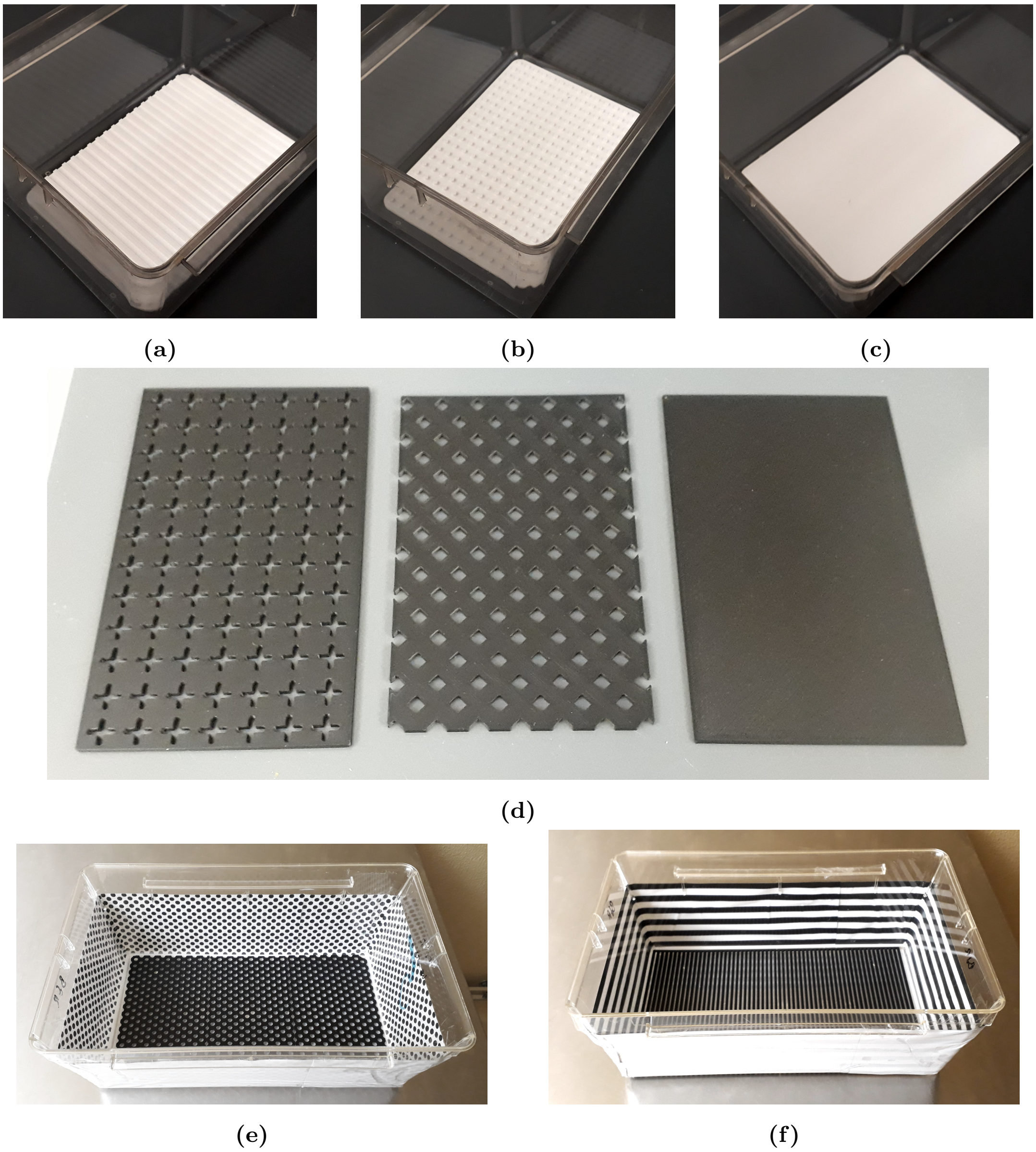

**Figure 4:** Plastic plates used as conditioned stimuli. A plate with bars (a) and a plate with dots (b) were used in experiment 9 in the procedure environment. A plate without additional tactile cues was used for habituation (c). In the conditioning setup, a with holed crosses (d, left) or a plate with holed squares (d, middle) were used, while a plate without additional tactile cues was used for habituation (d, right). Note that the plates have differing length because we used to of them for one compartment and they had to be split for 3D printing at a suitable line. (e) and (f) are the procedure environments used for experiment 8.1 (no CPP experiment), containing plates with holed dots or stripes.

backwards by its tail into the restrainer (LWH: 174 x 100 x 67 mm, Roth, Karlsruhe, Germany) until it reached the correct position (the restrainer was designed to facilitate blood sampling or an injection). A small barrier including a slit for the nose was added in front of the mouse. The mouse had to stay in the restrainer for 1 min, before the barrier was taken away and the handling tunnel was offered to the mouse for re-entering. Between mice, the restrainer and the smaller tunnel were cleaned with 70 % ethanol.

##### 3.3.2 Experiment 8

In Fig. 5 the custom-built restrainer used in experiment 8 is depicted.

##### 3.3.3 Experiment 8.1

In experiment 8.1 (no CPP experiment), we used the same custom-built restrainer as in experiment 8. The mice were taken out of the home cage by tunnel handling, and placed into a type III cage (LWH: 425 x 276 x 153 mm, Tecniplast, Italy) with patterns on the walls and tactile and visual patterns on the floor. After 15, the restrainer was placed inside the experimental cage and the mice were guided by hand into the restrainer. Immediately, the first (outer) barrier was inserted and the timer was started. Then the second (inner) barrier was then inserted, to further restrict the space. During the whole restrainer procedure, the restrainer was located inside the experimental cage, so the mouse could still see the patterns of the cage. After 1 min, the barriers were removed and the mouse could leave the restrainer back into the experimental cage. After the procedure, the mouse was taken out of the experimental cage by tunnel handling and returned to the home cage.
Between mice, cage and the floor plates were cleaned with 70 % ethanol. The restrainer (if used) was cleaned on day 2 (session 1) with 70 % ethanol, but because this destroyed the tunnel and it had to be replaced. From day 3 on, the restrainer was then cleaned with tissues free from alcohol and aldehydes (Microbac Tissues, Hartmann, Germany).

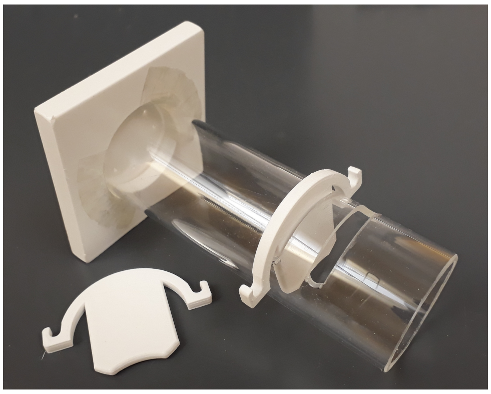

**Figure 5:** Self-build restrainer as used in experiment 8 and 8.1. The tunnel has 4 cm diameter and a length of 12 cm. Barriers are inserted at 7.5 and 9.0 cm.

#### 3.4 Open Field

##### 3.4.1 Experiment 4.2

The open field apparatus (rectangular arena 75 x 75 cm, with 57 cm high grey walls) was approximately evenly lit with 73 to 98 lux by two lights directly above the apparatus, ensuring an illumination without blinding or reflection (see Fig. 6). Because lights were initially too bright, paper sheets were attached in front of them for dimming. A metal construction (first out of beams by fischertechnik GmbH, Germany) had a WebCam (Logitech C390e WebCam, Switzerland) applied to it, so that the apparatus could be filmed from above.
The apparatus was cleaned with 70 % ethanol and dried before testing to eliminate any olfactory

cues by previously tested mice. For each trial, the mouse was taken out of the experimental cage by tunnel handling. The tunnel was placed in front of a small hole which was in one corner (4 cm diameter) through which the mouse then entered the open field. For the first minute, the mouse had only access to a smaller part of the open field directly in-front of the hole (15 x 15 cm) for acclimatization. We used a construction of two grey plastic plates for that. After 1 min, the walls were lifted and the mouse was free to explore the maze for 5 min, during which its behaviour was video recorded (using a Logitech C390e WebCam, Switzerland). Then the mouse was returned to the conditioning compartment by tunnel handling.

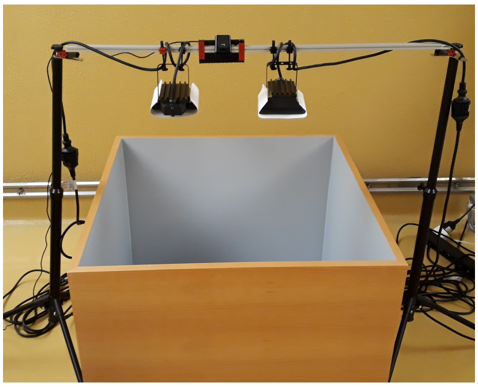

**(a)**

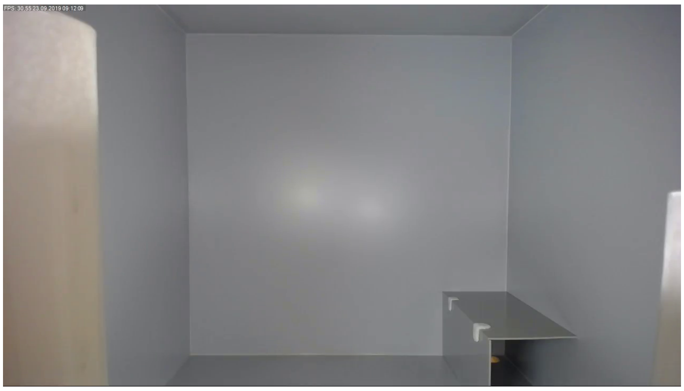

**(b)**

**Figure 6:** Setup of the open field as used in experiment 4.2 (no CPP experiment). The open field consists of a large box, which is equally illuminated and filmed from above. In the camera view (b) the rectangular walls restricting the mouse to a smaller part of the open field for acclimatization is visible. The white parts in the picture are paper sheets in front of the two lights for dimming.

#### 3.5 Home Cage

##### 3.5.1 Experiment 4.2

The mouse was taken out of the conditioning compartment by tunnel handling and placed back into the home cage for 3 min. Afterwards, the mouse was returned to the conditioning cage.

#### 3.6 Separation

##### 3.6.1 Experiment 4.2

For each mouse, a type II cage (LWH: 225 x 167 x 140 mm, Tecniplast, Italy) was used, containing two sheets of paper to cover the floor. Each mouse got its own cage to prevent any olfactory cues from previously tested mice. Cages were cleaned with 70 % ethanol between the experimental days.
For the procedure, a mouse was taken out of the conditioning compartment and placed into the type II cage. A filter top but no grid was placed on top of the cage. The mouse was left inside the cage for 10 min. Afterwards, the mouse was returned to the experimental cage by tunnel handling.

##### 3.7 Fluids

###### 3.7.1 Experiment 5

In Fig. 7 the presentation of fluids of experiment 5 is depicted.

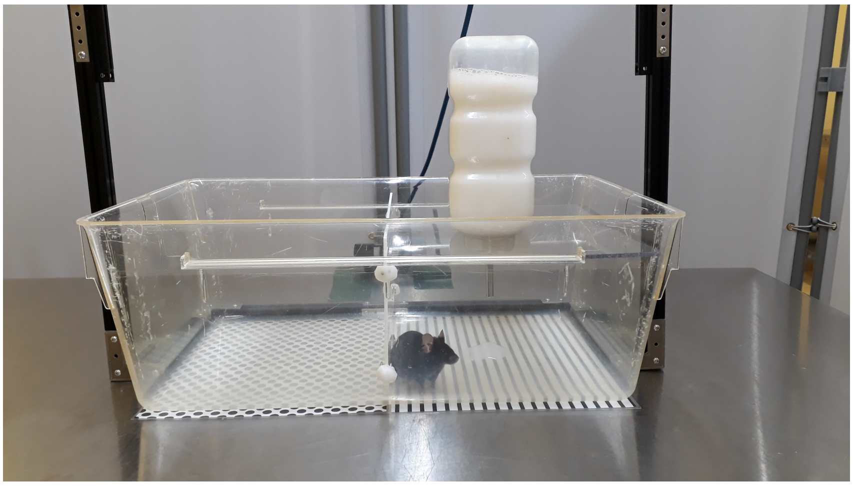

**Figure 7:** Presentation of the fluids. This is the already altered setup with a lower hanging bottle and greater filling height. Almond milk dripped stronger than tab water.

### 4 Cleaning

#### 4.1 Setups

##### 4.1.1 Setup 1: Two cages connected via a tunnel

In this setup, bedding material or different kinds of gravel were used as CS. Between mice, the CS were removed and cage, lid, and tunnel cleaned with 70 % ethanol.
For conditioning sessions, each mouse had its own conditioning cage to facilitate the procedure (no cleaning between mice). It was emptied, cleaned with 70 % ethanol and re-filled with the CS (bedding material or gravel) between the experimental days. In addition, between mice, the weighing and fixation surface were cleaned with 70 % ethanol.

##### 4.1.2 Setup 2: One cage separated by a barrier

In this setup, either different metal plates (experiment 3 and 4) or visual patterns (experiment 5 and 6) were used as CS, in both cases positioned on the floor only. As we had only one setup, in conditioning sessions, habituation, and the final test, the setup was cleaned between mice: In experiment 3 and 4, the setup and its barrier/plate was cleaned with 70 % ethanol between mice. The floor plates were first cleaned with water, dried with paper, and then cleaned with 70 % ethanol and dried with paper again. In experiment 5, between mice the setup was first cleaned with water (to wash away the liquid and general soiling) and then with 70 % ethanol. In experiment 6, between mice the setup was only cleaned with water if urination or defecation occurred. This was done to reduce potential odour effects of the ethanol itself (see also Discussion). In all experiments, the setup was cleaned with 70 % ethanol between experimental days.

###### 4.1.3 Setup 3: Two compartments separated by a wall

The setup was cleaned with 70 % ethanol before or after each experimental (and habituation) day. Between mice, the setup was only cleaned if faeces or urine were present, and cleaning was done just with water. This was done to reduce potential odour effects of the ethanol itself (see also Discussion).

##### 4.2 Conditioned Stimuli

###### 4.2.1 Metal Plates

**Experiment 3:** In experiment 3, it was noted while disinfecting the plates with ethanol in-between the mice that the metal went very cold from evaporation. Thus, depending on the time between cleaning and getting the next mouse into the conditioning cage, the “coldness” of the plate might have differed for every session and every mouse. Also, this “coldness” might have been different for the two plates due to their recess structures (many small holes or large long slits). This might have the effect of an additional cue.
To specify this effect, we took measurements with an infrared thermography camera (PIR uc 180 camera with IRBIS software, Version 3.1.82, both: InfraTec, Germany) after the end of experiment 3 using a infra red camera. We did not observe a temperature difference between the plates themselves. However, there was a temperature difference between the measurement directly after the cleaning vs. 90 s after cleaning (directly after cleaning: minimum temperature about 17.0 °C, area average about 19.0 °C; after 90 s: minimum temperature about 20.0 °C, area average about 20.5 °C).

**Experiment 4:** As we noted in experiment 3 that the metal plates used as CS got noticeably cold directly after cleaning with 70 % ethanol, in this experiment we waited at least 90 s after the cleaning before a mouse was placed onto the plates.

##### 4.3 Unconditioned Stimuli

###### 4.3.1 Fixation (Restraint by Hand)

**Experiment 1 and 2:** Before each mouse, the cage lid and the cage, on which the lid was placed, were cleaned with 70 % ethanol in order to eliminate any olfactory cues by previously tested mice.

**Experiment 3:** Before each mouse, the surface on which the mice were placed (on top of an upside down cage, type 1144B, LWH: 331 x 159 x 132 mm, Tecniplast, Italy) was cleaned with 70 % ethanol in order to eliminate any olfactory cues by previously tested mice.

###### 4.3.2 Weighing

**Experiments 1, 2, 3 and 4:** Before each mouse, the weighing vessel (experiment 1, 2, 4: glass jar; experiment 3: cage type 1144B, LWH: 331 x 159 x 132 mm, Tecniplast, Italy) was cleaned with 70 % ethanol in order to eliminate any olfactory cues by previously tested mice.

##### 4.3.3 Millet in Separate Cage

**Experiment 4:** Between mice, the cage was cleaned with 70 % ethanol and bedding material was changed.

**Experiment 8:** Each mouse had its own cage with bedding material for the experimental method, and between experimental days, bedding was removed, cages were cleaned with 70 % ethanol and filled with new bedding material.

**Experiment 9:** Between mice, the floor plate was cleaned with water, when the mice urinated or defecated. Between experimental days, the cage and its floor plate were cleaned with 70 % ethanol.

##### 4.3.4 Restrainer

**Experiment 8 and 9:** Between mice, the restrainer, barriers and seal (Experiment 9: and floor plate) were cleaned with water, when the mice urinated or defecated. Between experimental days, the restrainer was cleaned with special tissues free from alcohol and aldehydes (Microbac Tissues, Hartmann, Germany) and water, while the cage was cleaned with 70 % ethanol.

##### 4.3.5 Fluids

**Experiment 5:** Between mice, the experimental cage was cleaned with 70 % ethanol.

##### 4.3.6 Millet and Bedding

**Experiment 6 and 7:** Between mice, the experimental cage was cleaned with water if mice urinated or defecated.

### 5 Additional Information on CPP experiments

#### 5.1 Experiment 1 - Observations during the Conditioning

Twelve out of thirteen mice went voluntarily into the tunnel through out the whole experiment. There was only one mouse which had to be guided by hand into the tunnel by the 6th session. (It was also the mouse which had to be excluded later on, because it did not change compartments at all during the final preference test.)
However, it was observed (although not precisely recorded) that the duration for leaving the tunnel expanded through out the sessions, when the mice were not confronted with bedding material but the glass jar (weighing) or a grid (fixation) at the end of the tunnel. This hesitant behaviour was more pronounced for the grid than for the glass jar.
During fixation mice urinated and/or defecated in 43 out of 52 times the procedure was performed. During weighing this was never the case.

#### 5.2 Experiment 2 - Observations during the Conditioning

On average, fixation lasted 54.18 $\pm$ 9.50 s, while weighing took 32.15 $\pm$ 5.38 s, both procedures measured from the moment the mouse went from the conditioning compartment into the tunnel

to the moment it went into the tunnel offered directly after the procedure. In addition, mice vocalized in most of the fixation procedures (43 out of 48 times, 89.58%), defecated during half of the fixation procedures (52.08 %), urinated in one third (37.50 %). During weighing procedures, neither vocalization, nor defecation or urination was observed.
It was also observed that some mice seemed to gnaw on the gravel.

#### 5.3 Experiment 3 - Observations during the Conditioning

In general, during baseline it was noted that 6 out of 12 mice defecated (each 3 to 6 faeces) and 4 out of 12 mice urinated while in the conditioning cage. In the final test, 5 out of 12 mice defecated (each 2 to 8 faeces) and 1 urinated while in the conditioning cage.
During the conditioning sessions, mice vocalized in most of the fixation procedures (40 out of 48 times, 83.33%), defecated and urinated during half of the fixation procedures (62.5 % defecation, 56.25 % urination). During weighing procedures, defecation (one faeces) occurred only once and mice never vocalized or urinated. Urination and defecation levels seemed to slightly decrease in the course of the conditioning sessions (session 1: 5/12 mice, session 2: 3/12, session 3: 3/12, session 4: 2/12, session 5: 0/12, session 6: 1/12, session 7: 2/12 and session 8: 1/12 mice).
Fixation on top of the small cage was more complicated (slippery surface) than fixation on a lid and thus, and more fixation attempts were needed before getting the mouse in a good fixation position.
Comparing the latency with which the mice left the tunnel directly before the procedure, mice took 21.18 $\pm$ 23.70 s to move onto the surface for fixation and 6.07 $\pm$ 3.20 s to move into the jar for weighing. This measurement was done on the last day of conditioning (session 7 and 8). However, no baseline latency was measured on day 1, when surface were still unfamiliar.
During the last conditioning session, one mistake occurred: The wrong mouse was placed into the conditioning cage. The error was noted before proceeding to the experimental procedure and mice were switched. Nevertheless, one mouse thus experienced the plate with slits for 3 min at the “wrong” side and without the following experimental procedure.

#### 5.4 Experiment 4 - Observations during the Conditioning

In general, during the baseline it was noted that 8 out of 12 mice defecated (each 4 to 6 faeces) and 10 out of 12 mice urinated while in the conditioning cage. Thus, although mice this time were already in contact before with the conditioning cage, defecation and urination did not decrease compared to the other group before (which did not receive additional habituation sessions in the conditioning cage). During the weighing and reward procedure, mice never vocalized, defecated or urinated. However, defecation and urination inside the conditioning cage occurred in approximately 20 % (urination) or 60 % (defecation) of all sessions. During the final test, 6 out of 12 mice defecated (each 4 to 7 faeces) and 6 urinated while in the conditioning cage.
Comparing the latency with which the mice left the tunnel directly before the procedure, mice took 2.00 $\pm$ 1.40 s to move onto the cage connected with the reward procedure (it might even have been shorter, sometimes they were faster than it was possible to stop the time; for this reason in later experiments we switched to video analysis) and 19.20 $\pm$ 11.73 s to move into the

jar for weighing. In 7 out of 48 times, the mice did not take the millet (only during the first 2 days). In those cases in which the mice ate the millet, it took 25.19 ± 13.17 s for the mice to start feeding and 43.83 ± 20.24 s until they stopped again.
An overview of the progress of latency times (leaving the tunnel as well as start and stop feeding) is depicted in Fig. 8. Here, it is also visible, that the very short latency to leave the tunnel stayed the same for the reward procedure (bedding), while the latency to leave the tunnel for the weighing procedure (glass jar) started high and decreased slowly over the course of days (Fig. 8a). In addition, time to start feeding during the reward procedure decreased over time, although mice did not eat faster, reflected in the consistent stop time (Fig. 8b). Note that the time "stop feeding" does not necessarily correspond to the mice having consumed all of the millet, it could also relate to the mice losing premier interest in the millet consumption.

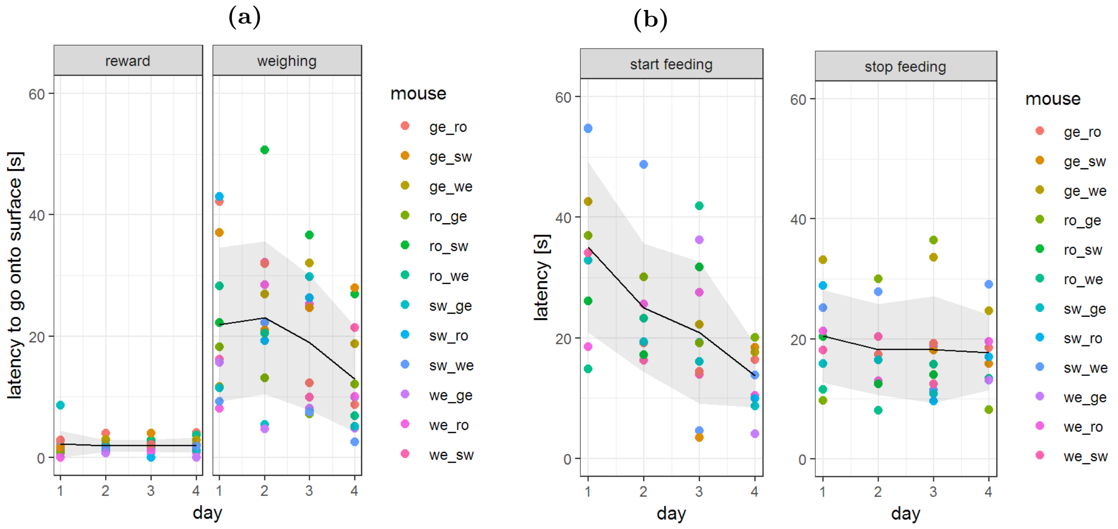

**Figure 8:** Experiment 4: Latency to go from tunnel onto surface (reward and weighing procedure procedures, (a)) and to start and stop feeding (reward procedure (b)). For weighing procedure the mice were placed into a glass jar with a lid, while for the food reward procedure the mice were placed into a type III cage containing bedding material and millet. Time was stopped for holding the tunnel above the surface until leaving the tunnel, from leaving the tunnel until feeding, from the beginning of the feeding until the end. Black line = mean, grey area = standard deviation.

#### 5.5 Experiment 5 - Observations during the Conditioning

Apart from the observations already made about the fluid presentation in section 3.7, which also lead to slight alterations in setup, three things were noted: First, some mice left food prints of almond milk in the handling tunnel and mice were seen licking these droplets from the handling tunnel inside the home cage. Second, nevertheless, not all mice tested the fluids presented and only one mouse was observed drinking it (i.e. testing it for more then just a few seconds). Third, none of the mice licked at the bottle, they all tested the fluid puddle on the floor.

#### 5.6 Experiment 6 - Observations during the Conditioning

All mice fed on the millet, whenever it was offered, and in 85.4 % (41 of 48 cases) not one millet grain was left afterwards. It was also noted that mice in most cases started feeding as soon as they entered the cage without even exploring it first.
During conditioning sessions (3 min in one half of the conditioning cage), defecation and urination levels were low: Only in two cases, one or two faeces were found. However, during baseline and preference test (10 min in complete conditioning cage), levels were much higher: During baseline, nine of twelve mice defecated (one up to six faeces) and two mice urinated. During the preference test (after conditioning sessions), eight of twelve mice defecated (one up to nine faces) and two urinated.

#### 5.7 Experiment 7 - Observations during the Conditioning

All mice fed on the millet, whenever it was offered, and in two of three cases (32 of 48 sessions) not one millet grain was left afterwards. Only in the first session one mouse did not feed at the millet at all.
During conditioning sessions (3 min in half of the conditioning setup), defecation and urination levels were low: Only in three cases, one faeces was found. However, during baseline and preference test (10 min in complete conditioning apparatus), levels were higher: During baseline, five of twelve mice defecated (two up to five faeces) and one mouse urinated. During the preference test (after conditioning sessions), four of twelve mice defecated (one up to two faeces) and none urinated.

#### 5.8 Experiment 8 - Observations during the Conditioning

All mice fed on the millet, whenever it was offered.
During conditioning sessions (1 min procedure + 3 min in one half of the conditioning apparatus), in about 23 % of cases mice defecated (22 of 96 sessions), and they never urinated. During baseline and preference test (10 min in complete conditioning apparatus) defecation and urination levels were similar: During baseline, four of twelve mice defecated and two mice urinated. During the preference test (after conditioning sessions), three of twelve mice defecated and none urinated.
Although some mice refused to go immediately into the restrainer when guided to it, mice always went into the handling tunnel by themselves and also left them voluntarily. Thus, tunnel handling was not impaired by the resemblance between handling and restrainer tunnel.

#### 5.9 Experiment 9 - Observations during the Conditioning

During millet procedure, only one mouse defecated (in the last conditioning session), while during restraint procedure, four mice defecated / urinated. However, in comparison to the other experiments, defecation / urination levels were low, only one of the three mice kept defecating through all four conditioning sessions. Notably, this was also the mouse which consumed the most millet grains during the millet procedure (see Table 1). During baseline measurement, two mice defecated (one to three faeces) and one mouse urinated. During the final test, no defecation or urination was observed.

Table 1: Experiment 9: Eaten millet grains and defecation / urination levels during the procedures. Defecation occurred only during the restrainer procedure, millet could only be eaten during the millet procedure. Mice had access to 16 grains of millet (approximately 0.1 g). Trial 1 corresponds to session 1 or 2, depending on the procedure which was experienced first by this mouse (see first column). Below the individual columns, the average is per trial for this procedure is calculated. Defecation also includes urination. One mouse also defecated during millet procedure (sw_ro, trial 4, 3 faeces). In marked in red are noticeable observations (consumed no grains / all grains or defecated).

| | | session 1 | session 2 | session 3 | session 4 | session 1 | session 2 | session 3 | session 4 |
|---|---|---|---|---|---|---|---|---|---|
| mouse | first procedure | millet (grains eaten) | | | | restrainer (feces) | | | |
| ro_ge | millet | 13 | 14 | 14.5 | 13 | 0 | 0 | 0 | 0 |
| ro_si | restrainer | 14 | 14 | 9 | 10.5 | 0 | 0 | 0 | 0 |
| ro_sw | restrainer | 15 | 14 | 13 | 11.5 | 0 | 0 | 0 | 0 |
| ro_we | millet | 12 | 12 | 12 | 12 | 0 | 0 | 0 | 0 |
| sw_ge | restrainer | 10 | 0 | 12 | 15 | 0 | 0 | 0 | 0 |
| sw_ro | restrainer | 16 | 15 | 15 | 14 | 3 | 2 | 4 | 2 |
| sw_si | millet | 10 | 12 | 11 | 10 | 0 | 0 | 0 | 0 |
| sw_we | restrainer | 2 | 15 | 9.5 | 15 | 1 | 0 | 0 | 0 |
| we_ge | millet | 4 | 12 | 13 | 11 | 0 | 0 | 0 | 0 |
| we_ro | millet | 15 | 14 | 12 | 15 | 0 | 0 | 0 | 0 |
| we_si | restrainer | 0 | 11 | 11.5 | 10 | 0 | 0 | 0 | 0 |
| we_sw | millet | 12 | 14 | 13 | 14 | 2 | 0 | 0 | 0 |
| average | millet | 10,33 | 9,17 | 13,00 | 14,00 | 0,67 | 0,33 | 0,67 | 0,33 |
| average | restrainer | 10,17 | 12,83 | 11,25 | 11,80 | 0,33 | 0,00 | 0,00 | 0,00 |

##### 5.9.1 Additional Video Analysis in Experiment 9

To record the latency to leave the handling tunnel, habituation and conditioning sessions were filmed from the side (procedure environment) and above (setup). Videos were analysed using the open source program BORIS (Behavioral Observation Research Interactive Software, Version 7.9.8, Friard and Gamba 2016). The latency was measured as the time point when the tunnel was placed above the floor, to the time point when the mouse left the tunnel with all four paws. Data was analysed with regard to floor pattern, procedure and session. We compared the latencies of trial 1 of the procedure/pattern (session 1 and 5) with the last trial of the procedure/pattern (session 4 and 8), as well as the last trials of each procedure/pattern against each other. To test for normal distribution, the Shapiro–Wilk test was performed in R. If the data was normal distributed ($p > 0.05$), we performed a Welch Two Sample t-test to compare the latencies. If it was not (in the case of trial 1 of each procedure, consisting of session 1 and 5), a Dependent-samples Sign-Test was conducted. In all statistical tests, significance level was set to 0.05.

##### 5.9.2 Results Latency

Habituation: We had four days in which we "practised" the procedure with the mice for habituation. As can be seen in Fig. 9a, mice decreased their latency to leave the handling tunnel during these four habituation trials. In addition, latency to enter the conditioning setup was shorter compared to the latency to enter the type II cage which was later on used as procedure environment. This was probably caused by the transparent walls of the type II cage and the opaque walls of the conditioning setup.

Conditioning sessions: Latency to leave the tunnel into the procedure environment decreased during conditioning sessions for both procedures (millet environment: $S = 11$, p-value $= 0.006348$; restrainer environment: $S = 10$, p-value $= 0.01172$, see Fig. 9b). Comparing the fourth conditioning session of both procedures, however, there was no difference between procedures (S = 5, p-value = 0.7744).

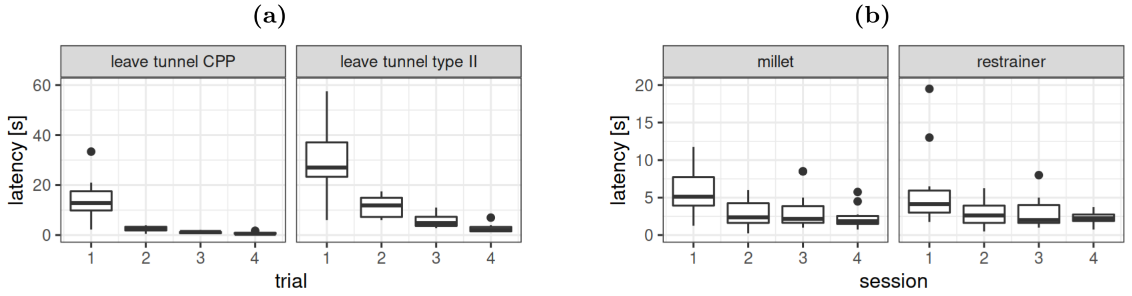

**Figure 9:** Experiment 9: Latency to leave the handling tunnel into the conditioning setup ("CPP") and the into type II cage (later procedure environment) during the four habituation sessions (a). (b) Latency to leave the handling tunnel into procedure environment during the conditioning sessions, when it was either the millet environment or the restrainer environment (differing by floor plates with other tactile structures).

### 6 Discussion: Latency to Leave Tunnel

In experiment 4 and 9 (and 8.1, no CPP experiment), in addition to the CPP, we measured the latency of the mice to leave the handling tunnel. Already in experiments 1 – 3, we had observed that with progressing conditioning sessions mice took more time to leave the tunnel when confronted with the environment in which they were fixated (experiment 1 and 2: grid, experiment 3: an upside-down cage) than when confronted with the environment in which they were weighed (experiment 1 and 2: glass jar, experiment 3: cage). Here, the procedure environment seemed to work as a CS on its own. However, in experiments 1 and 2, for the latency to leave the handling tunnel only descriptive notes were taken. In experiment 3, the latency was only measured during the last conditioning session. Thus, no comparison to a start latency and a potential bias between the different surfaces is possible.

In experiment 4, time was measured with a timer. However, there was a great difference between the used environments right from the start (cage with bedding vs. glass jar), so the data was not comparable. In experiment 8.1 (o CPP experiment), we performed an experiment only focussing on this latency to leave the tunnel. Here, we used visual in addition to tactile stimuli (because this group of mice had no whiskers) and compared access to millet with restraint in a self-made restrainer. No difference in the latency to leave the tunnel was found.

Arguing that this could be due to the whisker-loss, we included this measurement again in experiment 9 via analysis of the video recordings. Here, to distinguish the experimental environment only tactile stimuli were applied. Again, no difference was found. This could be due to the chosen procedure (restraint), in which the mouse was not in direct contact with the tactile stimulus but with the restrainer tube. Nevertheless, to get the mice into the restrainer, they had to be guided or sometimes chased into the tube, which itself might have worked as an aversive US, and this happened in direct contact with the tactile stimulus.

This missing association between procedure environment and procedure as seen in experiment 8.1 and 9 is very interesting. It shows that although sometimes (experiment 1 – 3) an association between procedure environment and procedure is formed, and causes mice to leave the tunnel more hesitantly, sometimes it is not. What causes the difference? In experiment 1 – 3 we used fixation by hand, in 8.1 and 9 restraint in a restrainer. Is it necessary to have an US with a more negative emotional effect to induce this form of conditioning? If so, could we now say that a short fixation for 20 s was therefore more severe than staying in a restrainer for 1 min? Or is it the procedure itself which facilitates the association because during fixation the mice have to be pressed shortly against the ground, and therefore are forced to have a more direct contact with the environmental stimuli? Unfortunately, these are questions which we are not able to answer with our experiments but will have to pass them on for future studies.

### 7 Experiment 4.1 (Choice Test)

#### 7.1 Motivation

This experiment was more similar to operant (instrumental) conditioning or a choice test than to CPP, which is why it is not included in the main paper. Here, the decision of the mice to stay in the left or right compartment (on which flooring material) determined the experimental procedure which was then performed with the mouse. Thus, instead of assessing what causes a more positive or negative association with a cue (and therefore leads to preference or avoidance), this test assesses how the mice behave when confronted with the stimulus: stay to perform the procedure or leave to get to perform the other procedure, similar to a Go/No-Go task (Frederick et al. 2011, Jones et al. 2017).

#### 7.2 Procedure

In ten trials, each mouse chose between the two compartments of setup 2 (and their flooring materials) and, by extension, between two experimental procedures (weighing and restraint in a restrainer). We expected the choices for the side corresponding to the preferred procedure to accumulate over the course of trials.
Experiments were conducted with group 2. Mice were tested individually and all ten trials consecutively on one day. To ensure that mice were only tested in the morning (close to their active phase), which meant three mice per day maximum. On experimental days, mice were taken to the experimental room in one of their home cages. (The connection between the two cages was removed and the hole in the cage wall sealed. This was done to reduce the possible additional disturbance for the mice not tested on this day which would have been caused by using a transport cage.) Before the start of each session, mice had 30 min of habituation to the experimental room. After the sessions, the mice were transported back to the husbandry room and both cages were reconnected. Between mice, the cage and experimental equipment were cleaned with 70 % ethanol.
Although planned differently, the experimental design was changed twice due to observations during the conduction, leading to three different experimental phases:

##### 7.2.1 Phase 1

The first three mice started with the initially planned design: Each mouse performed ten trials in close succession, with each trial starting immediately after the last. Each mouse performed 10 trials. The number of mice used for the experiment per day depended on the time required for each mouse. This was the reason why the whole group was transported into the conditioning

room each day.
For the test, each side of the test cage contained a different kind of metal plate with different surface structure as CS (metal grid or metal plate with holes), separated by a small barrier (setup 2). Mice were taken individually out of the cage by tunnel handling according to a predefined randomized order. At the start of each trial, the handling tunnel was held directly above the small middle barrier. After the mouse had left the tunnel (the other tunnel opening was closed by a hand), it had 15 s to choose a side. Then, the second half of the cage was made inaccessible by an opaque wall. In addition, a transparent plate was placed on top of the conditioning cage. The mouse was given 30 s inside the chosen half in contact with the CS (the flooring metal plates). Then, the mouse was taken out of the cage by tunnel handling and the respective procedure (US, weighing or 1 min in a restrainer) was performed. Afterwards, the mouse was returned by tunnel handling to the conditioning cage to start a new trial (15 s to choose, followed by 30 s with the CS, followed by procedure).
We decided against forced trials for both sides at the beginning because we expected the mice to alternate choices before the correlation was learned (similar to observations made in a T-Maze choice experiment, Habedank et al. 2021). However, it was observed during the first experiments that some mice did not show this expected behaviour. Instead, they often repeated their choices. Therefore, after the sixth repeated choice without any experience of the other procedure, the mouse was forced to go onto the other side of the cage. In addition, if a mouse chose again the restraint side after seven experienced restraint procedures (in total), it was directly returned to the home cage to reduce further stress instead of conducting the procedure. (This happened only in the last of the ten trials, so all ten choices were made but for the last choice, no procedure was conducted.)
If a mouse had not left the tunnel for 2 min when being held above the middle barrier to choose a side, the tunnel would have been tilted until the mice would have slipped softly out of the tunnel. The mouse would have been then given the chance to change the side for 15 s. However, this case never occurred.

##### 7.2.2 Phase 2

Already with the first three mice, the expected learning was not observable, as stated above. It is possible that the 15 s are to short a time for the mice to recover from the last procedure and focus on the task (choice). In a next step, we therefore decided to offer a break between trials without changing the procedure all together. Therefore, we changed the procedure: The next two mice performed the trials alternating, meaning after one trial with mouse A, the next trial was conducted with mouse B, and after that again with mouse A and so on. Thus, mice had a break of about 7 min before the next trial. Here, the setup and the used procedure equipment were cleaned with 70 % ethanol after each trial, because after each trial the mouse was exchanged. The rest of the procedure remained the same.

##### 7.2.3 Phase 3

The alterations also seemed not to enable the mice to understand the offered choice, i.e., no cumulation of a preferred side was observed. To prevent the last seven mice from performing the restraint in vain (without any useful results), we decided to alter the focus: Mice were now video recorded in the conditioning cage, extending the time frame of the choice to make an analysis of behaviour possible: In comparison to phase 1, the mice had now 2 min instead of 15 s to switch sides. During this time, mice were video recorded from above, using a webcam. After the 10th trial, mice were once again returned to the conditioning cage for an additional recording of 2 min before their return to the home cage. In this manner, it would have been possible for the mice to learn the correlation between stay on one side and the procedure and we would also get the observations needed.

#### 7.3 Additional Video Analysis in Experiment 4.1

For seven out of twelve mice, video recordings were made during the trials. We decided to analyse several behavioural parameters to get a general impression.
To compare the activity over the course of the trials, a virtual grid of 4 * 4 squares was placed above each half of the cage and it was counted how often mice stayed in each square during the 2 min at the start of each trial (before the separation of the sided). A square was defined as occupied whenever at least half of the mouse was on it. For quantification of activity, we analysed the area covered (how many squares were occupied at least once) and how many squares were passed in total (taken into account whether a squares was occupied multiple times).
Next, the video recordings were analysed for specific behaviours with the help of the open source program BORIS. We counted events of rearing (raise on the hind legs with stretched back, head oriented upwards) and stopped the time of grooming (based on Kalueff et al. 2007). We also counted whether a mouse started grooming within the first 10 s of a trial and how long the mouse did not move at all (without even head or nose moving). To keep it short, we will only report the major findings in the results.

#### 7.4 Results and Discussion

##### 7.4.1 Observations during the Experiment

After the first three mice (phase 1) it was already clear that these mice did not learn the correlation between side (or floor plates, respectively) and procedure. The first two mice just randomized their choices, leading to 50 % restraint procedures. The third mouse, however, stuck to the restraint side without trying the other side. Even after the forced choice (i.e.:after the sixth choice for one side without experiencing the other, the mouse is forced onto the other side) the mouse returned to the restraint side, although the stress already seemed to have a visible effect on the behaviour of the mouse (less movement, changed breathing).
For this reason, on the next experimental day mice performed trials alternately (see section 7.2.2). However, there was no effect of this change observable, here again one mouse chose the restrainer in 70 % of the choices.
With the argumentation that the remaining seven mice should not be tested in vain, the conduction was again slightly altered (see section 7.2.3) and the behaviour of the mice was video recorded.

##### 7.4.2 Choices

As mentioned above, during the conduction of the experiment it became clear that mice showed no preference for side with the supposedly less stressful procedure (weighing). In addition, some mice kept returning to the same side trial after trial independent from the associated procedure (e.g., two mice choice one side 9 out of 10 times, three mice chose the same side 7 or 8 times). This might be influenced by a material preference: On average, in 6.5 ± of the 10 trials the grid was chosen ($p < 0.01$, $t = 3.3166$) or in other words: 9 of 12 mice chose the grid at least six times. There was no general preference for side (on average in 5.16 ± of 10 trials left was chosen, $p = 0.7986$, $t = 0.26139$), procedure (5.16 ± of 10 trials restraint was chosen, $p = 0.7986$, $t = 0.26139$) or the compartment not chosen last (p-value = 0.7439, $t = -0.33503$). A summary is depicted in Fig. 10. Note that the results are merely for a general impression and have to be interpreted with caution as the experimental procedure was changed multiple times.

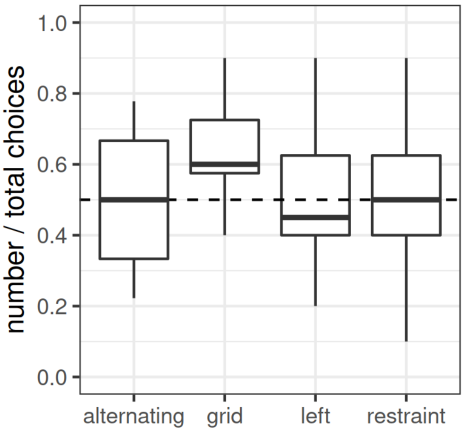

**Figure 10:** Experiment 4.1: Choices made by the twelve mice in the ten trials for the compartment which they did not choose before (alternating), the grid compartment (grid), the left compartment (left) or the compartment which was followed by the restrainer procedure (restraint). Note that grid, left and restraint contain ten choices, while alternating only contains nine (the first choice is independent). The dashed line represents chance level. Grid was preferred significantly over the metal plate with holes ($p < 0.01$).

##### 7.4.3 General Note on the Following Behaviour Results

The results now following are only of phase 3, using 7 mice. In addition, although we analysed the results as being mainly influenced by the last procedure, it is possible that the behaviour reflects a cumulative effect of all the procedures experienced so far. For example, a mouse which experienced restraint for the first time on this day might behave differently than a mouse which experienced it for the fifth time. However, we nevertheless summed up the behaviour of all trials, regardless of the trial number it was and what the choices were made before, to get a general impression. As some mice experienced one of the procedure more than others, this might have led to a strong influence of individuals. To prevent this, all results were first calculated as the mean per mouse and then for the whole group.

In addition, please note again that all of the following results are merely meant for a general impression to generate hypotheses, as they are received from an experiment with a changing procedure protocol.

##### 7.4.4 Activity

In the following, "squares" refers to the distance the mice moved, while "area" represents which squares were visited at all. A summary of the results is depicted in Fig. 11.
During the baseline, mice nearly used all squares. When looking at the activity after the procedures, the number of squares which were crossed was largely reduced. However, the number of squares crossed had a tendency to be significantly higher after the weighing procedure compared to restraint procedure ($p = 0.06387$, $t = -2.2677$).
Analysing the area that was covered during the 2 min in the experimental cage, after restraining the area used was in tendency smaller than after weighing ($p = 0.05874$, $t = -2.3287$).
Comparing changes in squares crossed and area covered with the baseline, mice seemed to change their activity stronger with regard to the distance they travelled (between 50 and 75 %) than with regard to the area in which they stayed (between 0 and 50 %).
In addition, we took a look at border areas. As border area or periphery we defined the 20 squares close to the wall, thus, including about 2/3 of the whole area. In theory, if the mice distributed their stay equally across all squares, they should have stayed to 62.5 % in border squares. In practice, the percentage was a bit higher but nearly the same for baseline and after the weighing procedure. After restraining, the squares the mice crossed were to a higher amount border squares compared to weighing (79.16 $\pm$ 8.06 % of squares crossed were border squares after restraining, 68.24 $\pm$ 7.12 after weighing, $p < 0.02$, $t = 3.1666$). For areas this difference was not significant.

##### 7.4.5 No Movement

During the conduction of the experiments as well as the analysis of the videos with regard to activity, we noted that the mice sometimes seemed to stop moving at all or "freeze". In such moments, not even the nose or ears moved but the mouse stayed motionless without any obvious focus. We noted the start and ending of these "no movement" behaviours during video analysis and calculated the total amount of time, the mice spent this way during each 2 min observation in the experimental cage. However, it has to be noted, that behaviours were only registered, if they were longer than 1 s.
During the baseline observation, mice spent 0.46 $\pm$ 1.23 s without movement, after restraint procedure 3.34 $\pm$ 2.41 s and after weighing 4.88 $\pm$ 3.43 s. Thus, the behaviour appeared equally after both procedures ($p = 0.3098$, $t = -1.1093$).

##### 7.4.6 Grooming

Using the recorded videos, the time mice spent grooming (face, body, tail or legs) was measured. Grooming increased after weighing and even more after restraint in comparison to the baseline measurement (see Fig. 12a). The difference between weighing and restraint procedure was significant ($p < 0.04$, $V = 27$). Duration of grooming bouts was similar after weighing and restraining (9.83 $\pm$ 5.23 s after restraint, 8.06 $\pm$ 5.73 s after weighing, $p = 0.2969$, $V = 21$).
Interestingly, mice started grooming within the first 10 s after entering the cage only after restraining but not after weighing (or during baseline). Mice exhibited this behaviour in 77.55 $\pm$ 19.90 % of measurements after restraint procedure. Reducing the time frame to only 5 s after entering the experimental cage still left 61.94 $\pm$ 25.96 % of trial after restraint, in which this

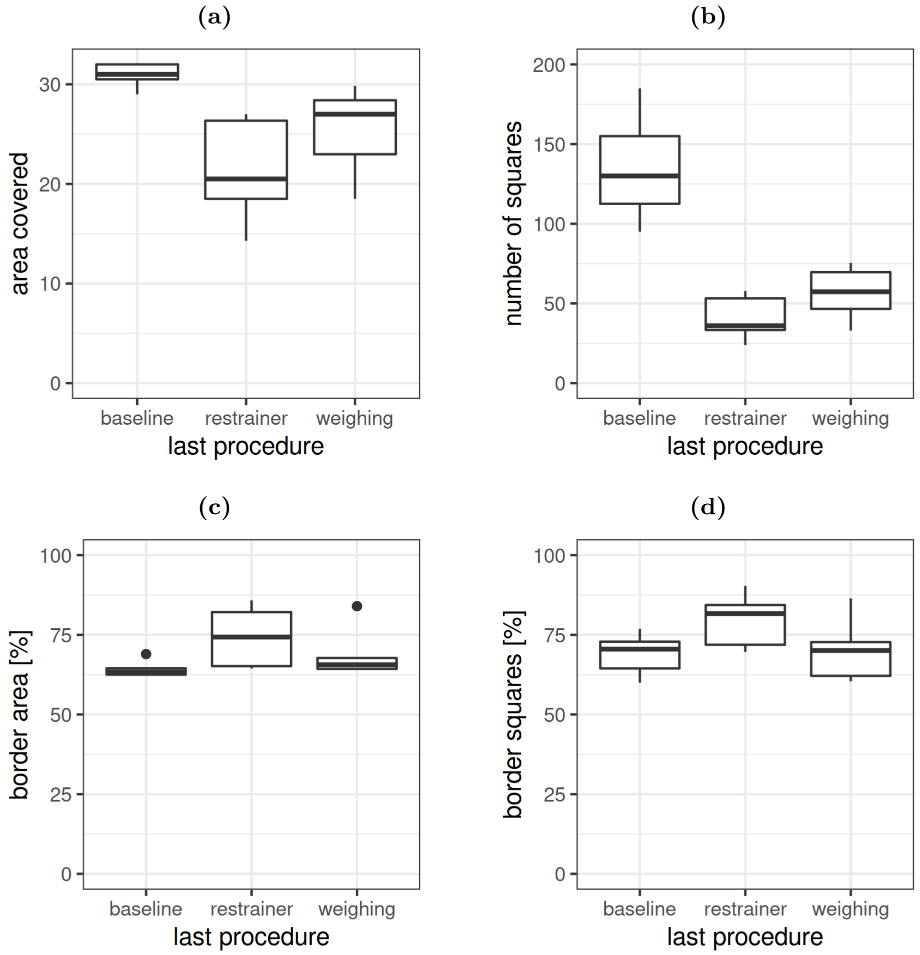

**Figure 11:** Experiment 4.1: Activity during baseline and after the procedures. (a) Area covered (squares visited at least one time). (b) Total number of squares visited (distance). (c) Percentage of border area visited in comparison to total area. (d) Percentage of border squares covered in comparison to total number of squares covered (distance). Note that the data is only of phase 3, using 7 mice, and that the results are summed up for all mice and all trials, although the mice experienced differing repetition numbers.

behaviour was shown. This is probably related to the restraint procedure itself: When mice defecated or urinated, they automatically wet their fur due to the reduced space.

##### 7.4.7 Rearing

After weighing, rearing behaviour was reduced compared to baseline, however, after restraining, it even dropped even further within the two recorded minutes after the procedure (see Fig. 12b). This difference between weighing and restraint was significant ($p < 0.005$, $t = -4.3232$).

### 8 Experiment 4.2 (Behavioural Comparison)

#### 8.1 Motivation

Based on the results from experiment 4.1, phase 3, we now conducted one experiment which did *not* focus on conditioning the mice to associate a specific stimulus with a specific procedure, but on the behaviour after the experimental procedure (formerly the US).

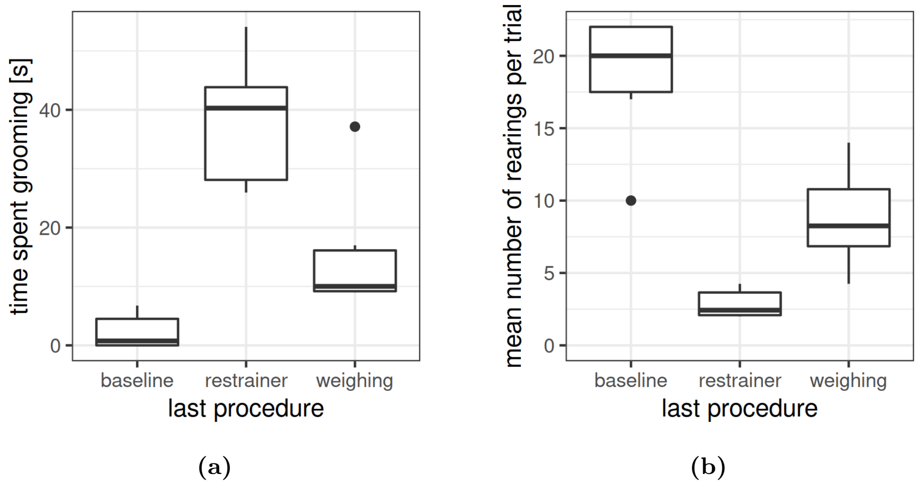

**Figure 12:** Experiment 4.1: Grooming and rearing behaviour during baseline and after the procedures. (a) Grooming duration. (b) Number of rearings per trial. Note that the data is only of phase 3, using 7 mice, and that the results are summed up for all mice and all trials, although the mice experienced differing repetition numbers.

#### 8.2 Procedure

Compared were a supposedly neutral procedure (home cage) with a potentially aversive procedure (open field) (round 1). As the open field might be a test which is very mild and might not be sufficient to cause behavioural changes compared to time spent in the home cage, we then repeated the test, using (instead of the open field test) a procedure which will be called hereafter "separation" (round 2). In previous experiments in our laboratory, this procedure had an effect at least on the home cage activity of the mice (own observations), so we assumed it should have a directly measurable effect on behaviour.

In each round, the test was conducted on four consecutive days, with each procedure tested two times (one trial on two days). The experiment was conducted with group 2 (n = 12). The order of mice taken for the test was randomized beforehand for each of the four experimental days. In addition, pairing of day and performed procedure was randomized, so that half of the mice performed one procedure and half of the mice the other procedure, before procedures were switched on the next day.

The tests took place in the husbandry room, so no transportation and habituation time to the room was necessary. However, during the conduction of the experiment, the filter tops had to be removed from both cages to facilitate handling of the mice. Thus, even the mice which were currently not outside the cage experienced a change in the home cage environment during the tests.

For each trial, the tunnel with the mouse was held directly above the small middle barrier separating the two halves of the experimental cage. After the mouse left the tunnel (the other opening was closed by a hand), it stayed for 2 min inside the experimental cage (setup 2). This time was used as a baseline measurement. During the stay in the experimental cage, a transparent plate was placed on top of the experimental cage to prevent the mouse from leaving the cage. Afterwards, the mouse was taken out of the cage to perform the perspective procedure (round 1: return to home cage vs. open field test, round 2: return to home cage vs. separation). The mouse was then placed back into the experimental cage for another 2 min (also with the transparent plate on top), before returning to the home cage.

Between mice, the cage and experimental equipment were cleaned with 70 % ethanol. During the first 2 min as well as during the second 2 min in the experimental cage, the mice were video recorded from the side and above for later analysis of their behaviour (see following section).

##### 8.2.1 Additional Video Analysis in Experiment 4.2

Video recordings of the time the animals spent in the experimental cage (2 min) were analysed, starting at the moment the mice left the handling tunnel and had all four paws on the floor of the experimental cage. The recordings were analysed for specific behaviours with the help of the open source program BORIS (Behavioral Observation Research Interactive Software, Version 7.7.5, Friard and Gamba 2016): number of supported rearings (raise on the hind legs with stretched back, head oriented upwards, forelegs supported on the cage wall or the barrier), number of unsupported rearings (raise on the hind legs with stretched back, head oriented upwards, forelegs in plain air), time spent grooming the face (paws stroke face and ears), time spent grooming the body (licking the fur).
To compare the activity over the course of the trials, we used an open source software package for animal pose estimation called "Deep Lab Cut" (`http://www.mousemotorlab.org/deeplabcut`, Nath et al. 2019) which is based on transfer learning with deep neural networks. Video recordings were cut beforehand using ShotCut (version 18.01.02, Meltytech, LLC) to contain the exact 2 min for analysis. Four videos out of the in total 96 videos were chosen for training, using the first videos of four mice which had slightly different colouring of their tail, to train the network with distinct appearances. Additionally, frames were excluded for which position change between the frames (velocity) exceeded the reasonable level and / or position of the tail root was detected outside the borders of the experimental cage floor.
During round 2, we used a different camera: a Go Pro camera (GoPro Hero7 Black, Inc., USA) camera instead of a WebCam. This camera was shifted multiple times for technical reasons (battery change). Thus, the coordinates of the experimental cage floor (used for exclusion of "wrong" tail root positions) had to be adjusted for each video. To do so, we sorted the videos into groups, for which the border position was the same, before running the analysis.

#### 8.3 Results and Discussion

##### 8.3.1 Observations during the Home Cage Procedure

As a control, we chose to return the mice to the home cage, expecting this to be a neutral or even some kind of "resetting" procedure, and thus, basically "no procedure". However, during the conduction of both experiments, it was observed that sometimes mice took longer (> 20 s) to enter the handling tunnel and even had to be guided by hand into the tunnel. In some cases this included slight chasing and consequently agitated the mice. As a comparison: During the separation procedure, mice had to be guided only one time by hand back into the handling tunnel, and it was never the case after the open field test.
In addition, it was observed that while some mice spent their time in the home cage feeding, other mice ran frequently from one cage to the other, thus also causing a different state of agitation for the mice. Thus, the home cage time might not have been working as a comparative, neutral procedure as expected.

##### 8.3.2 Activity

To get information on general activity, we tracked the position of the tail root during the 2 min in the experimental cage with the help of DeepLabCut.
Round 1: Comparing the distance, the tail root moved during this period, we found no difference comparing open field and home cage (p = 0.2605, t = 1.1862, see Fig. 13a). Comparing the difference between the distance moved in the baseline and after the procedure, there was a tendency towards a larger difference after open field compared to home cage (p = 0.07093,t = 1.9989).
Round 2: The distance the tail root moved did not differ significantly between the separation procedure and home cage (p = 0,146, S = 9, see Fig. 13b). Also, comparing the values to the baseline, there could no difference be found (separation: 34.00 ± 67.88 % difference to baseline, home cage: 17.82 ± 27,.90 % difference to baseline, p = 0.146, S = 9).

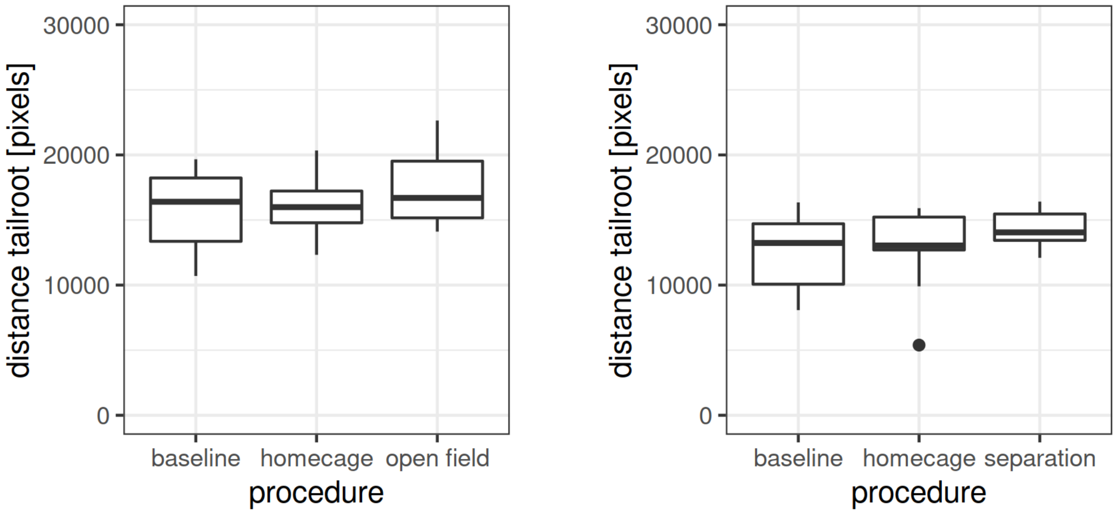

(a) (b)

**Figure 13:** Experiment 4.2: Activity (distance travelled) during baseline and after procedure. (a) Round 1, comparing home cage and open field. (b) Round 2, comparing home cage and separation. Results the average of two trials (one per day).

##### 8.3.3 Grooming

During round 1, open field vs. home cage, mice never groomed longer than 15 s during the 2 min. During round 2, separation vs. home cage, in seven measurements grooming lasted longer than 15 s (performed by five mice). For both experiments, time spent grooming grooming was not normal distributed and did not differ significantly between procedures (round 1: p = 0.1041, t = 1.7715; round 2: p = 0.7744, S = 5).

##### 8.3.4 Rearing

In both rounds, the number of rearings (supported and unsupported) did not differ significantly during procedures, neither for open field vs. home cage (round 1: 18.13 ± 4.,31 rearings after open field vs. 17.5 ± 4.22 rearings after home cage, p = 0.7324, 0.35077), nor for separation vs. home cage (round 2: 15.58 ± 4.48 rearings after separation vs. 14.96 ± 6.32 rearings after

home cage, p = 0.7844, t = 0.28035). However, in round 2, rearing numbers seemed to show a decreasing development across trials (data not shown).
In an additional analysis, rearings were divided into supported and unsupported rearings. Although there was no difference between procedures for unsupported rearings in both rounds (Wilcoxon signed rank test, round 1: p = 0.2078, V = 55.5; round 2: p = 0.7598, V = 31), number of supported rearings differed significantly between open field and home cage (round 1: $p < 0.03$, t = 2.5404, see Fig. 14a). This was not the case in round 2 (p = 0.1443, t = 1.5719, see Fig. 14b).

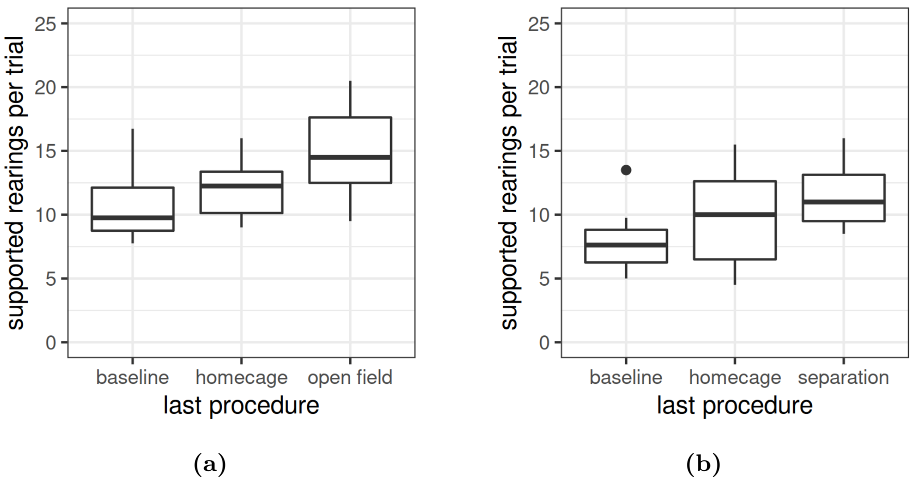

**Figure 14:** Experiment 4.2: Number of supported rearings during baseline and after procedure. (a) Round 1, comparing home cage and open field. (b) Round 2, comparing home cage and separation. Results the average of two trials (one per day).

### 9 Experiment 8.1 (Latency to Leave Tunnel)

#### 9.1 Motivation

Analysing all the CPP experiments we conducted so far, we found that although the place conditioning procedure did not work, in experiments 1 to 3 we observed that mice seemed to be more hesitant to leave the handling tunnel during conditioning sessions, if an unpleasant procedure such as fixation followed. However, this was merely a side observation (without accurate time measurement) and we now wanted to investigate it in an experiment.

#### 9.2 Procedure

In this experiment, we did not perform a conditioned place preference test but a more basic conditioning procedure: By tunnel handling mice were placed in an experimental cage with a specific CS and latency to leave the tunnel was recorded by a video camera. Mice had 15 s to explore the cage (and the CS) before the procedure started (US). Afterwards, mice were transported back to there home cage by tunnel handling.
One habituation session was conducted, during which mice were placed for 15 s inside the experimental cage without any following procedure. Then, mice experienced four sessions of each

experimental procedure, a two-day-break (weekend) and then four sessions with the other procedure. Which procedure and which pattern was experienced first was different for half of the mice, also pairing of CS and US differed for half of the mice.
As CS we used dots and stripes as visual wall patterns and plastic plates with slits or holes as visual and tactile floor stimuli. As US we either added 0.01 g millet to the experimental cage and mice had 1 min access to it, or the mice was transferred into a restrainer (placed in the experimental cage), in which the mice had to stay for 1 min.
The experiment was conducted in the husbandry room. Before the start of any experimental procedures, the filter top of the home cage was removed and mice were given 10 min for habituation to the altered illumination. The experiment was conducted in the morning.
The experimental cage (as depicted in Fig. 4e and f) was fit into a metal construction with a camera (Logitech C390e WebCam, Switzerland) which filmed the procedure from above using the open source recording program iSpy 64 (version 7.0.3.0). This metal construction was built out of MakerBeams (MakerBeam B.V., The Netherlands).
We used mouse group 3, and at the beginning of the experiment, the mice were about 12 months old. It was known before the experiment that eleven of the twelve mice lacked their whiskers (partly or completely), probably due to plucking / barbering. Although it is reported, that missing whiskers may lead to an alteration of behaviour due to the missing tactile information, we decided against ordering a new group of mice for this experiment. Instead we adjusted the experimental design (e.g., used visual in addition to tactile cues as conditioned stimuli). Mice experienced millet in previous experiments.

###### 9.2.1 Additional Video Analysis in Experiment 8.1

Videos were analysed using the open source program BORIS (Behavioral Observation Research Interactive Software, Version 7.9.8, Friard and Gamba 2016). The latency to leave the tunnel was measured as the time point when the tunnel was placed above the experimental cage floor to the time point when the mouse left the tunnel with all four paws.
Data was analysed with regard to floor pattern, procedure and session. We compared the latencies of trial 1 of the procedure/pattern (session 1 and 5) with the last trial of the procedure/pattern (session 4 and 8), as well as the last trials of each procedure/pattern against each other. To test for normal distribution, the Shapiro–Wilk test was performed in R. If the data was normal distributed ($p > 0.05$), we performed a Welch Two Sample t-test to compare the latencies. If it was not (in the case of trial 1 of each procedure, i.e., sessions 1 and 5), a Dependent-samples Sign-Test was conducted. In all statistical tests, significance level was set to 0.05.

##### 9.3 Results and Discussion

###### 9.3.1 Observations during the Conduction

Disclaimer: In the following, only descriptive statistics are given, no statistical testing was done. Mice only defecated during the restrainer procedure, never during the millet procedure (see Table 2). Defecation levels seemed to decrease during the trials, and seemed to be higher for the group that started with the restrainer procedure (and switched to millet in the second week).

Only during one millet trial one mouse did not consume a single grain, during all other trials mice consumed at least one grain. Mice seemed to consume more grains with each trial. In addition, mice which experienced millet in the second week (day 6 to 9) consumed more grains from the start than the other group (which experienced millet before the restrainer procedure). Moreover, at trial 4 of the first week, both groups had similar performance. This could indicate that there is a habituation effect to the setup itself, and more days of habituation are needed to lead to the maximum consumption / best performance.
It was also noted that some mice took their time to enter the handling tunnel after the procedure and sometimes even re-entered the restrainer tunnel before returning to the handling tunnel.

Table 2: Experiment 8.1: Eaten millet grains and defecation levels during the procedures. Defecation occurred only during the restrainer procedure, millet could only be eaten during the millet procedure. Mice had access to 0.1 g millet, which were between 13 and 17 grains. Only one mouse consumed all grains (represented by (all)). Trial 1 corresponds to session 1 or 5, depending on the procedure which was experienced first by this mouse (see first column). Below the individual columns, the average per trial for this procedure is given.

| | | trial 1 | trial 2 | trial 3 | trial 4 | trial 1 | trial 2 | trial 3 | trial 4 |
|---|---|---|---|---|---|---|---|---|---|
| mouse | first procedure | millet (grains eaten) | | | | restrainer (feces) | | | |
| ro_ge | restrainer | 0 | 13 | 15 | 11 | 0 | 0 | 0 | 0 |
| ro_si | restrainer | 13 | 14 | 14 | 13 | 0 | 1 | 1 | 0 |
| ro_sw | millet | 5 | 4 | 10 | 11 | 0 | 1 | 0 | 0 |
| ro_we | restrainer | 12 | 12 | 15 | 10 | 0 | 1 | 0 | 0 |
| sw_ge | millet | 1 | 2 | 13 | 15 | 0 | 0 | 0 | 0 |
| sw_ro | millet | 3 | 1 | 11 | 13 | 1 | 0 | 0 | 0 |
| sw_si | restrainer | 11 | 10 | 13 | 11 | 2 | 2 | 0 | 0 |
| sw_we | millet | 7 | 6 | 13 | 13 | 2 | 0 | 1 | 1 |
| we_ge | millet | 5 | 6 | 13 | 15 (all) | 0 | 0 | 0 | 0 |
| we_ro | millet | 1 | 11 | 10 | 12 | 2 | 2 | 0 | 0 |
| we_si | restrainer | 4 | 13 | 13 | 14 | 1 | 0 | 0 | 0 |
| we_sw | restrainer | 13 | 14 | 13 | 12 | 3 | 4 | 2 | 1 |
| average | millet | 3,67 | 5,00 | 11,67 | 12,80 | 0,83 | 0,50 | 0,17 | 0,17 |
| average | restrainer | 8,83 | 12,67 | 13,83 | 11,83 | 1,00 | 1,33 | 0,50 | 0,17 |

##### 9.3.2 Latency to leave the tunnel

Although there was a habituation session beforehand, during the habituation day and first contact with the setup mice took longer to leave the tunnel (see Fig. 15, experimental day 1 / trial 0 compared to the rest). We used trial 1 or 5 / experimental day 2 or 6 (depending on the procedure) as a baseline measurement, when mice were still naive to the following procedure (which was first experienced on that day). The latency of trial 1 was compared to trial 4 (experimental day 5 or 8, depending on the procedure) to see whether there was an increase or a reduction of the latency to leave the tunnel. For millet, latency to leave the tunnel was reduced (not normally distributed, $S = 10$, $p = 0.03857$). This was not the case for the restrainer procedure (not normally distributed, $S = 8$, $p = 0.3877$). Comparing the latency to leave the tunnel of both treatment procedures in trial 4 (after 3 experiences of the procedure), no difference was found ($t = -0.90693$, $p = 0.3776$).

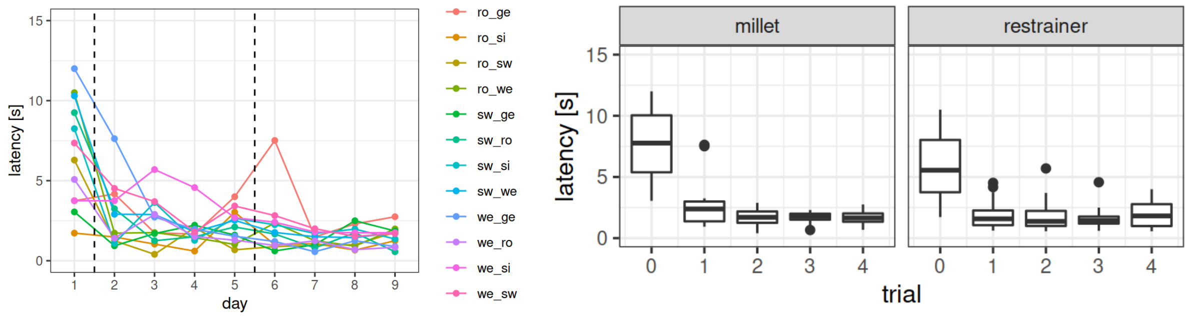

(a) (b)

**Figure 15:** Experiment 8.1: Latency to leave the tunnel (a) per mouse and day and (b) per conducted procedure. Twelve female C57BL/6J mice underwent a conditioning procedure, pairing one pattern (dots or stripes) with millet and the other floor pattern with restrainer. On day 1 (trial 0, habituation), mice spent 15 s in the presence of the pattern without a following procedure. Half of mice started with one pattern and half with the other. The first dotted line in **(a)** represents the beginning of the conditioning: Starting on day 2 (trial 1), after 15 s in presence of the pattern, the procedure followed for 1 min still in presence of the pattern). The second dotted line in **(a)** marks the procedure / pattern switch on day 5 (trial 1 other procedure). Note that in **(b)** trial 0 only contains 6 mice per procedure (habituation before conditioning without repetition after procedure / pattern switch), while all other trials contain 12. Latency was measured as the amount of time between positioning of the tunnel above the setup floor and the mouse leaving the tunnel with all four paws.
